# Ontogeny Dictates Oncogenic Potential, Lineage Hierarchy, and Therapy Response in Pediatric Leukemia

**DOI:** 10.1101/2025.03.19.643917

**Authors:** Ke Wang, Shayan Saniei, Nikita Poddar, Subrina Autar, Saul Carcamo, Meghana Sreenath, Jack H Peplinski, Rhonda E Ries, Isabella G Martinez, Clifford Chao, Anna Huo-Chang Mei, Noshin Rahman, Levan Mekerishvili, Miguel Quijada-Álamo, Grace Freed, Mimi Zhang, Katherine Lachman, Zayna Diaz, Manuel M Gonzalez, Jing Zhang, Giang Pham, Dan Filipescu, Mirela Berisa, Tommaso Balestra, Julie A Reisz, Angelo D’Alessandro, Daniel J Puleston, Emily Bernstein, Jerry E Chipuk, Mark Wunderlich, Sarah K Tasian, Bridget K Marcellino, Ian A Glass, BDRL, Christopher M Sturgeon, Dan A Landau, Zhihong Chen, Eirini P Papapetrou, Franco Izzo, Soheil Meshinchi, Dan Hasson, Elvin Wagenblast

**Affiliations:** Department of Oncological Sciences, Icahn School of Medicine at Mount Sinai, New York, NY, USA; Tisch Cancer Institute, Icahn School of Medicine at Mount Sinai, New York, NY, USA; Black Family Stem Cell Institute, Icahn School of Medicine at Mount Sinai, New York, NY, USA; Center for Advancement of Blood Cancer Therapies, Icahn School of Medicine at Mount Sinai, New York, NY, USA; Tisch Cancer Institute Bioinformatics for Next Generation Sequencing (BiNGS) core, Icahn School of Medicine at Mount Sinai, New York, NY, 10029, USA; Clinical Research Division, Fred Hutchinson Cancer Research Center, Seattle, WA, USA; Division of Hematology and Medical Oncology, Department of Medicine and Meyer Cancer Center, Weill Cornell Medicine, New York, NY, USA; Sandra and Edward Meyer Cancer Center, Weill Cornell Medicine, New York, NY, USA; Physiology, Biophysics and Systems Biology Graduate Program, Weill Cornell Medicine, New York, NY, USA; Department of Immunology and Immunotherapy, Icahn School of Medicine at Mount Sinai, New York, NY, USA; Department of Dermatology, Icahn School of Medicine at Mount Sinai, New York, NY, USA; Division of Oncology and Center for Childhood Cancer Research, Children’s Hospital of Philadelphia, Philadelphia, PA, USA; Department of Biochemistry and Molecular Genetics, University of Colorado Anschutz Medical Campus, Aurora, CO, USA; The Diabetes, Obesity, and Metabolism Institute, Icahn School of Medicine at Mount Sinai, New York, NY, USA; Division of Experimental Hematology and Cancer Biology, Cincinnati Children’s Hospital Medical Center, Cincinnati, OH, USA; Department of Pediatrics and Abramson Cancer Center, University of Pennsylvania Perelman School of Medicine; Philadelphia, PA, USA; Division of Hematology and Medical Oncology, Icahn School of Medicine at Mount Sinai, New York, NY, USA; Department of Pediatrics, University of Washington, WA, USA; Department of Cell, Developmental, and Regenerative Biology, Icahn School of Medicine at Mount Sinai, New York, NY, USA; New York Genome Center, New York, NY, USA; Department of Medicine, Icahn School of Medicine at Mount Sinai, New York, NY, USA; Department of Pediatrics, Division of Pediatric Hematology-Oncology, Icahn School of Medicine at Mount Sinai, New York, NY, USA; Mindich Child Health & Development Institute, Icahn School of Medicine at Mount Sinai, New York, NY, USA

**Keywords:** Acute myeloid leukemia, CRISPR/Cas9 modeling, fetal hematopoiesis, hematopoietic stem cells, leukemia stem cells, NUP98::NSD1, ontogeny, pediatric cancer, quiescence, therapy resistance

## Abstract

Accumulating evidence links pediatric cancers to prenatal transformation events, yet the influence of the developmental stage on oncogenesis remains elusive. We investigated how hematopoietic stem cell developmental stages affect leukemic transformation, disease progression, and therapy response using a novel, humanized model of NUP98::NSD1-driven pediatric acute myeloid leukemia, that is particularly aggressive with WT1 co-mutations. Fetal-derived hematopoietic stem cells readily transform into leukemia, and *WT1* mutations further enhance stemness and alter lineage hierarchy. In contrast, stem cells from later developmental stages become progressively resistant to transformation. Single-cell analyses revealed that fetal-origin leukemia stem cells exhibit greater quiescence and reliance on oxidative phosphorylation than their postnatal counterparts. These differences drive distinct therapeutic responses, despite identical oncogenic mutations. In patients, onco-fetal transcriptional programs correlate with worse outcomes. By targeting key vulnerabilities of fetal-origin leukemia cells, we identified combination therapies that significantly reduce aggressiveness, highlighting the critical role of ontogeny in pediatric cancer treatment.

## Introduction

Pediatric cancers arise with relative frequency despite minimal exposure to mutagens^1^, presenting a perplexing challenge in oncology. Unlike adult cancers, which typically develop through gradual accumulation of somatic mutations over decades^2^, many childhood cancers originate from genetic disruptions that occur during fetal or early postnatal development^3^. These early-life mutations frequently drive aggressive malignancies, such as neuroblastoma^4^ and leukemia^5^. Challenges associated with our inability to reliably model human cancer development across life stages have hindered progress in understanding the role of developmental context on cancer initiation and elucidating the mechanisms by which ontogeny shapes cancer biology.

Leukemia is the most common form of pediatric cancer. Acute myeloid leukemia (AML) is characterized by high relapse rates, therapy resistance, and suboptimal long-term survival.^6–9^ Pediatric AML is genetically heterogeneous, and chromosomal translocations such as *NUP98* rearrangements are key drivers of leukemogenesis.^7^ In *NUP98*-rearranged AML, the nuclear pore complex member NUP98 fuses with various partners such as *NSD1* and *KDM5A*.^10–14^ This AML subtype is characterized by specific features, including aberrant overexpression of HOXA/B cluster genes that drive self-renewal and block differentiation.^14^

The NUP98::NSD1 AML rearrangement frequently includes additional genetic aberrations, such as *WT1* loss-of-function mutations strongly associated with relapse and dismal outcomes^15,16^ and reported to be among the most common (*i.e.,* approximately 25% of cases) genetic alterations in relapsed pediatric AML.^16^ Patients harboring co-occurring *WT1* mutations are resistant to induction chemotherapy and have a 4-year event-free survival rate of less than 10%.^17,18^ These clinical observations underscore the importance of elucidating the molecular mechanisms involved in therapy resistance and prognosis worsening driven by the developmental context and co-occurring *WT1* mutations.

In this study, we systematically investigated how ontogeny shapes the oncogenic program of NUP98::NSD1-driven AML, which predominantly occurs in pediatric, adolescent, and young adult patients and is nearly absent in older adults.^14^ We applied CRISPR/Cas9-mediated genome engineering to developmentally defined human primary hematopoietic stem and progenitor cells (HSPCs) and precisely modeled leukemia initiation and progression across development (*i.e.,* from human fetal stages to postnatal adult contexts). Our studies revealed that the temporal origin of leukemia profoundly influences cancer biology, therapeutic vulnerabilities, and clinical outcomes, even in cases with identical driver mutations. These results suggest that leveraging ontogeny-specific features can enhance risk stratification and facilitate the development of tailored therapies for pediatric cancer patients.

## Results

### NUP98::NSD1 Fusion Induces Leukemia in a Developmental Stage-Specific Manner

We first investigated the impact of HSPC developmental stages on NUP98::NSD1-mediated initiation of leukemic transformation by comparing human HSPCs from fetal liver (FL), postnatal cord blood (CB), and pediatric and adult bone marrow (BM). We induced chromosomal translocations at the natural break sites of *NUP98* on chromosome 11 and *NSD1* on chromosome 5 using CRISPR/Cas9-mediated genome engineering and generated in-frame NUP98::NSD1 fusions in CD34+ HSPCs at each developmental stage (**Figure 1A**). A fraction of the cells (*i.e.,* 2-5%) harbored the NUP98::NSD1 fusion at the genomic DNA level; endogenous fusion expression was confirmed at the RNA level (**Supplementary Figure 1A-C)**. To model WT1 loss-of-function (WT1ko), we targeted exon 2 of *WT1* using a guide RNA (gRNA) and achieved >90% editing efficiency (**Supplementary Figure 1D**), which resulted in near complete loss of WT1 protein expression (**Supplementary Figure 1E**).

**Figure 1.**
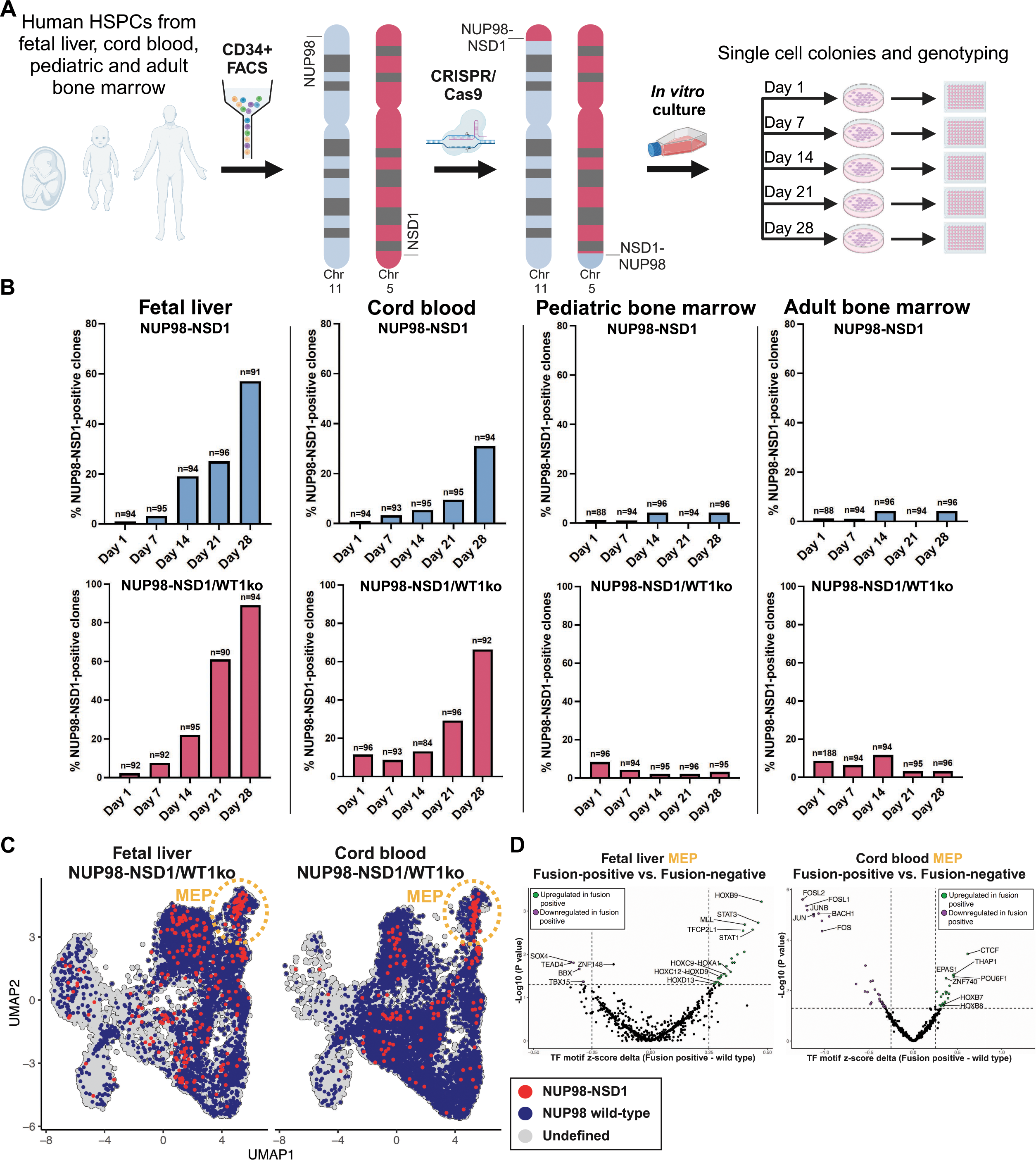
Developmental Stage-Specific Susceptibility of HSPCs to NUP98::NSD1-Driven Leukemogenesis. **(A)** Overview of the experimental approach to assess the developmental specificity of NUP98::NSD1 driven AML *in vitro* using CRISPR/Cas9 to edit HSPCs from different developmental sources, culturing them for 28 days, and plating a subset on methylcellulose every 7 days. **(B)** Percentages of NUP98::NSD1-positive colonies across developmental stages over four weeks, as shown in (A). The number of colonies (n) is indicated on top of each bar. **C)** UMAP based on GoT-ChA of FL- and CB-derived HSPCs edited with NUP98::NSD1 and WT1ko, cultured *in vitro* for 4 weeks. Genotypes are overlaid on UMAP clusters. Red dots represent NUP98::NSD1 fusion-positive cells, blue dots represent NUP98 wild-type cells, and gray dots represent cells with undefined genotypes. The orange-dotted lines represent the most primitive MEP cluster. **(D)** Volcano plots of differential transcription factor motifs between wild-type and NUP98::NSD1 fusion-positive cells within the MEP population in FL and CB.

Next, we generated four groups of human CD34+ HSPCs using editing approaches. The groups included control (targeting the olfactory receptor OR2W5), NUP98::NSD1, WT1 knockout (WT1ko), and NUP98::NSD1/WT1ko (**Figure 1A**). Cells were cultured for four weeks, and subsets were plated weekly on methylcellulose colony-forming media to monitor the enrichment of self-renewing colony-forming cells over time. Single-cell-derived colonies were subsequently genotyped for the NUP98::NSD1 fusion. Strikingly, NUP98::NSD1-positive clones from FL-derived HSPCs exhibited a strong selective advantage, increasing from a 2% initial abundance to 57% in NUP98::NSD1 and 89% in NUP98::NSD1/WT1ko by week 4 (**Figure 1B**). CB-derived HSPCs displayed similar positive selection trends, albeit with lower enrichment. In contrast, NUP98::NSD1-positive clones derived from pediatric and adult BM showed no evidence of positive selection. Colony genotyping of control and WT1ko groups confirmed consistently high editing efficiency (**Supplementary Figure 1F**).

Next, we investigated the mechanisms underlying the clonal advantage conferred by NUP98::NSD1 in FL and CB HSPCs. We employed single-cell genotyping of targeted loci by adapting GoT-ChA^19^ to allow simultaneous determination of chromatin accessibility profile and genotype at the NUP98::NSD1 break site, thereby distinguishing fusion-positive from wild-type cells in the same sample (**Supplementary Figure 1G**). To validate this approach, we first performed GoT-ChA on a 1:1 mixture of NUP98::NSD1-positive cells and wild-type FL CD34+ HSPCs. As expected, mutant and wild-type cells clustered distinctly, reflecting unique chromatin states (**Supplementary Figure 1H).** Mutation overlay confirmed the accurate classification of fusion-positive cells with a low false-positive rate (**Supplementary Figure 1I)**. We then applied GoT-ChA to *in vitro*-cultured FL and CB HSPCs four weeks post-NUP98::NSD1/WT1ko editing. Cell states, captured *via* chromatin accessibility, were predominantly aligned along the myeloid-erythroid differentiation pathway, consistent with *in vitro-*priming taking place in HSPCs (**Supplementary Figure 1J**). Genotyping overlay on cluster assignments revealed an expansion of fusion-positive cells in the most primitive megakaryocyte-erythroid progenitors (MEPs, **Figure 1C**).

To elucidate the mechanisms involved in FL and CB HSPC clonal advantage induced by NUP98::NSD1/WT1ko editing, we conducted transcription factor (TF) motif enrichment analyses comparing wild-type and NUP98::NSD1 fusion-positive cells in the MEP cluster (**Figure 1D and Supplementary Table S1**). FL fusion-positive cells displayed increased enrichment with several HOX cluster genes, *MLL,* and *STAT1/3*.^14^ CB fusion-positive cells exhibited higher enrichment for other *HOX* cluster genes, *CTCF,* and reduced enrichment for AP-1 inflammatory mediators compared to wild-type cells. Interestingly, FL- and CB-edited HSPCs showed largely distinct TF enrichment patterns, suggesting that the fusion protein drives individual expression programs to initiate leukemia in a developmental stage-specific fashion.

### NUP98::NSD1 Fusion and WT1 Knockout Initiate AML with Enhanced Leukemic Propagation and Stem Cell Characteristics

We next explored the impact of developmental context on the ability of NUP98::NSD1, with or without WT1 loss-of-function, to drive leukemogenesis *in vivo*. We established a humanized xenotransplantation mouse model using CRISPR/Cas9-engineered HSPCs at distinct developmental stages. First, we utilized human FL CD34+ HSPCs and transplanted four edited cell groups – *i.e.*, control, NUP98::NSD1, WT1ko, and NUP98::NSD1/WT1ko **–** followed by engraftment in immunodeficient NOD scid gamma (NSG) mice for 16 weeks **(Figure 2A)**. The bone marrow of FL-xenografted mice NUP98::NSD1 and NUP98::NSD1/WT1ko groups showed a significant increase in CD33+ myeloid cells and an accumulation of CD34+CD117+ stem and progenitor populations compared to controls (**Figure 2B** and **Supplementary Figures 2A and B**). In contrast, there were no differences in engraftment levels or lineage differentiation when comparing control and WT1ko cells. Since genetically edited NUP98::NSD1-positive and - negative HSPCs co-propagated in individual mice, we assessed leukemia stem cell (LSC) frequency and self-renewal capacity through secondary xenotransplantation, using limiting dilution assays (LDA) in NSG mice (**Figure 2C**). NUP98::NSD1-expressing FL cells exhibited markedly enhanced self-renewal compared to control-edited cells (**Figure 2D**). Control FL xenografts displayed a low initiating-cell frequency of 1 in 830,000 cells, while NUP98::NSD1 and NUP98::NSD1/WT1ko FL xenografts displayed >50-fold higher initiating-cell frequencies of <1 in 9,300 and 15,600 cells, respectively. Morphological cytospin analyses confirmed the presence of leukemic blasts and, therefore, the leukemic phenotype of these mice (**Figure 2E**). Secondary FL NUP98::NSD1/WT1ko xenograft mice exhibited similar overall engraftment levels compared with their NUP98::NSD1 counterparts but harbored significantly higher proportions of CD34+CD117+ stem and progenitor cells, lower percentages of CD33+ myeloid cells, and a relative increase in CD19+ lymphoid cells (**Figure 2F, Supplementary Figure 2C and D**).

**Figure 2.**
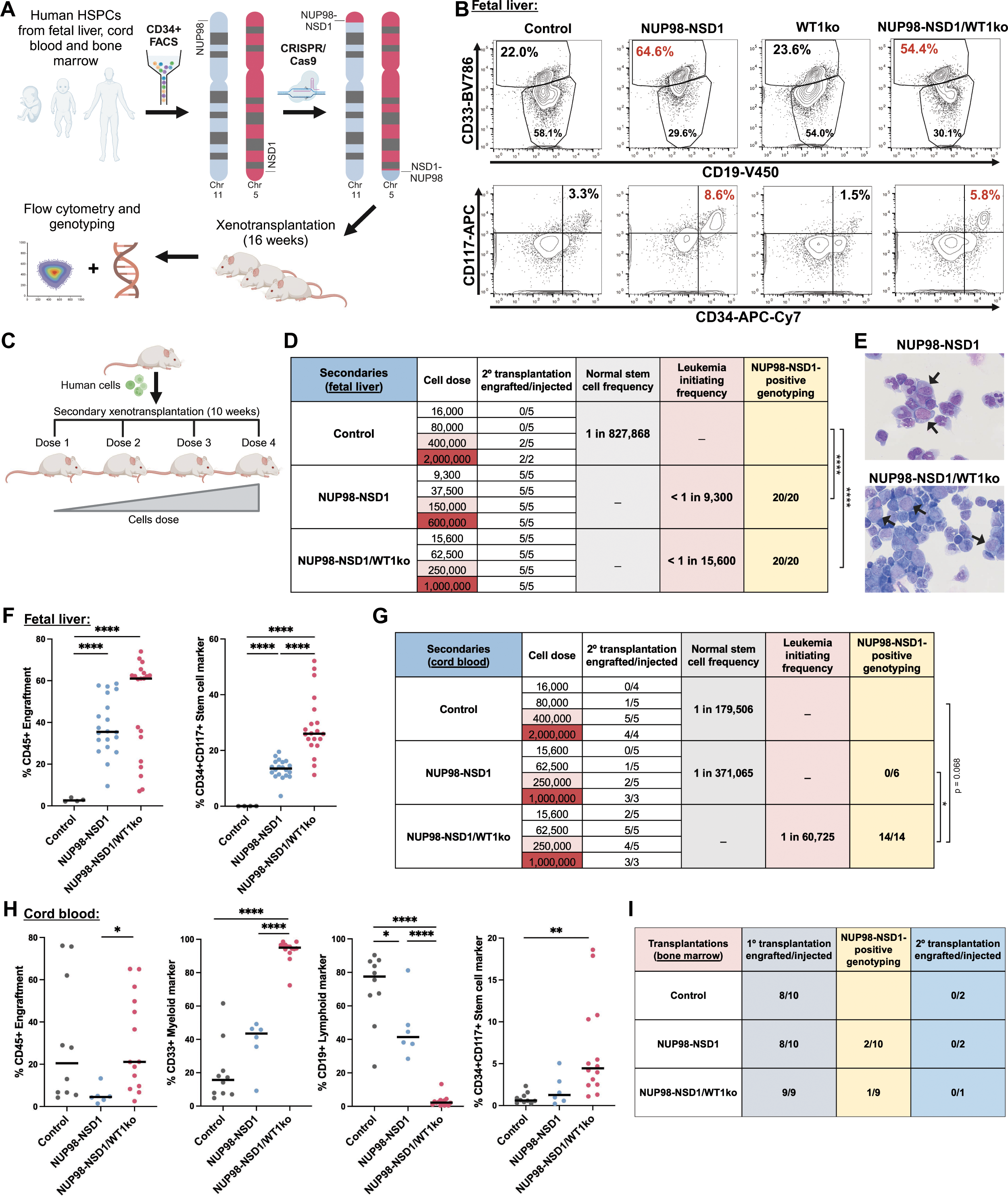
Developmental Context Determines Leukemogenic Potential and Stemness in NUP98::NSD1 AML. **(A)** Schematic of the experimental approach for endogenous induction of NUP98::NSD1 rearrangement in human HSPCs *via* CRISPR/Cas9, followed by intrafemoral injection into NSG mice and 16 weeks of engraftment. **(B)** Representative flow plots showing CD33 myeloid cells, CD19 lymphoid cells, and CD34+CD117+ stem cells in control, NUP98::NSD1, WT1ko, and NUP98-NSD/WT1ko-edited primary xenografts. n = 5-16 mice per condition from three independent cohorts. Only mice with confirmed CRISPR/Cas9 edits and >90% efficiency in control and WT1ko groups and NUP98::NSD1 fusion-positive status were included in downstream analyses. **(C)** Schematic of our strategy to establish secondary xenografts from control, NUP98::NSD1, WT1ko, and NUP98-NSD/WT1ko-edited primary xenografts. Cells were injected into NSG mice using a limiting dilution assay (LDA) approach and were allowed to engraft for 10 weeks. **(D)** Normal stem cell frequency for control xenografts and leukemia-initiating stem cell frequency for FL-derived HSPCs based on the LDA assay described in (C). ****p < 0.0001 using Pearson’s chi-squared test, n = 17-20 mice per condition (2-5 mice per dosage). **(E)** Morphological analysis of human cells in FL NUP98::NSD1 and NUP98::NSD1/WT1ko secondary xenografts (arrows depict blast cells, 63x magnification). **(F)** Engraftment of secondary FL xenografts in NSG mice, assessed by human CD45+ cells in injected bone marrow; stem and progenitor cell populations based on CD34+CD117+ double-positive cells. The line represents the mean value. ****p < 0.0001, unpaired t-test; n = 4-20 mice per condition. **(G)** Normal stem cell frequency in control xenografts and leukemia-initiating stem cell frequency in CB-derived NUP98::NSD1 generated models were determined using the LDA approach described in (C). *p < 0.05, analyzed by Pearson’s chi-squared test, n = 18 mice per condition (4-5 mice per dosage). **(H)** Engraftment and percentage of CD33 myeloid cells, CD19 lymphoid cells, and CD34+CD117+ stem cells in secondary CB xenografts in NSG mice; the line represents the mean value. *p < 0.05, **p < 0.01, and ****p < 0.0001, unpaired t-test; n = 6-14 mice per condition. **(I)** Number of mice transplanted with bone marrow-derived edited HSPCs, number of NUP98::NSD1 fusion-positive mice, and number of mice with engraftment in secondary transplanted xenografts. n = 8 - 10 mice per condition from 2 independent cohorts.

To characterize clonal architecture, we placed bone marrow cells from FL secondary xenografts on methylcellulose colony-forming media. Genotyping of individual single-cell-derived colonies revealed near 100% positivity for the NUP98::NSD1 fusion while maintaining one wild-type *NUP98* allele (**Supplementary Figure 2E**). Sanger sequencing of bulk cells confirmed the presence of in-frame fusions (**Supplementary Figure 2F**). *In vitro* culture of FL NUP98::NSD1-harboring leukemic xenografts confirmed their capacity to retain stem-like and myeloid features. Specifically, NUP98::NSD1/WT1ko cells contained a higher percentage of CD34+ cells and lower percentages of mature myeloid markers than NUP98::NSD1 cells, consistently with *in vivo* findings (**Supplementary Figure 2G**). Karyotyping revealed no additional chromosomal abnormalities (**Supplementary Figure 2H**). Lastly, we observed that both NUP98::NSD1 and NUP98::NSD1/WT1ko FL-derived leukemia cells displayed biomolecular condensates (**Supplementary Figure 2I**), a feature of NUP98 fusion proteins caused by their intrinsically disordered regions.^20,21^ In summary, using FL-derived HSPCs, we generated a tractable humanized model of NUP98::NSD1-rearranged AML incorporating WT1 loss-of-function as a cooperating mutation.

We next sought to assess the developmental specificity of NUP98::NSD1-driven AML *in vivo*. Using a modeling approach like that described for FL, we transplanted control, NUP98::NSD1, NUP98::NSD1/WT1ko-edited CB, and adult BM CD34+ HSPCs into NSG mice. After 16 weeks, primary xenotransplantations with CB NUP98::NSD1-edited HSPCs yielded fusion-positive human xenografts (**Supplementary Figure 2J**). To evaluate self-renewal potential, we transplanted human cells from control and NUP98::NSD1-positive primary CB xenografts into secondary NSG mice using LDA (**Figure 2G**). Secondary xenografts demonstrated significantly enhanced stem cell frequency in NUP98::NSD1/WT1ko-edited CB cells compared to NUP98::NSD1 and control. Importantly, all secondary xenografts derived from NUP98::NSD1/WT1ko-edited CB cells were fusion-positive, whereas secondary xenografts from NUP98::NSD1-edited CB cells showed no detectable fusion, suggesting no positive selection advantage in the latter. Immunophenotypic analyses revealed significant enrichment of CD33+ myeloid cells and CD34+CD117+ stem and progenitor populations in CB NUP98::NSD1/WT1ko xenografts compared to NUP98::NSD1 and control (**Figure 2H and Supplementary Figure 2K**). Importantly, morphological analyses confirmed the presence of leukemic blasts only in CB NUP98::NSD1/WT1ko xenografts (**Supplementary Figure 2L**). These results highlight that NUP98::NSD1 requires cooperating mutations, such as WT1 loss-of-function, to induce leukemogenesis in developmentally advanced HSPCs, such as cord blood cells. In contrast, NUP98::NSD1 alone appears sufficient for AML initiation during early developmental stages, as modeled by FL-derived HSPCs.

As expected, NUP98::NSD1-edited BM-derived HSPCs failed to produce fusion-positive primary xenografts (**Figure 2I**). Two primary xenografts from NUP98::NSD1- and one from NUP98::NSD1/WT1ko-edited BM cells contained detectable fusion-positive cells, but secondary transplantation yielded no human engraftment. These findings point to the inability of NUP98::NSD1, even with WT1 loss-of-function, to transform BM-derived HSPCs. Our results underscore the developmental specificity of NUP98::NSD1-driven AML, which is restricted to fetal and early postnatal windows. The fusion progressively loses its ability to transform HSPCs as they age, indicating a critical dependency of leukemia initiation on the stage of cellular development.

### Single-Cell Analysis Reveals Distinct Transcriptional and Epigenetic Landscapes in NUP98::NSD1-Driven AML

Next, we tested the hypothesis that the transcriptional and epigenetic landscapes contribute to the developmental specificity and aggressive nature of NUP98::NSD1-driven leukemia. We performed single-cell RNA and ATAC sequencing (scRNA-seq and scATAC-seq) on human CD45+ cells harvested from our FL xenografts. Given that serially transplanted mice harbored only NUP98::NSD1-positive cells due to the exhaustion of wild-type HSPCs (**Figure 2D**), we analyzed secondary xenografts for NUP98::NSD1 and NUP98::NSD1/WT1ko models and primary xenografts for control and WT1ko models. Using cluster markers and a single-cell reference dataset^22^, we annotated the hematopoietic lineages in our merged scRNA-seq atlas (**Supplementary Figure 3A**). We analyzed high-quality cells (n=64,156), revealing 17 transcriptionally distinct clusters spanning hematopoietic stem cells/multipotent progenitor cells (HSC/MPP) to mature myeloid and lymphoid lineages (**Figure 3A**). Control and WT1ko xenografts were predominantly composed of lymphoid cells with limited myeloid and progenitor populations, consistent with wild-type lymphoid-biased hematopoiesis in NSG mice (**Figure 3B**). In contrast, NUP98::NSD1 and NUP98::NSD1/WT1ko FL xenografts displayed significant enrichment of HSPCs and myeloid lineages (**Figure 3B and C**). Notably, NUP98::NSD1/WT1ko xenografts showed a substantial expansion of primitive bipotent stem-like cells and myeloid-primed progenitors, including lympho-myeloid primed progenitors (LMPPs), early granulocyte-monocyte progenitors (GMPs), and immature and committed B-lymphoid progenitors, compared to NUP98::NSD1 xenografts. To validate our findings using an independent approach, we analyzed NUP98::NSD1 AML patient-derived xenografts (PDXs) with and without *WT1* co-mutations. PDX model cells with *WT1* mutations transcriptionally aligned with primitive LMPP/Early GMPs and GMPs, while PDX cells without *WT1* mutations mapped to later-stage progenitors, mirroring our *in vivo* models (**Figure 3D and Supplementary Figure 3B**).

**Figure 3.**
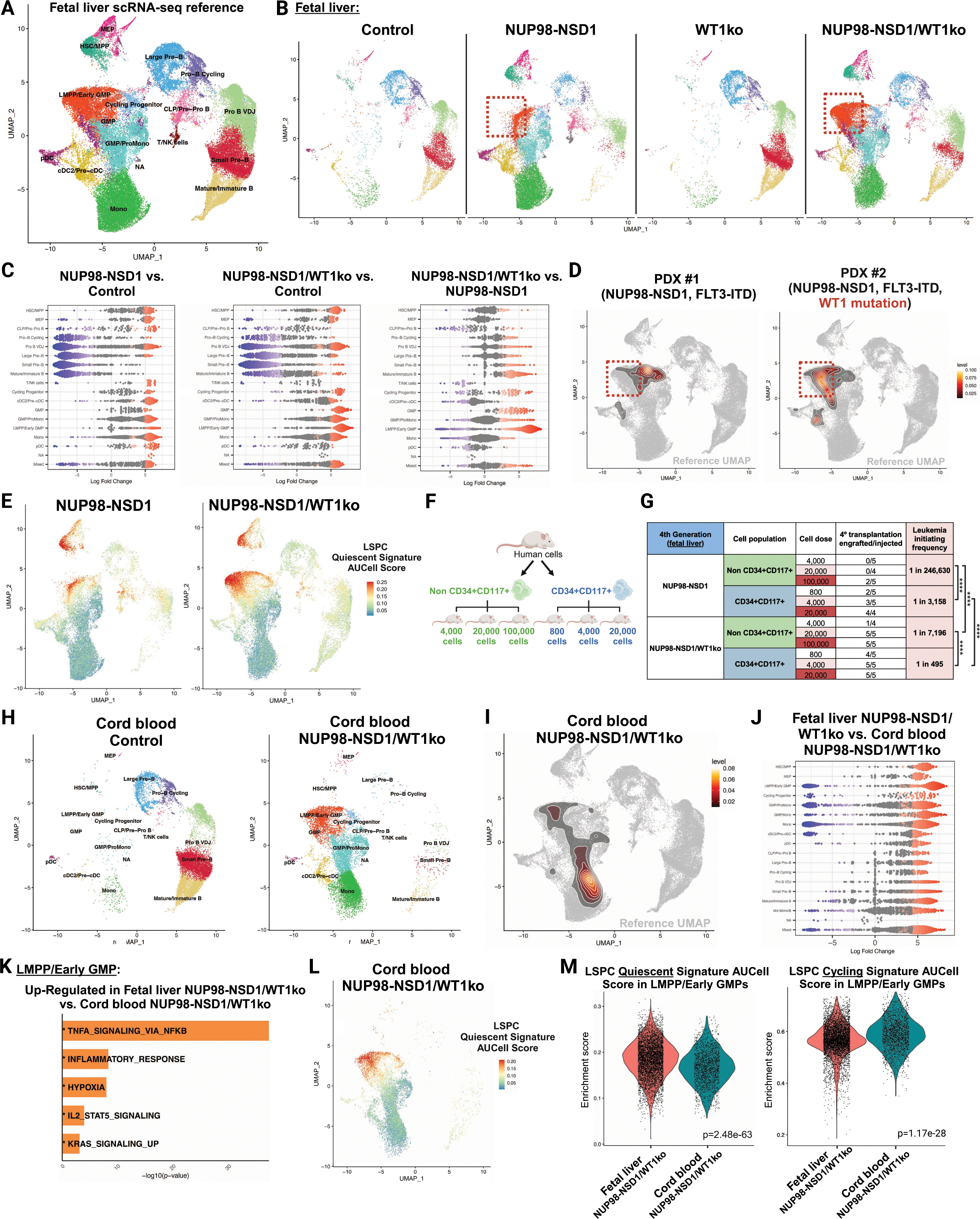
Single-Cell Profiling Reveals Developmentally Regulated Transcriptional and Epigenetic Programs in NUP98::NSD1-Driven AML. **(A)** UMAP of merged scRNA-seq reference, including normal and leukemic xenografts, categorized into 17 clusters. Control, n = 2 biological replicates; NUP98::NSD1, n = 4; WT1ko, n = 2, and NUP98::NSD1/WT1ko, n = 3. **(B)** Contribution of each group to the merged scRNA-seq UMAP. The red dotted box highlights LMPP/Early GMPs in NUP98::NSD1 and NUP98::NSD1/WT1ko condition. **(C)** Beeswarm plot comparing log-fold changes in nearest neighbor cells from different cell type clusters computed with Milo between NUP98::NSD1 to control, NUP98::NSD1/WT1ko to control, and NUP98::NSD1/WT1ko to NUP98::NSD1. Significant neighborhoods identified at FDR < 0.01 are indicated in color. **(D)** Mapping of scRNA-seq for PDX #1 (NUP98::NSD1, FLT3-ITD) and PDX #2 (NUP98::NSD1, FLT3-ITD, WT1 mutation) onto the merged scRNA-seq reference, displaying relative cell abundance density. **(E)** UMAP of FL NUP98::NSD1 and NUP98::NSD1/WT1ko samples, by Area Under the Curve (AUC) score for the LSPC quiescent gene signature. Higher AUC scores (red) indicate stronger enrichment of the quiescent signature, while lower scores (blue) represent weaker enrichment. **(F)** Schematic experimental overview of the LDA experiment with CD34+CD117+ enriched LSCs and non-CD34+CD117+ cells sorted from FL NUP98::NSD1 and NUP98::NSD1/WT1ko xenografts. **(G)** Leukemia initiation stem cell frequency for CD34+CD117+ enriched LSCs and non-CD34+CD117+ cells based on the LDA approach described in (F). ****p < 0.0001 using Pearson’s chi-squared test, n = 14-15 mice per condition (4-5 mice per dosage). **(H)** UMAP of secondary xenografts from CB-derived HSPCs including control and NUP98::NSD1/WT1ko. n = 2 biological replicates per condition. **(I)** Mapping scRNA-seq for CB NUP98::NSD1/WT1ko onto the merged FL-derived scRNA-seq reference, contoured with relative abundance density of cells in each sample. **(J)** Besswarm plot showing the log-fold change comparing FL NUP98::NSD1/WT1ko xenografts and CB NUP98::NSD1/WT1ko xenografts for groups of nearest neighbor cells from different cell type clusters computed with Milo. Significant neighborhoods identified at an FDR < 0.01 are indicated in color. n = 3 for FL NUP98::NSD1/WT1ko and n = 2 for CB NUP98::NSD1/WT1ko. **(K)** Bar plot showing significantly upregulated hallmark pathways in FL NUP98::NSD1/WT1ko cells relative to CB NUP98::NSD1/WT1ko in the LMPP/Early GMP cluster. Pathways in bold and with an asterisk (*) are statistically significant based on the cutoff p-value < 0.05. The x-axis represents -log10(p-value), with higher values indicating greater statistical significance. **(L)** UMAP of CB NUP98::NSD1/WT1ko with cells colored by their AUC score for the LSPC quiescent signature using AUCell. **(M)** Violin plots showing AUCell enrichment scores for LSPC quiescent (left) and LSPC cycling (right) in LMPP/Early GMPs from FL NUP98::NSD1/WT1ko and CB NUP98::NSD1/WT1ko. Wilcoxon rank-sum test.

We next characterized the chromatin accessibility landscape using scATAC-seq; we analyzed 48,662 high-quality cells across NUP98::NSD1, NUP98::NSD1/WT1ko, and control FL xenografts (**Supplementary Figure 3C-E**). Consistent with the scRNA-seq data, leukemic xenografts were enriched for myeloid and progenitor populations, with NUP98::NSD1/WT1ko xenografts showing LMPP/Early GMP expansion. Cell type-specific motif enrichment analyses^23^ revealed active TF motifs such as *CTCF*, *MEIS2*, and *PBX2* in HSC/MPPs, while CEBP family TFs displayed heightened activity along the myeloid differentiation trajectory **(Supplementary Table S2)**. *RUNX1* and *SPI1* gained significant activity, specifically in LMPP/Early GMPs, while inflammation-associated TFs such as *JUN*, *FOS,* and *SMAD* were enriched across the stem and myeloid compartments (**Supplementary Figure 3F**), aligning with general molecular programs in AML.^24^

Previous work showed that quiescent leukemia stem and progenitor cells (LSPCs) correlate with resistance to therapy and poor prognosis^25^. Further, we found significant enrichment of the quiescent LSPC signature in LMPP/Early GMP populations, particularly in NUP98::NSD1/WT1ko xenografts (**Figure 3E**). Cell cycle scoring confirmed that these cells were primarily in the G1 phase (**Supplementary Figure 3F**). RNA velocity analyses^26^ indicated differentiation trajectories originating from cycling progenitors toward LMPP/Early GMPs, myeloid progenitors, and lymphoid cells, suggesting a bipotential differentiation axis at the progenitor stage (**Supplementary Figure 3G**). Diffusion pseudotime analyses^27^ further positioned cycling progenitors as upstream intermediates, preceding LMPP/Early GMPs (**Supplementary Figure 3H and I**). These findings suggest a hierarchical relationship whereby LMPP/Early GMPs and cycling progenitors are closely associated at the top of a myeloid/lymphoid bipotent differentiation axis, where cycling progenitors can either establish highly quiescent LMPP/Early GMPs and lymphoid populations (as in NUP98::NSD1/WT1ko) or skew towards committed myeloid progenitors (as in NUP98::NSD1).

These findings led us to hypothesize that *WT1* loss-of-function mutations in NUP98::NSD1-driven AML enhance LSC self-renewal by promoting the expansion of LMPP/Early GMPs. We isolated LMPP/Early GMPs and experimentally validated their role as the primary reservoir of the most potent LSCs. CD34 and CD117 expressions are aligned with the transcriptionally defined LMPP/Early GMP compartment (**Supplementary Figure 4A**). To functionally test this population, we sorted CD34+CD117+ and non-CD34+CD117+ fractions from FL NUP98::NSD1 and NUP98::NSD1/WT1ko xenografts and transplanted them into NSG mice using LDA (**Figure 3F**). CD34+CD117+ cells demonstrated a significantly higher LSC frequency than non-CD34+CD117+ cells, confirming that the most potent LSCs reside in this compartment (**Figure 3G**). Notably, NUP98::NSD1/WT1ko xenografts exhibited a markedly higher LSC frequency than NUP98::NSD1 alone, underscoring the cooperative effect of WT1 loss in enhancing self-renewal potential. To further assess the lineage plasticity of NUP98::NSD1/WT1ko LSCs, or their ability to generate both myeloid and lymphoid lineages, we sorted CD19-negative cells (myeloid-enriched) from NUP98::NSD1 and NUP98::NSD1/WT1ko secondary FL xenografts and transplanted them into NSG mice (**Supplementary Figure 4B**). Strikingly, CD19-negative NUP98::NSD1/WT1ko LSCs generated significantly larger xenografts containing a high percentage of CD19+ B-lymphoid cells, demonstrating their dual myeloid-lymphoid differentiation potential (**Supplementary Figure 4C**). In contrast, CD19-negative NUP98::NSD1 LSCs failed to produce a substantial CD19+ B-lymphoid population, reflecting their restricted lineage output. Our findings indicate that WT1 loss promotes a more stem-like, bipotent state in NUP98::NSD1-driven AML, resulting in a more shallow and primitive leukemia hierarchy enriched for LMPP/Early GMP-like LSCs. WT1 loss further enhances plasticity and self-renewal, expands the pool of LSCs, and confers leukemogenic potential to this progenitor population, likely accounting for the aggressive nature of NUP98::NSD1/WT1ko leukemia.

To further delineate the developmental specificity of NUP98::NSD1-driven AML, we performed scRNA-seq on NUP98::NSD1/WT1ko secondary xenografts derived from CB CD34+ HSPCs and compared them to control-edited CB primary xenografts. NUP98::NSD1 with retained wild-type *WT1* was excluded because it failed to induce leukemia in CB HSPCs *in vivo* (**Figure 2G**). Overlaying CB models onto the scRNA-seq reference UMAP highlighted significant concordance between control-edited CB and FL xenografts; both predominantly consisted of B lymphocytes (**Supplementary Figure 4D and E**). In CB NUP98::NSD1/WT1ko xenografts, we observed a lineage distribution generally resembling the FL NUP98::NSD1/WT1ko model but noted critical differences. While the LMPP/Early GMP compartment was comparable between CB and FL xenografts, B-lymphoid populations were more abundant in FL xenografts (**Figure 3H - J**). Immunophenotypic data supported this finding, showing diminished B-lymphoid output and primarily myeloid differentiation trajectories in secondary CB NUP98::NSD1/WT1ko xenografts (**Figure 2H**). Therefore, CB-originating LSCs exhibit lower lineage plasticity, with reduced ability to generate both myeloid and lymphoid lineages, compared to their FL counterparts.

Aberrant *HOXA/B* overexpression is a hallmark of NUP98-driven AML^14^. Thus, we evaluated the expression of 13 *HOXA/B* genes across our models (**Supplementary Figure 4F**). Control xenografts exhibited negligible *HOX* cluster expression limited to a small subset of HSC/MPP populations. In contrast, FL NUP98::NSD1 and NUP98::NSD1/WT1ko xenografts displayed high *HOX* gene expression across multiple cell types, consistent with endogenous NUP98::NSD1 fusion activation. Aberrant *HOX* expression was also observed in CB NUP98::NSD1/WT1ko cells, albeit at lower levels than FL leukemic xenografts.

Differential expression analyses comparing leukemic xenografts and their respective controls unveiled pathways enriched in LSCs; we focused our analyses on progenitor populations such as GMP/ProMono. CB and FL-derived LSCs shared enrichment in pathways such as hypoxia, NF-κB signaling, reactive oxygen species (ROS), and inflammation (**Supplementary Figure 4G**). However, comparing LMPP/Early GMP populations from FL versus CB NUP98::NSD1/WT1ko xenografts revealed significant transcriptional differences (**Supplementary Table S3)**. Notably, NF-κB signaling, inflammatory pathways, hypoxia, and *STAT5* signaling were markedly enriched in FL-originating LSCs compared to their CB counterparts (**Figure 3K**). Single-cell normalized LSPC quiescent signature enrichment confirmed that CB-originating LMPP/Early GMPs exhibited quiescent characteristics (**Figure 3L**). However, this quiescent enrichment was less pronounced in CB-originating LMPP/Early GMPs compared to FL counterparts, as they also exhibited greater enrichment in the LSPC cycling signature (**Figure 3M**). The higher quiescent state of FL-originating primitive LSCs is intriguing, given that wild-type FL HSPCs are typically associated with a higher proliferative capacity than postnatal CB-derived HSPCs. Our findings suggest that the developmental timing of NUP98::NSD1 fusion acquisition profoundly restructures AML lineage hierarchies, altering self-renewal properties and LSC transcriptional programs in a stage-specific manner.

### Transcriptional and Epigenetic Impacts of WT1 Loss-of-Function

To elucidate the role of WT1 loss-of-function in transcriptional rewiring, quiescence, and resistance to therapy in NUP98::NSD1-driven AML, we analyzed the LSC transcriptional and epigenetic landscapes. First, we explored the functional consequences of WT1ko *in vivo*. Previous studies have shown that *WT1* expression, which is low in normal HSPCs, is upregulated and critical in multiple adult AML subtypes.^28^ Loss-of-function *WT1* mutations are associated with poor outcomes, therapy resistance, and relapse of several AML subtypes, including the majority of NUP98-rearranged cases.^17^ Interestingly, our scATAC-seq data revealed that *WT1* motif enrichment is elevated in the LMPP/Early GMP population in NUP98::NSD1 compared to control, suggesting that WT1 TF activity increases following NUP98::NSD1 translocation (**Supplementary Figure 5A**). This observation may explain why WT1ko in NUP98-rearranged AMLs leads to downstream consequences such as the expansion of stem-like populations and enhanced quiescence. A comparison of NUP98::NSD1 versus NUP98::NSD1/WT1ko motif enrichment in LMPP/Early GMPs indicated that *WT1* exhibited decreased motif enrichment in NUP98::NSD1/WT1ko, confirming that our model accurately recapitulates WT1 loss-of-function phenotypes (**Supplementary Figure 5B**).

To further investigate the downstream consequences of WT1 loss-of-function in LSCs, we analyzed differential gene expression within the LMPP/Early GMP population by comparing FL NUP98::NSD1/WT1ko versus NUP98::NSD1. This analysis revealed significant transcriptional reprogramming resulting from WT1 loss-of-function (**Figure 4A**). Interestingly, while the AP-1 family of TF involved in inflammatory responses exhibited greater motif accessibility in NUP98::NSD1/WT1ko than in NUP98::NSD1 (**Supplementary Figure 5B**), inflammatory pathways, including TNF-α *via* NF-κB signaling, were downregulated with WT1ko (**Figure 4B**). Although both NUP98::NSD1 and NUP98::NSD1/WT1ko LSCs exhibited heightened inflammation compared to healthy hematopoiesis (**Supplementary Figure 4G**), NUP98::NSD1/WT1ko LSCs exhibited reduced inflammatory stress. This attenuation may prevent cellular exhaustion and facilitate the maintenance of a more quiescent and unprimed state, potentially distinguishing NUP98::NSD1/WT1ko from NUP98::NSD1 in their response to inflammatory cues.

**Figure 4.**
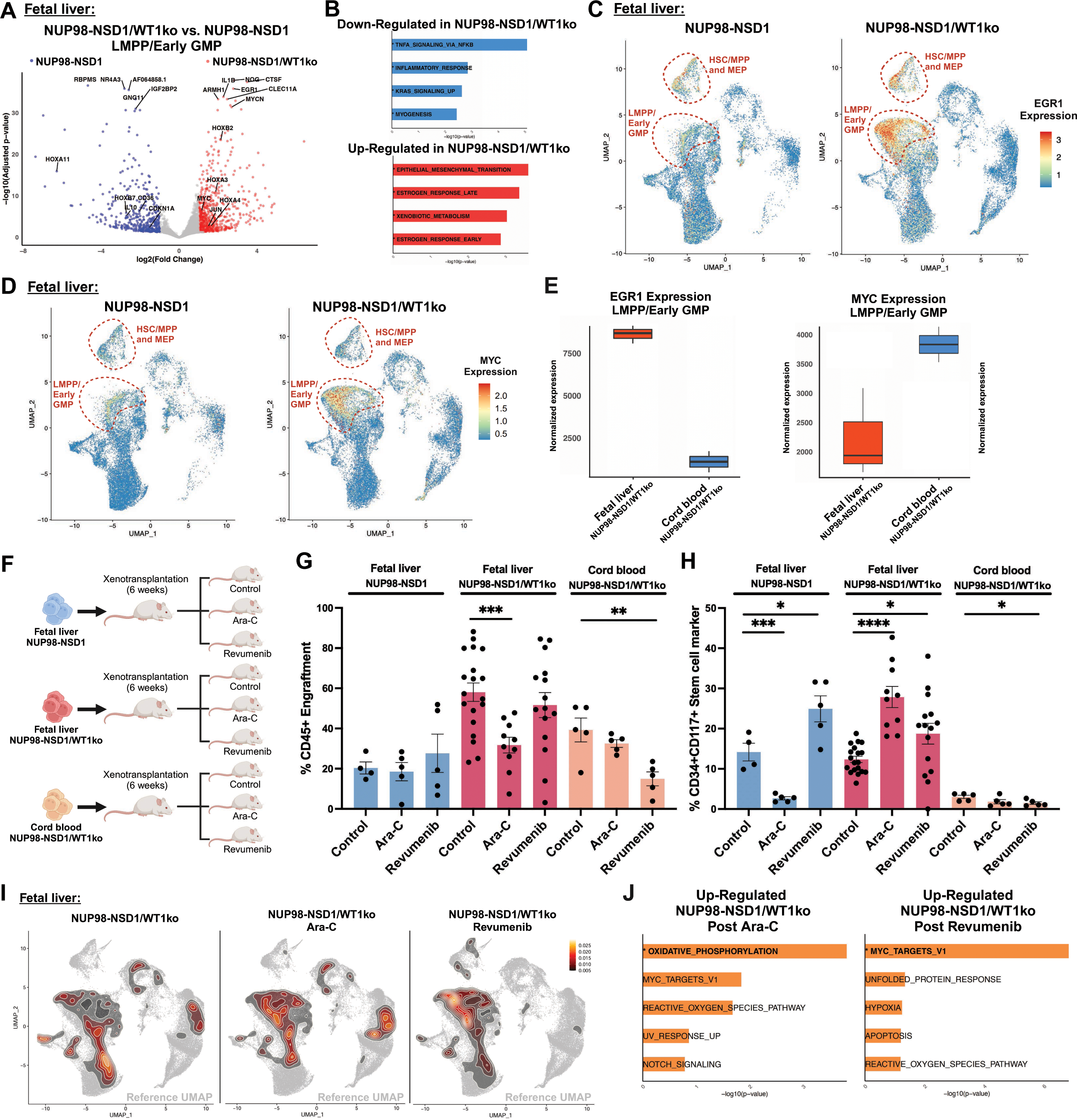
WT1 Loss and Ontogeny Shape Differential Therapy Responses in NUP98::NSD1-Driven AML. **(A)** Volcano plot of differentially expressed genes in LMPP/Early GMPs between FL NUP98::NSD1/WT1ko and NUP98::NSD1. Genes with significant adjusted p-values < 0.05 and log2FC > 1 are colored; representative upregulated and downregulated genes are indicated. **(B)** Bar plots showing Over Representation Analysis (ORA) results of differentially expressed LMPP/Early GMPs genes between NUP98::NSD1/WT1ko and NUP98::NSD1. Significant hallmark pathways are highlighted in bold and with an asterisk (*) based on p-value < 0.05. **(C)** UMAP of *EGR1* expression in FL NUP98::NSD1 and NUP98::NSD1/WT1ko xenografts. **(D)** UMAP of *MYC* expression in FL NUP98::NSD1 and NUP98::NSD1/WT1ko xenografts. **(E)** Boxplots showing normalized expression of *EGR1* (left) and *MYC* (right) in LMPP/Early GMPs from FL NUP98::NSD1/WT1ko and CB NUP98::NSD1/WT1ko xenografts based on pseudobulk DESeq2 analysis. The solid lines in each box indicate median expression values, and the height of the box indicates the interquartile range. *EGR1*: FC = 2.77, p = 3.42e-09; *MYC*: FC = -0.69, p = 0.002. **(F)** Schematic experimental overview of our *in vivo* studies to characterize the impact of cytarabine (Ara-C) and menin inhibitor (revumenib) treatment of FL NUP98::NSD1, FL NUP98::NSD1/WT1ko, and CB NUP98::NSD1/WT1ko xenografts. **(G)** Engraftment of human CD45+ cells in control, cytarabine-treated, or revumenib-treated FL NUP98::NSD1, FL NUP98::NSD1/WT1ko, and CB NUP98::NSD1/WT1ko xenografts. Error bars represent standard deviation. **p < 0.01 and ***p < 0.001, unpaired t-test; n = 4-19 mice per condition. **(H)** Percentage of CD34+CD117+ enriched LSCs in control, cytarabine-treated, and revumenib-treated NUP98::NSD1 and NUP98::NSD1/WT1ko xenografts. Error bars represent standard deviation. *p < 0.05, ***p < 0.001 and ****p < 0.0001, unpaired t-test. **(I)** UMAP of scRNA-seq data from control, cytarabine- and revumenib-treated FL NUP98::NSD1/WT1ko xenografts, mapped to the merged scRNA-seq reference, contoured by relative abundance density of cells. **(J)** Bar plots showing ORA results for pathways up-regulated in NUP98::NSD1/WT1ko after treatment with cytarabine and revumenib. Significant hallmark pathways are highlighted in bold and marked with an asterisk (*) based on a 0.05 p-value cutoff.

Previous work on *WT1*-mutated Wilms’ tumors reported *EGR1* and *MYC* upregulation as direct consequences of WT1 loss-of-function.^29^ The *MYC* promoter harbors a *WT1* binding motif, which, upon WT1 binding, negatively regulates the *MYC* family.^30,31^ *MYC* overexpression has been implicated in therapy resistance in AML.^32,33^ Moreover, *WT1* and *EGR1* share a DNA-binding motif. Studies have shown that WT1 can suppress *EGR1* expression, and despite the shared binding motif, WT1 and EGR1 regulate distinct and sometimes antagonistic transcriptional networks.^28,34,35^ EGR1 is a known mediator of long-term HSC function and quiescence^36^ and is associated with inflammatory pathways.^37^ Our comparisons of FL NUP98::NSD1/WT1ko versus NUP98::NSD1 stem and progenitor cell populations revealed that *MYC* and *EGR1* were among the highest differentially expressed genes (**Figure 4C and D**). These TFs also exhibited notable motif enrichment based on scATAC-seq analyses (**Supplementary Figure 5C and D**). This pattern was recapitulated in cells derived from NUP98::NSD1 PDX models where *WT1* mutation was associated with *EGR1* overexpression (**Supplementary Figure 5E**). In addition, *EGR1* expression was significantly elevated in FL-originating NUP98::NSD1/WT1ko LSCs compared to their CB-originating counterparts (**Figure 4C, E and Supplementary Figure 5F**). The *MYC* expression pattern was reversed with higher LMPP/Early GMPs expression in CB NUP98::NSD1/WT1ko xenografts (**Figure 4E)**. WT1ko-mediated *MYC* and *EGR1* upregulation may contribute to the expansion of quiescent LSCs, induce altered inflammatory cascades, and promote resistance to therapy associated with *WT1*-mutated AMLs. The downstream effects of *WT1* mutations likely differ to some extent between early and later developmental stages, as reflected by the unequal impact on EGR1 and MYC overexpression in FL and CB LSCs.

### Developmental Stage-Dependent Resistance to Therapy

Our next objectives were to functionally verify the therapeutic resistance of NUP98::NSD1/WT1ko cells and investigate whether the observed transcriptional differences between FL- and CB-derived AMLs influence therapeutic responses *in vivo*. Cytarabine chemotherapy (Ara-C) was administered for five days to serially established FL NUP98::NSD1, FL NUP98::NSD1/WT1ko, and CB NUP98::NSD1/WT1ko 4th generation xenograft mice (**Figure 4F**). Cytarabine did not affect overall engraftment in FL NUP98::NSD1 or CB NUP98::NSD1/WT1ko xenografts but significantly reduced it in their FL NUP98::NSD1/WT1ko counterparts (**Figure 4G**). Importantly, in contrast to FL NUP98::NSD1 LSCs, which were targeted by cytarabine, FL NUP98::NSD1/WT1ko LSCs exhibited marked resistance and were significantly enriched post-chemotherapy (**Figure 4H**). LSC populations in CB NUP98::NSD1/WT1ko xenografts were also resistant to the chemotherapy.

Menin is a nuclear protein that binds to lysine methyltransferase 2A (*KMT2A*) and other partners to form a complex that regulates gene expression through epigenetic modulation of transcription.^38^ Menin inhibition has emerged as a promising therapy for *KMT2A*-rearranged AMLs and other specific fusion-driven leukemias. However, common co-mutations (*e.g.*, in *WT1*) may attenuate the therapeutic effectiveness of menin inhibition.^39^ Recent studies also highlighted that acquiring *MEN1* mutations and epigenetic rewiring under therapeutic pressure can drive resistance to menin inhibition in LSCs.^40^ To investigate these issues in our experimental system, we fed mice with menin inhibitor (revumenib)-containing chow for two weeks (**Figure 4F**). We found that FL NUP98::NSD1 and FL NUP98::NSD1/WT1ko xenograft mice were entirely resistant to revumenib; we observed no effects on overall engraftment and detected a significant increase in the percentage of CD34+CD117+ enriched LSCs following treatment (**Figure 4G and H**). Surprisingly, CB NUP98::NSD1/WT1ko xenografts exhibited substantial sensitivity to revumenib; significant reductions in engraftment and LSC fractions were observed. These findings suggest that *WT1*-mutant NUP98::NSD1-driven AMLs exhibit significant developmental stage-dependent therapeutic resistance and illustrate the impact of hierarchical and transcriptional differences between FL and CB-originating AML models on therapeutic responsiveness.

To further characterize the cytarabine and revumenib resistance of FL NUP98::NSD1/WT1ko xenografts, we performed scRNA-seq on sorted CD45+ cells from control and treated FL NUP98::NSD1/WT1ko xenografts. Overlaying scRNA-seq data from treated xenografts onto our reference UMAP revealed significant enrichment of LMPP/Early GMPs in both cytarabine- and revumenib-treated NUP98::NSD1/WT1ko xenografts (**Figure 4I**). A comparative analysis of the percentages showed a significantly higher proportion of LMPP/Early GMPs, particularly in revumenib-treated xenografts (**Supplementary Figure 5G**). In contrast, cytarabine-treated xenografts displayed a greater abundance of lymphoid populations, which were depleted following revumenib treatment. Pathway analysis of the NUP98::NSD1/WT1ko resistant xenografts post-chemotherapy revealed significant enrichment of oxidative phosphorylation (OXPHOS) and *MYC* pathways, both known mediators of chemotherapy resistance in AML^33^ (**Figure 4J**). Moreover, revumenib treatment also led to the upregulation of MYC-regulated pathways in resistant leukemia cells, suggesting that this response accounts, at least in part, for resistance to menin inhibition. Unfortunately, direct therapeutic targeting of MYC remains challenging.^41^ Together, our results demonstrate that the acquisition of *WT1* mutations confers significant resistance to conventional chemotherapy and that the developmental origin of leukemia determines resistance to menin inhibitors.

### Quiescent LSCs Exhibit a Distinct, Fatty Acid Oxidation-Driven Metabolic Program that Underlies Cellular Dependency on OXPHOS

We next investigated how metabolic dependencies in NUP98::NSD1/WT1ko AML support LSC quiescence and self-renewal. Single-cell-normalized signature enrichment showed pronounced OXPHOS and ROS pathway enrichment in FL NUP98::NSD1 fusion leukemia xenografts, specifically in quiescent LMPP/Early GMPs (**Figure 5A**). This population is significantly expanded following WT1ko (**Figure 3B**); thus, we hypothesized that NUP98::NSD1 AML with *WT1* mutations highly depends on OXPHOS. Consistent with scRNA-seq data, flow cytometry-based MitoTracker assays revealed increased mitochondrial numbers in NUP98::NSD1 and NUP98::NSD1/WT1ko cells compared to control FL CD34+ cells (**Figure 5B**). We then tested whether the higher OXPHOS signature of NUP98::NSD1/WT1ko cells had functional significance by assessing oxygen consumption using Seahorse assays. We found significantly higher oxygen consumption rates (OCR) in NUP98::NSD1/WT1ko compared to NUP98::NSD1 leukemia cells (**Figure 5C**). Intermediates generated through glycolysis, glutamine metabolism, or fatty acid oxidation (FAO) can feed into and drive OXPHOS.^42^ Thus, we individually inhibited upstream carbohydrate, amino acid, and lipid metabolic pathways in NUP98::NSD1/WT1ko cells to identify potential impacts on OCR (**Figure 5D**). Inhibition of glycolysis and glutamine metabolism with the glucose analog 2DG and the selective allosteric glutaminase inhibitor BPTES, respectively, resulted in slight, statistically insignificant OCR reductions. In contrast, inhibiting FAO using etomoxir, a carnitine-palmitoyl transferase 1 (CPT1) inhibitor, significantly reduced maximal respiration and spare respiratory capacity (SRC). Single-cell-normalized signature enrichment using a fatty acid metabolism gene set further confirmed that FAO is particularly enriched in quiescent LSCs, concurrent with OXPHOS enrichment (**Figure 5E**). These results suggest that quiescent LSCs, expanded by *WT1* mutation, exhibit a heightened reliance on FAO, which drives their increased dependency on OXPHOS in NUP98::NSD1/WT1ko AML.

**Figure 5.**
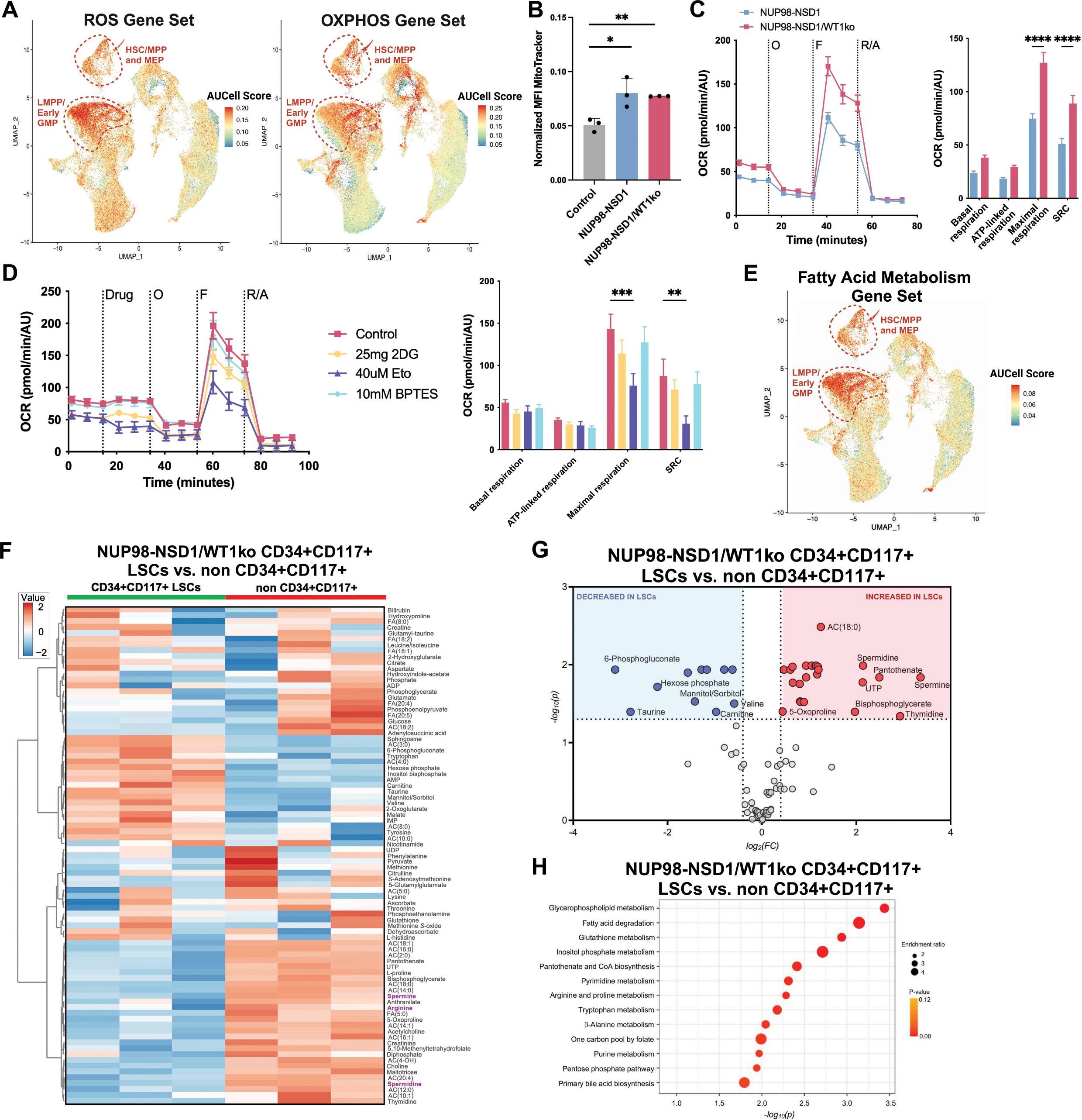
Quiescent LSCs Depend on Fatty Acid Oxidation to Sustain OXPHOS and Maintain Leukemic Self-Renewal. **(A)** UMAP of the merged scRNA-seq reference showing the hallmark ROS and OXPHOS gene set signature enrichment using AUCell. **(B)** Normalized Median Fluorescence Intensity (MFI) in cultured FL-derived CD34+ HSPCs, NUP98::NSD1 and NUP98::NSD1/WT1ko leukemia cells using MitoTracker assay. MFI was normalized by cell size using the median FCS-A value of the target cells. Error bars represent standard deviation. *p < 0.05 and **p < 0.01, unpaired t-test; n = 3 biological replicates per condition. **(C)** Oxygen Consumption Rate (OCR) in NUP98::NSD1 and NUP98::NSD1/WT1ko leukemia cells using Seahorse assays. Analysis of basal respiration, ATP-linked respiration, maximal respiration, and spare respiration in NUP98::NSD1 and NUP98::NSD1/WT1ko leukemia cells. Error bars represent SEM. ****p < 0.0001 using two-way ANOVA; n = 5 biological replicates per condition. O=oligomycin; F=FCCP [Carbonyl cyanide-4 (trifluoromethoxy) phenylhydrazone]; R=rotenone; A=antimycin A. **(D)** OCR in NUP98::NSD1/WT1ko cells under different inhibitor treatments using Seahorse assays. Analysis of basal respiration, ATP-linked respiration, maximal respiration, and spare respiration under inhibitor-treated and control conditions. Error bars represent SEM. **p < 0.01 and ***p < 0.001 using a two-way ANOVA test; n = 5 biological replicates per condition. 2DG=2-deoxyglucose; Eto=etomoxir; BPTES=bis-2-(5-phenylacetamido-1,2,4-thiadiazol-2-yl)ethyl sulfide. **(E)** UMAP of the merged scRNA-seq reference showing hallmark fatty acid metabolism gene set signature enrichment using AUCell. **(F)** Heatmap of metabolomics in CD34+CD117+ enriched LSCs and non-CD34+CD117+ cells from FL NUP98::NSD1/WT1ko xenograft. n = 3 biological replicates per condition. **(G)** Volcano plots of differentially abundant metabolites between CD34+CD117+ enriched LSCs and non-CD34+CD117+ cells. Significant changes are considered for FC > 1.5 and FDR p < 0.05 using an unpaired t-test followed by Benjamini-Hochberg post hoc test; n = 3 biological replicates per condition. **(H)** Pathway enrichment bubble plot for differentially abundant metabolites between CD34+CD117+ enriched LSCs and non-CD34+CD117+ cells.

Our next goal was to identify vital metabolic programs utilized by quiescent LSCs in more detail. We previously demonstrated that the most stem-like LSCs reside in the CD34+CD117+ fraction. Thus, we sorted CD34+CD117+ and non-CD34+CD117+ fractions (**Figure 3F and G**) and performed mass spectrometry-based metabolomics analyses to quantify individual metabolites. The results revealed distinct metabolic signatures in the two groups (**Figure 5F and Supplementary Table S4**). Recent studies unveiled a high dependency of LSCs on polyamines^43^, and our differential metabolite analyses showed a significant elevation of polyamines in CD34+CD117+ enriched LSCs, including spermine, spermidine, and the polyamine precursor arginine (**Figure 5G**). Our studies also revealed that several subtypes of acylcarnitines (ACs) and fatty acids (FAs) were significantly more abundant in CD34+CD117+ enriched LSCs. Pathway analysis of the differential metabolites demonstrated that fatty acid degradation and glycerophospholipid metabolism were the most enriched pathways in CD34+CD117+ compared to non-CD34+CD117+ enriched LSC fractions. These results suggest that quiescent LSCs exhibit a unique metabolic profile compared to more differentiated cells and primarily rely on fatty acid metabolism to sustain their high OXPHOS dependency.

### PRDM16 is a Master Regulator of Oncogenic Programs in Quiescent LSCs

Next, we sought to identify novel therapeutic targets and address therapeutic resistance by identifying master regulators of leukemic and metabolomic programs in quiescent LSCs. Regulon analysis on our single-cell dataset with pySCENIC^44^ predicted PRDM16 as one of the top regulons in LMPP/Early GMPs alongside CEBPA, MYC, and HOXB3 (**Figure 6A**). Previous studies linked *PRDM16* expression in adult AMLs with poorer outcomes.^45,46^ In pediatric AML, high *PRDM16* levels have been associated with alterations in DNA methylation at *AP-1* and *STAT5* binding sites.^47^ Consistent with these findings, the *AP-1* complex, including *JUN* and *FOS*, showed significant motif enrichment based on scATAC-seq (**Supplementary Figure 5B**); *STAT5A* was identified as a top regulon in LMPP/Early GMPs (**Figure 6A**). A previous characterization of patients with NUP98::NSD1-driven AML identified low *MECOM* and high *PRDM16* expression as hallmarks of this AML subtype.^13^ We validated high *PRDM16* expression in a cohort of 164 patients with NUP98-rearranged AML. *PRDM16* is significantly upregulated in cells from patients with NUP98::NSD1 AML and also expressed in NUP98::KDM5A cases (**Supplementary Figure 6A**). This TF is a critical regulator in maintaining quiescence and stemness.^48^ Interestingly, PRDM16 has been extensively studied in the context of beige fat biogenesis, where it activates mitochondrial and metabolomic programs such as FAO.^49,50^ We hypothesize that PRDM16 is a promising therapeutic target with significant prognostic value in AML risk stratification. Targeting PRDM16, which plays a central role in regulating FAO, OXPHOS, and stem cell quiescence, has the potential to simultaneously address the stemness and metabolomic dependencies of LSCs.

**Figure 6.**
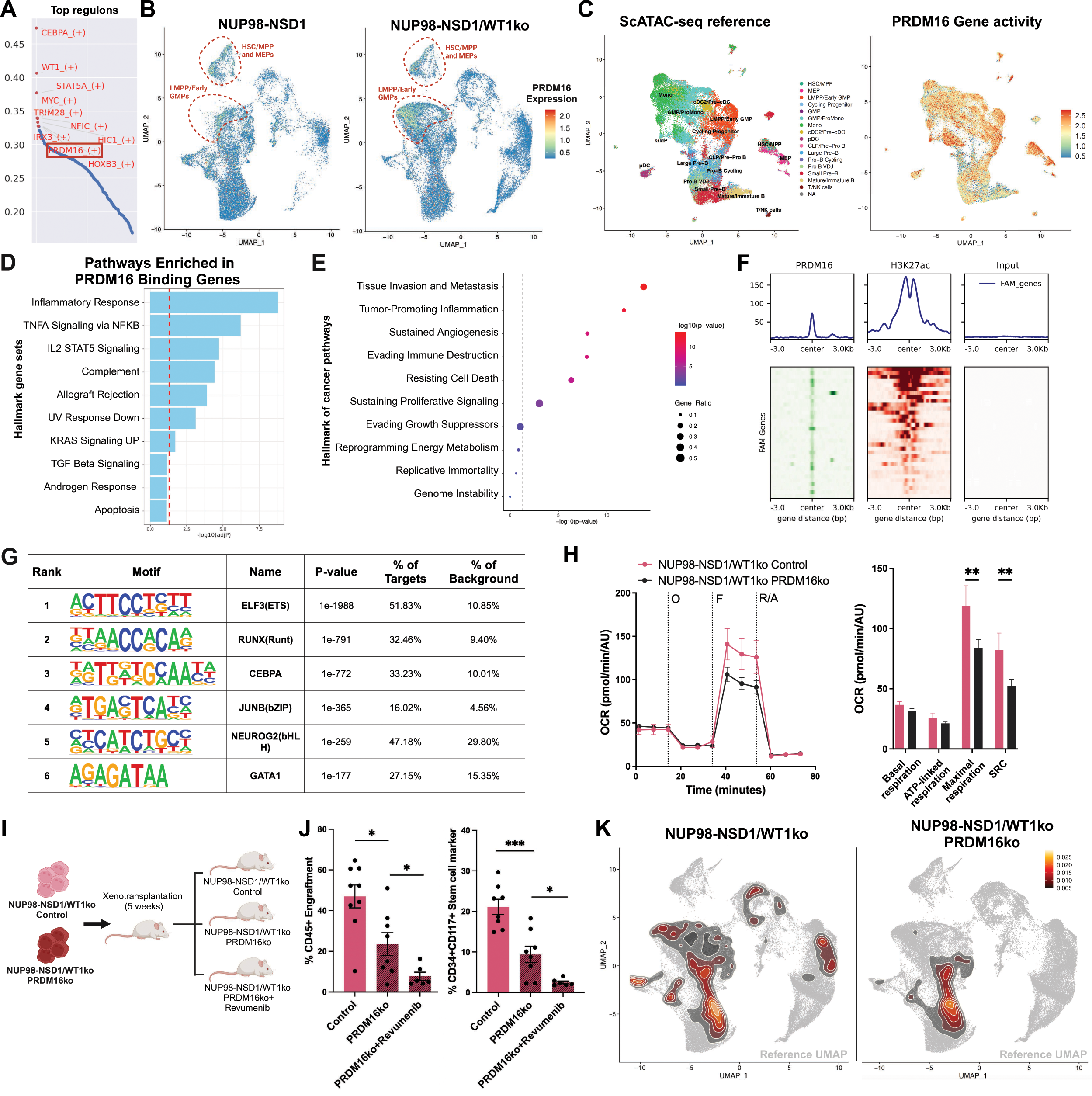
PRDM16 Drives Oncogenic, Metabolic, and Quiescent Programs in LSCs. **(A)** Top 10 regulons in LMPP/Early GMPs based on the reference merged scRNA-seq inferred by pySCENIC. The x-axis indicates regulon specificity score. **(B)** UMAP of *PRDM16* expression in NUP98::NSD1 and NUP98::NSD1/WT1ko secondary xenografts. **(C)** UMAP of the merged scATAC-seq reference, with cell populations annotated based on scRNA-seq data and *PRDM16* gene activity scores displayed on the same UMAP. Control, n = 2 biological replicates; NUP98::NSD1, n = 2 and NUP98::NSD1/WT1ko, n = 4. **(D)** Bar plot showing the hallmark pathway enrichment of PRDM16 binding genes based on PRDM16 ChIP-seq in NUP98::NSD1/WT1ko leukemia cells. The red vertical dashed line indicates the significance value. **(E)** Dot plot displaying the top enriched hallmark cancer pathways associated with PRDM16-binding genes, with dot size representing the gene ratio and color indicating -log10(p-value) significance. The dotted line indicates the threshold for statistical significance, p-value < 0.05. **(F)** Average signal plots (top) and heatmap/tornado plots of transcription start site ±3 kb (bottom) for PRDM16, H3K27ac, and input, showing average signal profiles and signal intensity across Fatty Acid Metabolism (FAM) genes. **(G)** PRDM16 gene binding motif analysis based on ChIP-seq data in NUP98::NSD1/WT1ko leukemia cells. **(H)** OCR in NUP98::NSD1/WT1ko control and NUP98::NSD1/WT1ko PRDM16ko cells detected by Seahorse assays. Analysis of basal respiration, ATP-linked respiration, maximal respiration, and spare respiration. Error bars represent SEM. **p < 0.01 using two-way ANOVA test; n = 6-8 biological replicates per condition. **(I)** Experimental *in vivo* strategy to assess the role of PRDM16. **(J)** Engraftment of human CD45+ cells and CD34+CD117+ enriched LSCs under different conditions. Error bars represent standard deviation. *p < 0.05 and ***p < 0.001 using unpaired t-test, n = 6-9 mice per condition. **(K)** UMAP of scRNA-seq of NUP98::NSD1/WT1ko control and NUP98::NSD1/WT1ko PRDM16ko xenografts mapped on the merged scRNA-seq reference UMAP, contoured with relative abundance density of cells in each sample.

*PRDM16* expression was restricted to the most primitive stem and progenitor cells, as shown in scRNA-seq analyses, and was significantly expanded in NUP98::NSD1/WT1ko xenografts (**Figure 6B**). However, gene activity scores derived from scATAC-seq predicted high *PRDM16* activity across all leukemic xenograft lineages, including differentiated monocytes and lymphocytes, indicating sustained gene accessibility throughout the leukemic hierarchy (**Figure 6C**). Thus, to elucidate the role of PRDM16 in leukemogenesis, we performed ChIP-seq on NUP98::NSD1/WT1ko LSCs using antibodies targeting PRDM16 and the histone 3 lysine 27 acetylation mark (H3K27ac) associated with active chromatin. We identified PRDM16 binding sites (**Supplementary Table S5)**, many of which overlapped with regions marked by high H3K27ac signals (**Supplementary Figure 6B**). These binding sites were frequently located within promoter, intronic, and distal intergenic regions (**Supplementary Figure 6C**). Pathway analysis of PRDM16 target genes revealed roles in inflammatory cascades, NF-κB signaling, and STAT5 pathways (**Figure 6D**) and promoting cancer through regulation of immune evasion and cell death (**Figure 6E**). Notably, we observed significant enrichment of pathways related to oxidative stress, mitochondrial organization, and phospholipid/energy metabolism, underscoring a role for PRDM16 in metabolic programming (**Supplementary Figure 6D**). PRDM16 binding overlapped with numerous fatty acid metabolism genes, marked by high H3K27ac signals at the same sites (**Figure 6F**). For instance, our ChIP-seq dataset showed that PRDM16 binds to regulatory elements of *ASCL1*, *SLC22A15*, and *SLC22A16*, confirming a role for PRDM16 in driving LSC OXPHOS (**Supplementary Figure 6E**). Analysis of motif enrichment at PRDM16 binding sites revealed a significant overrepresentation of binding motifs for *ELF3, RUNX, CEBPA, JUN-B,* and *GATA1,* known for their involvement in stem cell, quiescence, and inflammatory programs (**Figure 6G**).

To assess the dependency of NUP98::NSD1-driven AML on PRDM16, we induced a large *PRDM16* deletion in NUP98::NSD1/WT1ko cells using two guide RNAs (**Supplementary Figure 6F**). Cells were cultured for 4 weeks, and their immunophenotype was assessed at weeks 2 and 4. PRDM16 knock-out (PRDM16ko) resulted in a sustained reduction of CD34+ cells and a significant increase in differentiated CD11b+ myeloid cells (**Supplementary Figure 6G**). Our studies (**Figure 5D and E**) revealed links between PRDM16 and FAO, the main driver of OXPHOS in NUP98::NSD1 LSCs. Thus, we tested whether PRDM16 mediates this metabolic LSC dependency using Seahorse assays and found decreased OCR in PRDM16ko cells compared to controls (**Figure 6H).** Further, we assessed OCR changes following PRDM16ko with and without acute treatment with the FAO inhibitor etomoxir. PRDM16ko and etomoxir treatment resulted in similar reductions in OCR; the combination was not additive (**Supplementary Figure 6H**), pointing to shared mechanisms leading to OCR changes. We further explored the dependency of PRDM16 in both healthy and leukemic hematopoiesis by knocking out PRDM16 in NUP98::NSD1/WT1ko leukemia cells and wild-type FL CD34+ HSPCs, plating the cells, and growing them in methylcellulose colony-forming media (**Supplementary Figure 6I**). PRDM16ko did not affect healthy hematopoiesis (*i.e.,* assessed in wild-type FL HSPC responses) but impacted this process in settings of leukemia as shown by decreased colony formation and percentage of CD34+ cells and promotion of myeloid differentiation in NUP98::NSD1/WT1ko leukemia cells. Further, culturing PRDM16ko-edited NUP98::NSD1/WT1ko leukemia cells led to an increase in late apoptotic cells, in contrast to PRDM16ko-edited wild-type FL CD34+ HSPCs, suggesting that PRDM16 may function as an anti-apoptotic mediator in LSCs (**Supplementary Figure 6J**). These findings indicate a selective dependency of the LSC but not the healthy hematopoietic compartment on PRDM16, revealing an attractive, previously unappreciated opportunity for therapeutic intervention.

To determine whether our observations using cellular approaches are recapitulated *in vivo*, we assessed the reliance of NUP98::NSD1-driven AML on PRDM16 by knocking out PRDM16 in NUP98::NSD1/WT1ko cells derived from xenografts and re-transplanting them back into NSG mice (**Figure 6I**). We found significantly reduced overall engraftment of NUP98::NSD1/WT1ko cells upon PRDM16ko (**Figure 6J**). Flow cytometry analyses of the xenotransplanted mice revealed that PRDM16ko targeted the LSC-enriched CD34+CD117+ fraction and reduced its proportion by half. We then evaluated the therapeutic potential of PRDM16ko in overcoming menin inhibitor resistance by subjecting a subset of mice xenotransplanted with PRDM16ko-edited NUP98::NSD1/WT1ko leukemia cells to a two-week revumenib treatment. The combination of PRDM16ko and menin inhibition further reduced the CD34+CD117+ fraction and overall engraftment, indicating a potential role for PRDM16ko in mitigating menin inhibitor resistance. Our finding that menin inhibitor resistance is predominantly driven by MYC-regulated pathways (**Figure 4J**) is likely explained by our ChIP-seq data, which shows PRDM16 binding to the *MYC* locus (**Supplementary Figure 6K**). This suggests that PRDM16ko disrupts the MYC-driven menin inhibition resistance program. ScRNA-seq analyses of control and PRDM16ko xenografts revealed that the former displayed a balanced representation of stem, progenitor, and differentiated cells (**Figure 6K and Supplementary Figure 6L**). In contrast, PRDM16ko xenografts showed reduced LMPP/Early GMPs and were predominantly composed of differentiated ProMono and Monocytes. Together, our results establish PRDM16 as a critical regulator of the most stem-like LSCs in NUP98::NSD1-rearranged AML. Knocking out PRDM16 significantly reduced leukemia burden by specifically eradicating primitive LSCs and had negligible effects on healthy hematopoiesis. Moreover, combining PRDM16 targeting with existing therapeutic approaches for AML resulted in significant additive effects, thus demonstrating the high potential of this strategy to overcome therapeutic resistance.

### Oncofetal Signatures Predict Poor Outcomes

Given the developmental dependency of patients with NUP98-rearranged AML, we investigated whether enrichment for transcriptional signatures associated with fetal-derived LSCs correlate with clinical outcomes in patients. In our patient cohort (n = 164), the age of diagnosis varied significantly according to the fusion partner, as expected (**Supplementary Figure 7A**).^13^ NUP98::KDM5A cases were diagnosed in the youngest children, with an average of 2.5 years of age. NUP98::NSD1 fusions exhibited the broadest diagnostic age range, from infancy to adulthood, with an average age of 10 years at diagnosis. NUP98-rearranged AMLs involving alternative fusion partners had a median age of diagnosis at 7.5 years. Amongst the 104 patients with NUP98::NSD1 AML in this cohort, we segregated cases based on enrichment for a fetal-versus postnatal-originated oncogenic signature, composed of the 100 most significantly expressed genes between fetal and postnatal LMPP/Early GMPs (**Supplementary Table S3**). Remarkably, patients with a more fetal-oncogenic program exhibited substantially lower overall and event-free survival (**Figure 7A and Supplementary Figure 7B**).

**Figure 7.**
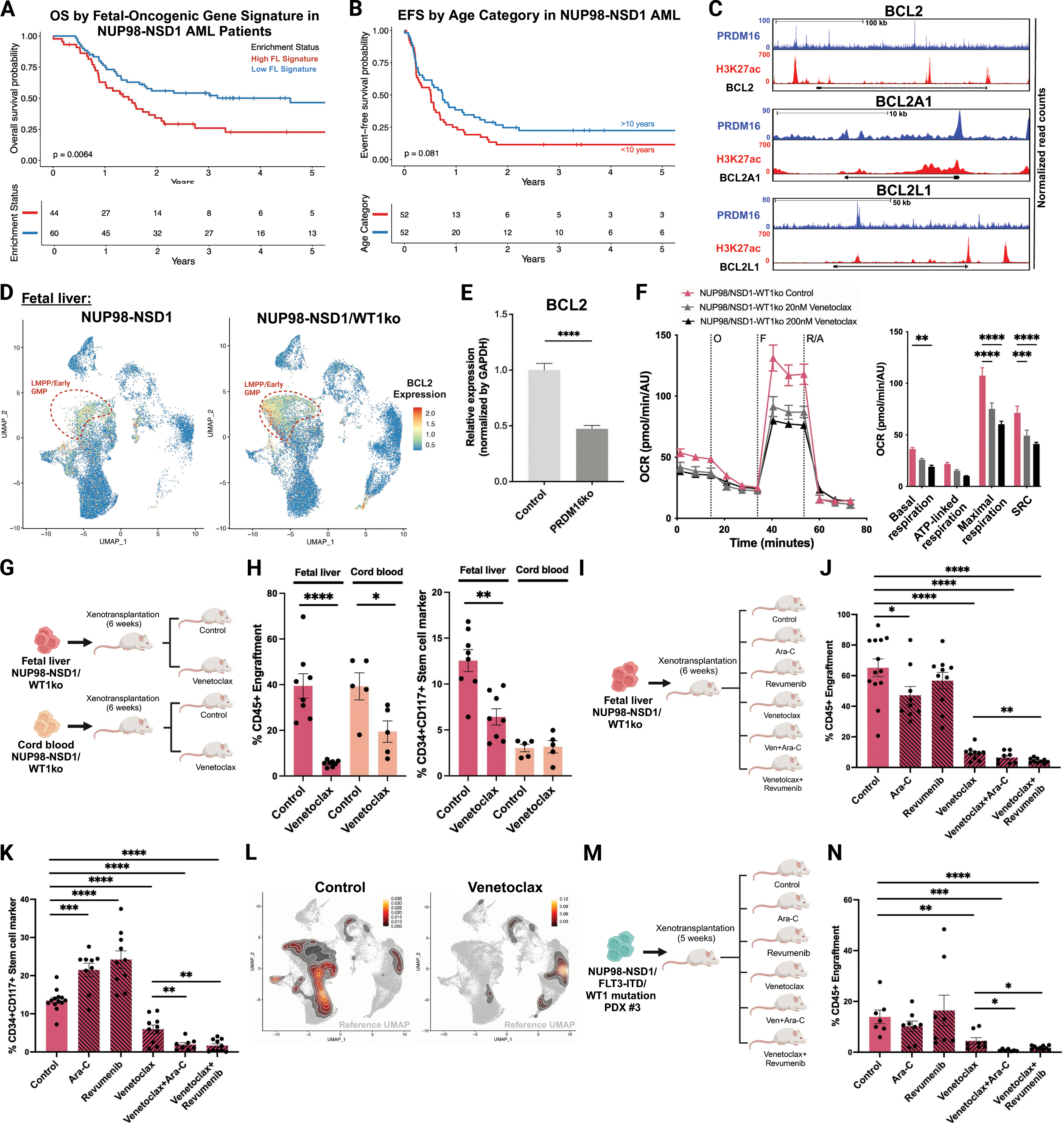
Targeting BCL2 Dependency in Fetal-Origin LSCs with Venetoclax and Combination Therapies Diminished AML. **(A)** Overall survival of patients with NUP98::NSD1 AML stratified by fetal-oncogenic gene signature. The light-blue and red curves show the survival of patients with low and high fetal-oncogenic gene signatures, respectively. Kaplan-Meier method and log-rank test; n = 104 patients. **(B)** Event-free survival of patients with NUP98::NSD1 AML stratified by age. The light-blue and red curves show patients above and below 10 years of age, respectively. Kaplan-Meier method and log-rank test; n = 104 patients. **(C)** ChIP-seq tracks at *BCL2*, *BCL2A1*, and *BCL2L1* loci from NUP98::NSD1/WT1ko leukemia cells for PRDM16 and H3K27ac; n = 3 biological replicates per condition. **(D)** UMAP of *BCL2* expression in FL NUP98::NSD1 and NUP98::NSD1/WT1ko secondary xenografts. **(E)** *BCL2* mRNA expression in NUP98::NSD1/WT1ko leukemia cells with control or PRDM16ko detected by RT-qPCR. Error bars represent standard deviation; ****p < 0.0001 using unpaired t-test; n = 3 biological replicates per condition. **(F)** OCR in NUP98::NSD1/WT1ko leukemia cells with venetoclax treatment using Seahorse assays. Analysis of basal respiration, ATP-linked respiration, maximal respiration, and spare respiration. Error bars represent SEM; **p < 0.01, ***p < 0.001, ****p < 0.0001 using two-way ANOVA test; n = 5 biological replicates per condition. **(G)** Schematic experimental overview of *in vivo* assessment of venetoclax in FL and CB NUP98::NSD1/WT1ko xenotransplanted mice. **(H)** Engraftment of human CD45+ cells and percentage of CD34+CD117+ enriched LSCs. Error bars represent standard deviation, *p < 0.05, **p < 0.01 and ****p < 0.0001 using unpaired t-test, n = 5-8 mice per condition. **(I)** Experimental *in vivo* strategy to assess effects of cytarabine (Ara-C), revumenib, venetoclax, and combinations of (a) cytarabine + venetoclax and (b) revumenib + venetoclax on FL NUP98::NSD1/WT1ko cells. **(J)** Engraftment of human CD45+ cells. Error bars represent standard deviation. *p < 0.05, **p < 0.01 and ****p < 0.0001 using unpaired t-test, n = 8-13 mice per condition. (K) Percentage of CD34+CD117+ enriched LSCs. Error bars represent standard deviation. **p < 0.01, ***p < 0.001 and ****p < 0.0001 using unpaired t-test, n = 8-13 mice per condition. **(L)** UMAP visualization of scRNA-seq data from NUP98::NSD1/WT1ko: control versus venetoclax-treated conditions. Data are mapped on the merged scRNA-seq reference, contoured with the relative abundance density of cells in each sample. **(M)** Experimental *in vivo* strategy to assess effects of cytarabine (Ara-C), revumenib, venetoclax, and combinations of (a) cytarabine + venetoclax and (b) revumenib + venetoclax in a relapsed NUP98::NSD1/FLT3-ITD/WT1-mutant AML PDX model. **(N)** Engraftment of human CD45+ cells. Error bars represent standard deviation. *p < 0.05, **p < 0.01, ***p < 0.001 and ****p < 0.0001 using unpaired t-test, n = 7-10 mice per condition.

Furthermore, younger age at NUP98::NSD1 AML diagnosis, likely reflecting a fetal origin, displayed a trend towards poorer outcomes in this cohort (**Figure 7B**). Interestingly, among the top 100 genes in our fetal-oncogenic signature, we identified CD52, a cell surface marker associated with lineage-plastic MPPs and immune cells^51^, as a gene expressed at higher levels in fetal-derived LSCs compared to CB counterparts (**Supplementary Figure 7C**) and demonstrated significant prognostic value in identifying worse outcomes among NUP98::NSD1 AML cases (**Supplementary Figure 7D**). Although the complete fetal-oncogenic signature significantly surpasses CD52 in predictive power, CD52 can be readily evaluated using existing flow cytometry-based diagnostic approaches and potentially serve as a practical surrogate marker for the oncofetal signature. In conclusion, enrichment for onco-fetal programs is significantly associated with poorer outcomes in a sizable AML patient cohort, guiding our subsequent efforts to target the aggressive fetal-origin NUP98-rearranged AML.

### The Dependency of Fetal-Origin LSCs on BCL2 and OXPHOS Sensitizes AML Cells to Venetoclax

Our last goal was to identify strategies to treat fetal-origin AML, as this sub-type was significantly associated with poorer outcomes in our NUP98::NSD1 model. PRDM16 inhibition holds therapeutic promise for high PRDM16-expressing pediatric AMLs, such as those harboring NUP98::NSD1 rearrangements, but no clinical approaches targeting this pathway have thus far been developed. However, the LSC vulnerabilities identified in this study may offer attractive therapeutic options using clinically available agents. Our observation that quiescent LSCs strongly depend on OXPHOS, a feature even more pronounced in NUP98::NSD1/WT1ko AML, suggests that PRDM16 regulates this metabolic dependency (**Figure 6H**). Additionally, PRDM16 is a potential anti-apoptotic mediator in LSCs (**Supplementary Figure 6J**). ChIP-seq data reported here and elsewhere^52^ revealed that this TF directly binds to the *BCL2* locus and other members of the *BCL2* family (**Figure 7C**). Our scRNA-seq data showed concomitant *BCL2* expression and *PRDM16* transcription, predominantly in the quiescent LMPP/Early GMPs, which increased further in the presence of *WT1* mutations (**Figure 7D**). Notably, PRDM16ko in NUP98::NSD1/WT1ko cells significantly reduced *BCL2* expression (**Figure 7E**). These observations suggest that in LSCs, PRDM16-mediated effects on mitochondrial functions and apoptosis inhibition are partially mediated through direct BCL2 upregulation. Thus, we tested the hypothesis that BCL2 inhibition interferes with PRDM16-regulated signaling in NUP98::NSD1-driven AML. We treated NUP98::NSD1/WT1ko cells and wild-type FL CD34+ HSPCs with various doses of venetoclax, a clinically approved BCL2 inhibitor. Early and late apoptosis markers were assessed by immunophenotyping at 24-, 48-, and 72 hours post-treatment (**Supplementary Figure 7E**). We found that venetoclax significantly reduced overall cell numbers and the percentage of CD34+ cells in NUP98::NSD1/WT1ko cells and markedly increased the percentage of early- and late apoptotic cells. In contrast, the treatment did not affect overall cell numbers, CD34+ percentages, or apoptosis in wild-type FL HSPCs. Seahorse assays performed on venetoclax-treated NUP98::NSD1/WT1ko cells revealed a significant, dose-dependent reduction in OCR, indicating that BCL2 inhibition also targets OXPHOS in LSCs (**Figure 7F**). Combined treatment with venetoclax and etomoxir did not result in additional OCR reductions compared to single treatments (**Supplementary Figure 7F**), suggesting that these agents impair mitochondrial respiration through the same pathway/s.

To evaluate the efficacy of BCL2 inhibition *in vivo*, we treated FL NUP98::NSD1/WT1ko and CB NUP98::NSD1/WT1ko xenografts with venetoclax for two weeks (**Figure 7G**). FL-originating AML had a remarkable response to venetoclax, with engraftment reduced 8-fold to less than 5% of control levels (**Figure 7H**). Moreover, CD34+CD117+ enriched LSCs were specifically targeted and significantly depleted. Consistent with our previous findings, which highlighted differential therapy responses based on whether leukemic transformation occurred in FL or CB HSPCs, CB-originating AML showed a more muted response to venetoclax. This pattern was reflected by the lower degree of CD45+ engraftment reduction and the absence of significant changes in the percentage of CD34+CD117+ enriched LSCs (**Figure 7H**). We hypothesize that the distinct milieu and quiescent state of FL-versus CB-originating LSCs govern their differential responses to venetoclax. Indeed, scRNA-seq data from our CB and FL AML models support this hypothesis: FL xenografts contained a higher proportion of *BCL2*-expressing LSCs, with significantly elevated average *BCL2* expression in LMPP/Early GMPs compared to CB-originating AML (**Figure 7D and Supplementary 7G and H**).

Treatment of resistant FL-originating LSCs with cytarabine and revumenib exacerbated OXPHOS and promoted MYC*-*dependent signatures (**Figure 4J**), further inducing resistance pathways. We hypothesized that combining these therapies with venetoclax could yield a more pronounced response than venetoclax monotherapy because the induction of resistance pathways by cytarabine and revumenib would render LSCs more sensitive to venetoclax. We tested this premise by administering cytarabine, revumenib, venetoclax, venetoclax + cytarabine, or venetoclax + revumenib to FL NUP98::NSD1/WT1ko xenograft mice and compared the responses to those of control-treated animals (**Figure 7I**). As expected, FL NUP98::NSD1/WT1ko xenografts were highly resistant to individual cytarabine or revumenib treatments: we observed no changes in leukemia burden and an enrichment of CD34+CD117+ enriched LSCs (**Figure 7J**). In contrast, venetoclax treatment resulted in a significant reduction in both engraftment and CD34+CD117+ enriched LSCs (**Figure 7K**). Notably, combining venetoclax with either cytarabine or revumenib further reduced engraftment and, most strikingly, nearly eradicated the CD34+CD117+ population. ScRNA-seq analyses of venetoclax-treated xenografts confirmed the near-complete elimination of primitive LSCs, including LMPP/Early GMPs and cycling progenitors, compared to control xenografts (**Figure 7L and Supplementary Figure 7I).** We then validated this treatment strategy in a relapsed NUP98::NSD1/FLT3-ITD/WT1-mutant AML PDX model (**Figure 7M**)^53^. As in our humanized system, the AML PDX model was highly resistant to cytarabine and revumenib monotherapies but showed robust responses to venetoclax. Notably, combined treatment with venetoclax + cytarabine or revumenib almost entirely eradicated AML in the PDX model (**Figure 7N**).

## Discussion

In this study, we explored the developmental specificity of NUP98::NSD1 AML *in vitro* and *in vivo*. We demonstrated that the developmental stage of HSPCs profoundly influences the oncogenic potential of the translocation at the time of transformation. Using CRISPR/Cas9-mediated genome engineering, we endogenously induced NUP98::NSD1 rearrangements and *WT1* co-mutations in human HSPCs from FL, postnatal CB, pediatric, and adult bone marrow. *In vitro*, NUP98::NSD1 fusion conferred the strongest clonal selection advantage to HSPCs obtained earliest in development, with WT1 loss further accelerating this expansion. In contrast, the fusion failed to provide any competitive advantage in BM-derived HSPCs. *In vivo*, xenotransplantation confirmed such findings, as FL-derived NUP98::NSD1-expressing HSPCs developed leukemia, with *WT1* mutations further enhancing leukemogenesis. CB-derived HSPCs required WT1 loss to initiate leukemia, while BM-derived HSPCs remained resistant. GoT-ChA revealed distinct regulatory epigenetic programs mediated by the fusion in FL compared to CB HSPCs. Previous studies have shown dynamic transcriptional and epigenetic remodeling of HSPCs during the FL-to-BM transition.^54,55^ Our findings suggest that the HSPC developmental stage is critical for the oncogenic activity of NUP98::NSD1 rearrangement in AML, which is governed by unique epigenetic and transcriptional landscapes that are actively being reshaped with hematopoietic aging.

We comprehensively characterized the transcriptional and epigenetic landscapes of NUP98::NSD1 AML at single-cell resolution, revealing leukemic hierarchies. We identified a primitive LSC population resembling LMPP/Early GMPs that expands significantly upon WT1 loss. These LSCs were enriched for a quiescent LSPC signature, positioned at the apex of the myeloid/lymphoid bipotent axis, and demonstrated markedly higher self-renewal potential than other populations. Furthermore, we observed elevated expression of *EGR1*, a regulator of HSC quiescence and inflammatory signaling, and *MYC*, a driver of oxidative metabolism and therapy resistance, in NUP98::NSD1/WT1ko fetal LSCs compared to those harboring NUP98::NSD1 and wild-type *WT1*. These findings suggest that WT1 loss promotes a more stem-like, quiescent state while amplifying key oncogenic pathways, further enhancing the aggressive and resistant phenotype of NUP98::NSD1-driven leukemia.

Fetal and postnatal-originating NUP98::NSD1/WT1ko AML exhibited distinct biological and clinical profiles. Fetal-originating leukemic xenografts displayed higher lymphoid output and greater enrichment for quiescence and inflammatory dysregulation in primitive LSCs, correlating with resistance to chemotherapy and menin inhibitors. In contrast, postnatal-originating AML was sensitive to menin inhibition, highlighting ontogeny as a critical determinant of therapeutic response. These findings demonstrate that the temporal origin of LSCs restructures leukemic hierarchy and significantly alters resistance to therapy.

Our study highlights distinct metabolic dependencies of primitive LSCs on OXPHOS and FAO as key drivers of elevated oxidative metabolism. We identified PRDM16 as an indispensable regulator of these metabolic programs, quiescence, and anti-apoptotic mechanisms in LSCs. PRDM16 knockout significantly reduced AML burden by selectively targeting the LMPP/Early GMP-like population while sparing healthy hematopoiesis. Furthermore, PRDM16 disruption impaired MYC-mediated resistance to menin inhibitors in fetal-origin AML, pointing to a role in modulating therapy resistance. Elevated PRDM16 expression has been linked to poor outcomes^45,56^; thus, our findings suggest that PRDM16 is a valuable prognostic marker and therapeutic target. Stratifying pediatric AML cases based on PRDM16 expression and integrating its inhibition into treatment regimens is an attractive strategy to improve outcomes for patients with therapy-resistant leukemia.

Since conventional PRDM16 inhibitors are unavailable, we searched for downstream effectors, determined that PRDM16 directly binds to BCL2, and established that knocking out PRDM16 leads to BCL2 downregulation. The dependency of NUP98::NSD1/WT1ko LSCs on the PRDM16/BCL2 axis, coupled with high OXPHOS metabolism, renders these cells highly sensitive to venetoclax, a highly effective NUP98::NSD1/WT1ko AML treatment, particularly when the fusion oncoprotein originates in fetal HSPCs. This developmental context confers a more quiescent program and higher *BCL2* expression in transformed LSCs. In our humanized and PDX AML models, chemotherapy-induced resistance pathways further increased oxidative reliance, enhancing venetoclax efficacy and facilitating complete leukemia eradication *in vivo*. While menin inhibitors show promise in postnatal-originated AML, our findings highlight venetoclax, particularly in combination with chemotherapy, as a promising and highly effective strategy to overcome resistance in more aggressive fetal-originated AML.

Our study unveils the critical role of ontogeny in leukemia initiation. However, twin and retrospective studies have shown that a significant gap can exist between fetal-initiating mutations and disease onset.^57,58^ For instance, a recent adult myeloproliferative neoplasm study reported that a fetal JAK2-V617F mutation persisted as a preleukemic clone for over three decades before progression to overt disease.^59^ This observation indicates that while the timing of the initial oncogenic event is crucial, postnatal secondary genetic mutations and microenvironmental factors additionally contribute to disease development. Our experimental model offers an innovative approach to investigate how the temporal onset of genetic alterations and the developmental state influence leukemogenesis. The ability to engineer oncogenic mutations in primary human HSPCs isolated at distinct life stages provides an experimentally controlled framework to dissect how ontogeny-regulated programs modulate oncogenic potential and responses to therapy.

Lastly, our work established a clear stratification of patients with NUP98::NSD1-rearranged AML based on age at diagnosis and transcriptional enrichment for the fetal-origin leukemic signature. Our results revealed that younger patients and cases with greater enrichment for oncofetal programs had significantly poorer outcomes and reduced survival. Interestingly, the other predominant AML subtype (*i.e.,* a form that harbors NUP98::KDM5A rearrangements) occurs exclusively in children under five years, whereas NUP98::NSD1 cases span a broader age range. This observation points to distinct developmental dependencies for different NUP98 fusion partners, with NUP98::KDM5A fusion likely originating solely during fetal development and NUP98::NSD1 having a more expansive, pediatric-restricted, oncogenic window. These findings provide compelling evidence that the temporal origin of oncogenic events fundamentally dictates oncogenic potential, influences prognosis, and impacts therapeutic responses. This ontogeny-driven cancer development paradigm is likely also applicable to solid pediatric tumors. For instance, in neuroblastoma, the timing of oncogenic transformation and tumor evolution strongly predicts clinical outcomes^4^. Similarly, the transformation of fetal brain stem cells has been identified as the original event leading to medulloblastoma.^60^ These considerations underscore the potential utility of stratifying pediatric patients with NUP98::NSD1 fusions and other AML sub-types by combining age at diagnosis with fetal-oncogenic signatures. Such a precision medicine approach can help address the significant clinical heterogeneity observed among patients harboring identical oncogenic mutations acquired at distinct stages of ontogeny, enable clinicians to select optimal therapies for individual patients, and facilitate the development of targeted treatments for children with aggressive cancers.

## Supplementary Figure Legends

**Supplementary Figure 1.**
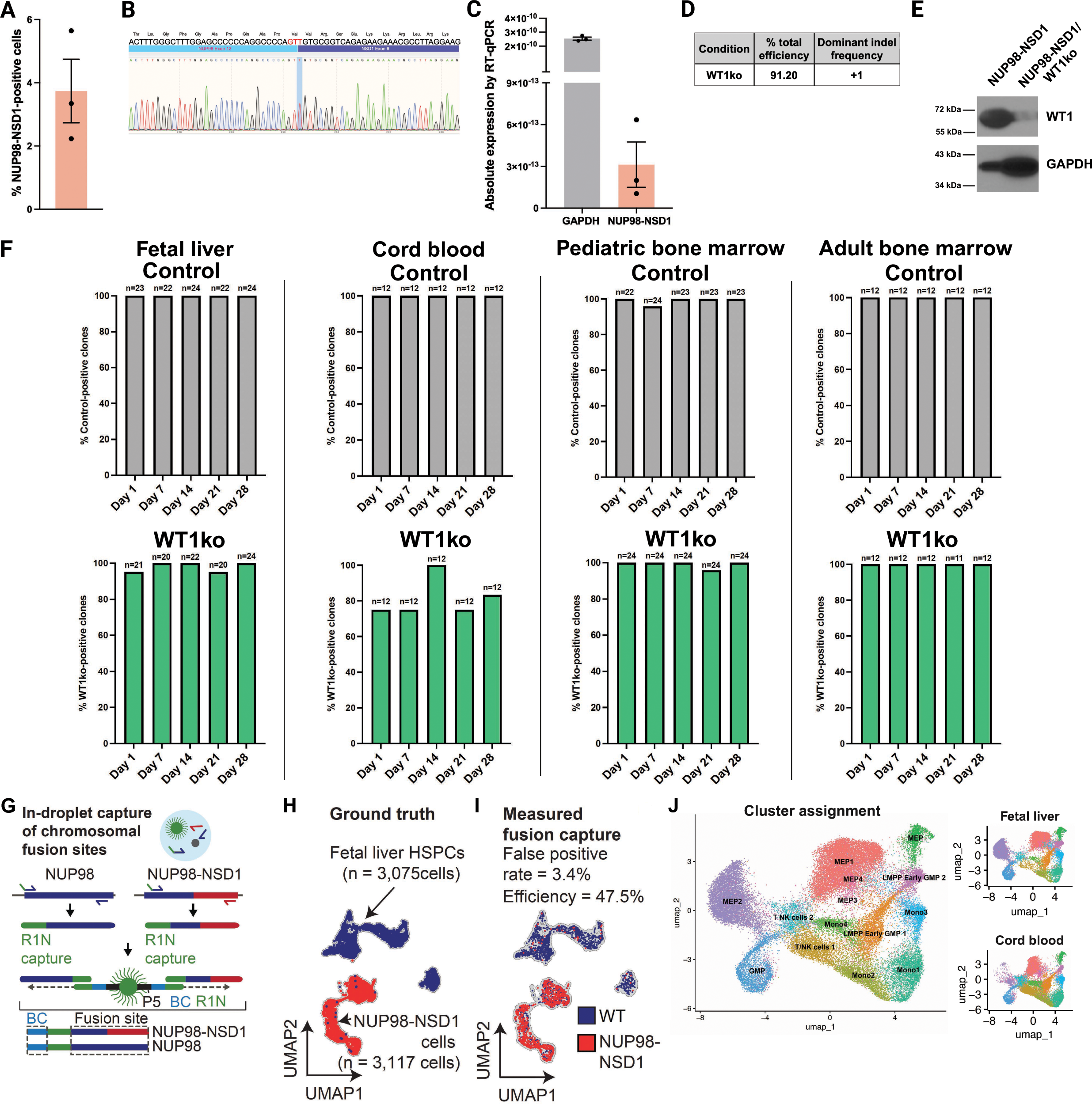
Validation of NUP98::NSD1 Fusion and WT1 Knockout in Human HSPCs. **(A)** CRISPR/Cas9 mediated NUP98::NSD1 editing efficiency in human FL-derived CD34+ cells as determined by droplet digital PCR (ddPCR). Error bars represent standard deviation, n = 3 biological replicates. **(B)** In-frame NUP98::NSD1 fusion sequence identified through Sanger sequencing. **(C)** NUP98::NSD1 mRNA expression levels quantified by RT-qPCR. Error bars represent standard deviation, n = 3 biological replicates. **(D)** *WT1* loss of function editing efficiency was assessed using Sanger sequencing and TIDE analysis. **(E)** Western blot analysis of cultured NUP98::NSD1 and NUP98::NSD1/WT1ko xenografts using anti-WT1 and anti-GAPDH antibodies. **(F)** Percentage of control-edited and WT1ko-edited colonies over four weeks. The number of colonies (n) is indicated on top of each bar. **(G)** Overview of the GoT-ChA experiment using genotyping primers for NUP98::NSD1 and NUP98 wild-type. **(H)** Merged UMAP of a 1:1 mixture of NUP98::NSD1-positive cells and wild-type FL CD34+ HSPCs processed by GoT-ChA, genotyping results overlaid on UMAP clusters. Red dots indicate NUP98::NSD1 fusion-positive cells, blue dots indicate NUP98 wild-type cells, and gray dots represent cells with undefined genotypes. **(I)** Genotyping classification accuracy, with false-positive rate overlay on UMAP clusters. **(J)** UMAP of FL and CB HSPCs four weeks after NUP98::NSD1/WT1ko editing, profiled by GoT-ChA. The left panel shows the merged UMAP, while the right panel shows the contribution of FL HSPCs and CB HSPCs to the merged UMAP.

**Supplementary Figure 2.**
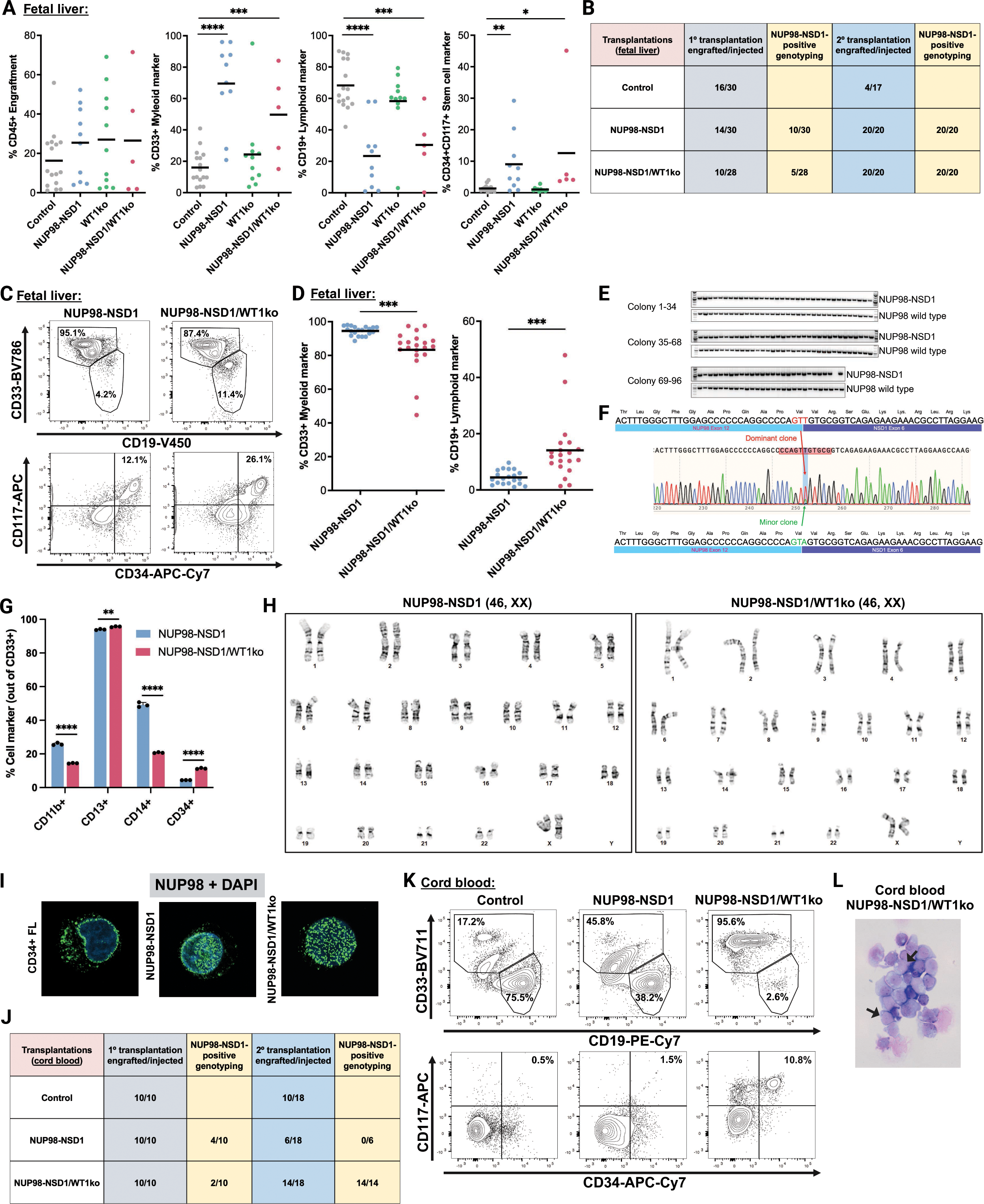
Developmental Context Dictates Leukemic Transformation Potential of NUP98::NSD1-Edited HSPCs. **(A)** Engraftment of human CD45+ cells and the percentage of CD33+, CD19+, and CD34+CD117+ cells in control, NUP98::NSD1, WT1ko, and NUP98-NSD/WT1ko-edited primary xenografts. Only mice with confirmed CRISPR/Cas9 edits (>90% efficiency in control and WT1ko groups and NUP98::NSD1 fusion-positive status) were included in downstream analyses. Lines represent the mean value. *p < 0.05, **p < 0.01, ***p < 0.001 and ****p < 0.0001 using unpaired t-test; n = 5-16 mice per condition from three independent cohorts. **(B)** Number of mice transplanted with fetal liver-derived edited HSPCs, the number of NUP98::NSD1 fusion-positive mice, and the number of mice with engraftment in secondary transplanted xenografts and NUP98::NSD1 fusion-positive genotyping. n = 28-30 mice per condition from 3 independent cohorts. **(C)** Representative flow plots showing CD33+ myeloid, CD19+ B-lymphoid cells, and CD34+CD117+ cells in FL secondary xenografts. **(D)** Engraftment of CD33+ and CD19+ cells from secondary FL NUP98::NSD1 and NUP98::NSD1/WT1ko xenografts. Lines represent the mean value. ***p < 0.001, using unpaired t-test; n = 20 mice per condition. **(E)** Genotyping of 96 individual methylcellulose-cultured colonies from FL NUP98::NSD1 secondary xenografts. The top PCR product indicates the presence of the NUP98::NSD1 fusion, while the bottom band indicates the presence of wild-type NUP98. **(F)** Sanger sequencing of the NUP98::NSD1 break site from bulk cells. **(G)** Immunophenotypic analysis of *in vitro-*cultured cells from secondary FL NUP98::NSD1 and NUP98::NSD1/WT1ko for myeloid differentiation markers (including CD11b, CD13, and CD14) and the stem cell maker CD34. Error bars represent standard deviation. **p < 0.01 and ****p < 0.0001 using unpaired t-test, n = 3 biological replicates. **(H)** Karyotyping of cultured NUP98::NSD1 and NUP98::NSD1/WT1ko cells from secondary FL xenografts, n = 20 metaphases per condition. **(I)** Immunofluorescence using an N-terminal anti-NUP98 antibody in FL-derived CD34+ HSPCs, NUP98::NSD1, and NUP98::NSD1/WT1ko cells. **(J)** Number of mice transplanted with cord blood-derived edited HSPCs, number of NUP98::NSD1 fusion-positive mice, and number of mice with engraftment in secondary transplanted xenografts with NUP98::NSD1 fusion-positive genotyping, n = 10 mice per condition from 2 independent cohorts. **(K)** Representative flow plots of CD33+, CD19+, and CD34+CD117+ cells in control, NUP98::NSD1, and NUP98::NSD1/WT1ko in CB secondary xenografts. **(L)** Morphological analysis of human cells in CB NUP98::NSD1/WT1ko secondary xenografts (arrows indicate blast cells, 63x magnification).

**Supplementary Figure 3.**
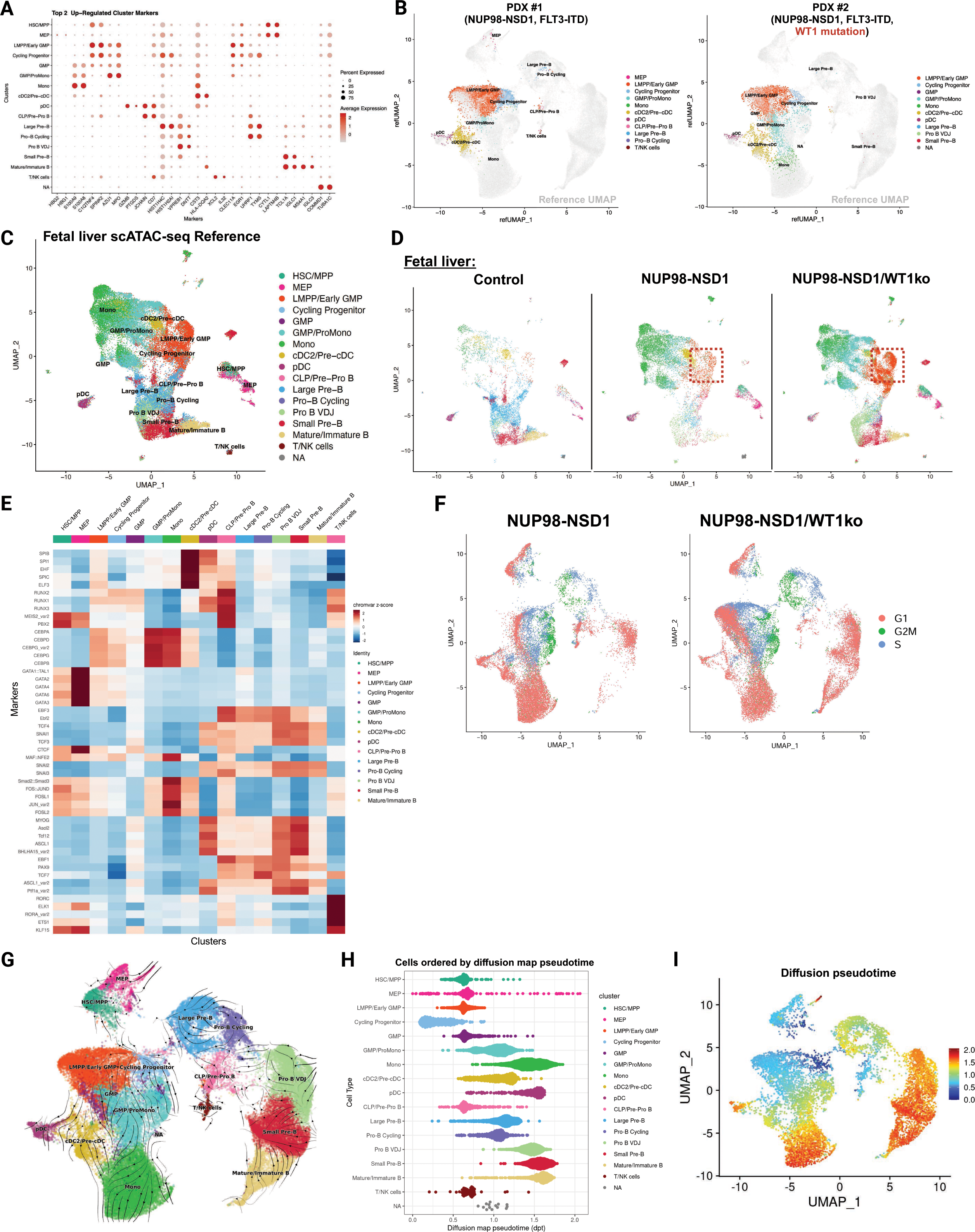
Single-Cell Analysis Reveals Primitive Stem and Progenitor Expansion in NUP98::NSD1 Leukemia. **(A)** Dot plot displaying the percentage of cells expressing (size) and mean expression (color) of the top two most significantly upregulated markers across 17 clusters of the merged scRNA-seq reference. **(B)** UMAP representations of PDX #1 (NUP98::NSD1, FLT3-ITD) and PDX #2 (NUP98::NSD1, FLT3-ITD, WT1 mutation) on the merged scRNA-seq reference. **(C)** Merged scATAC-seq reference composed of FL control, NUP98::NSD1, and NUP98::NSD1/WT1ko secondary xenografts. Control: n = 2 biological replicates; NUP98::NSD1: n = 4; NUP98::NSD1/WT1ko: n = 4. **(D)** Contribution of each condition to the merged scATAC-seq UMAP in (C). **(E)** Heatmap illustrating the top 5 transcription factors in each cell state based on chromVAR z-scores, calculated from the scATAC-seq data. The z-scores represent relative chromatin accessibility associated with each transcription factor across different clusters. **(F)** UMAP visualization of scRNA-seq data showing cell cycle phases in FL xenografts of NUP98::NSD1 and NUP98::NSD1/WT1ko. Cells are color-coded based on their cell cycle phase: G1 (red), G2M (green), and S (blue). **(G)** UMAP illustrating predicted cellular differentiation trajectories based on RNA velocity in the merged scRNA-seq reference. **(H)** Violin plot showing the diffusion map pseudotime across different cell types, with cells ordered along the pseudotime axis. **(I)** UMAP of the merged scRNA-seq reference, colored by the diffusion map pseudotime from (H).

**Supplementary Figure 4.**
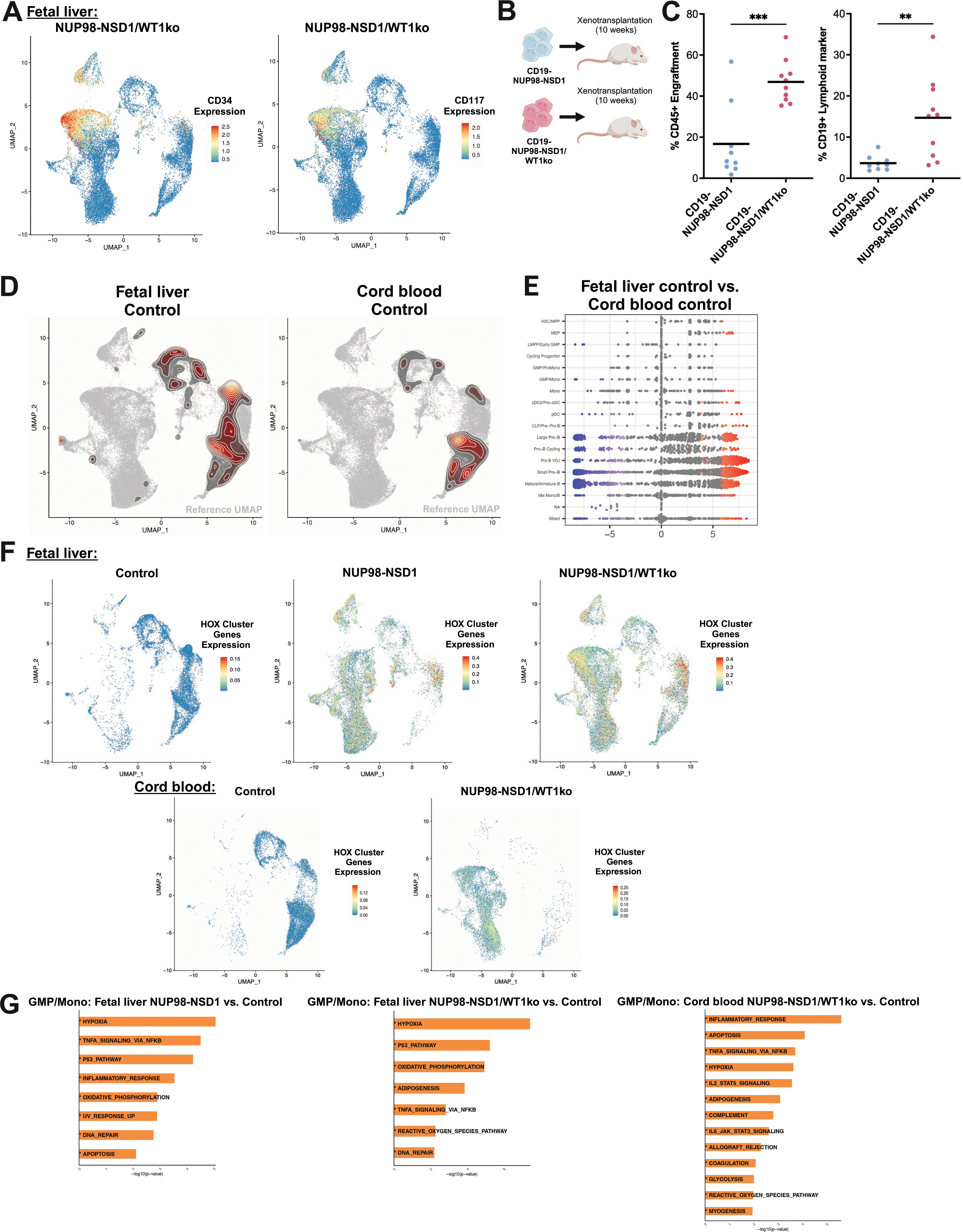
Developmental Origin Shapes Transcriptional and Epigenetic Programs in NUP98::NSD1/WT1ko Leukemia. **(A)** UMAP showing CD34 and CD117 stem cell marker expression in FL NUP98::NSD1/WT1ko. **(B)** Schematic overview of the experimental procedure for serially transplanting CD19-negative cells, myeloid enriched, from FL NUP98::NSD1 and NUP98::NSD1/WT1ko xenografts into NSG mice. **(C)** Engraftment of human CD45+ cells and CD19+ cells in xenografts. Lines represent mean value. **p < 0.01 and ***p < 0.01 using unpaired t-test, n = 9-10 mice per condition. **(D)** UMAP of scRNA-seq data from control secondary xenografts (FL and CB) mapped onto the merged scRNA-seq reference, with contour lines indicating the relative cell abundance in each sample. **(E)** Beeswarm plot displaying the log-fold change between FL control and CB control, analyzed using Milo differential abundance testing. Colored points indicate significant neighborhoods (FDR < 0.01). **(F)** UMAPs of HOX cluster gene expression based on a defined expression module in FL and CB control, FL NUP98::NSD1, FL NUP98::NSD1/WT1ko, and CB NUP98::NSD1/WT1ko. **(G)** Bar plot highlighting the top upregulated hallmark pathways from ORA analysis of differentially expressed genes in GMP/Mono cluster across xenograft comparisons of FL NUP98::NSD1 versus FL control, FL NUP98::NSD1/WT1ko versus FL control, and CB NUP98::NSD1/WT1ko versus CB control. Statistically significant pathways (p-value < 0.05) are bolded and marked with an asterisk (*).

**Supplementary Figure 5.**
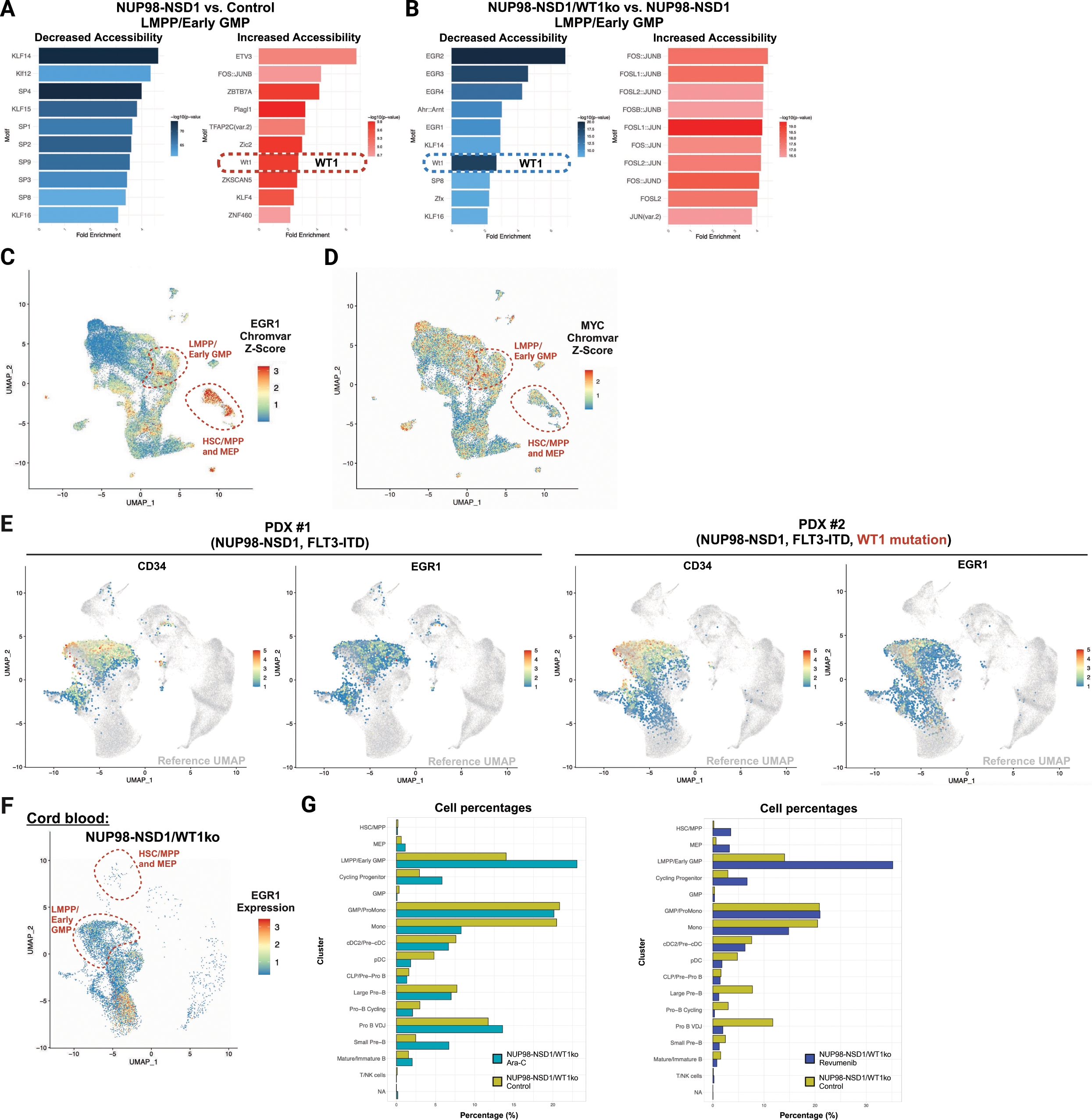
WT1 Loss Amplifies Stemness, EGR1 and MYC Expression in NUP98::NSD1 AML. **(A)** Bar plot displaying the top 10 motifs with reduced (blue) or increased (red) chromatin accessibility in LMPP/Early GMPs of FL NUP98::NSD1 versus FL control, as determined by scATAC-seq analysis. **(B)** Same as (A), but for FL NUP98::NSD1/WT1ko versus FL NUP98::NSD1. **(C)** UMAP visualization of *EGR1* motif enrichment scores using ChromVAR in the merged scATAC-seq dataset. Cells are color-coded by ChromVAR z-score, representing relative motif accessibility across distinct populations. **(D)** UMAP plot of *MYC* motif enrichment scores using ChromVAR in the merged scATAC-seq dataset. **(E)** UMAP plot of *CD34* and *EGR1* gene expression in PDX #1 (NUP98::NSD1, FLT3-ITD) and PDX #2 (NUP98::NSD1, FLT3-ITD, WT1 mutation) overlaid on the merged scRNA-seq UMAP. **(F)** UMAP plot of *EGR1* expression in CB NUP98::NSD1/WT1ko xenografts overlaid on the merged scRNA-seq UMAP. **(G)** Bar plot displaying the percentage of each cell state in NUP98::NSD1/WT1ko and control xenografts treated with cytarabine or revumenib, based on scRNA-seq.

**Supplementary Figure 6.**
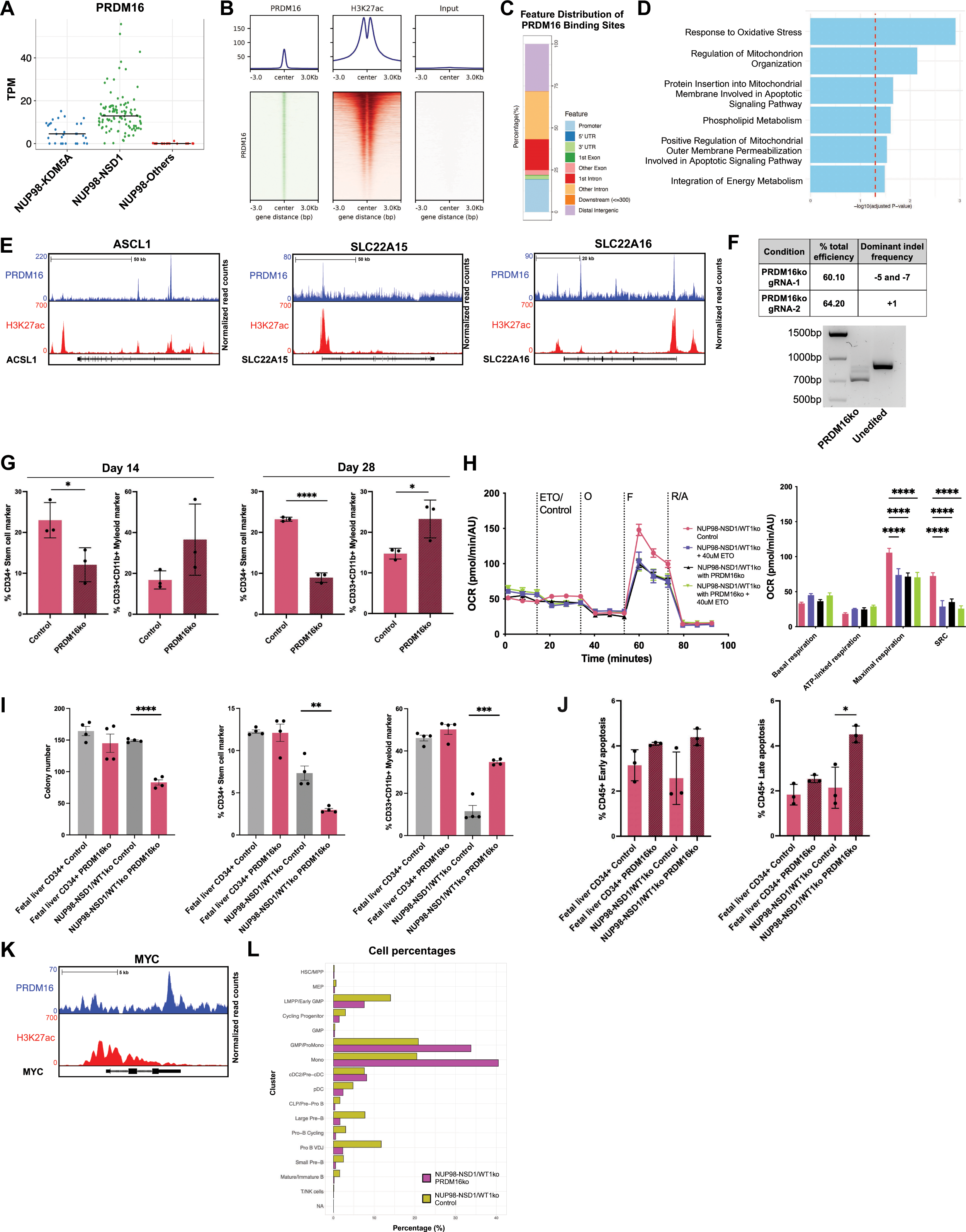
PRDM16 Regulates Leukemic Stemness, Metabolism, and Quiescence in NUP98::NSD1/WT1ko AML. **(A)** *PRDM16* expression in a clinical cohort of NUP98-rearranged pediatric patients. Gene expression was calculated in transcripts per million (TPM), with the line representing the median value. NUP98::KDM5A: n = 30; NUP98::NSD1: n = 104; NUP98 with other gene partners: n = 20. **(B)** Genome-wide average signal plots (top) and heatmap/tornado plots (bottom) of transcription start sites (±3 kb) for PRDM16, H3K27ac, and input, illustrating average signal profiles and signal intensity across PRDM16 binding sites. **(C)** Stacked bar plot depicting the feature distribution of PRDM16 binding sites. **(D)** Bar plot illustrating Gene Ontology (biological process) and Reactome pathway enrichment of PRDM16-binding genes based on ChIP-seq. The dotted line represents the statistical significance threshold (p < 0.05). **(E)** ChIP-seq tracks of PRDM16 and H3K27ac at select FAO-related genes (*ASCL1*, *SLC22A15,* and *SLC22A16*) in NUP98::NSD1/WT1ko leukemia cells. **(F)** Editing efficiency of PRDM16-targeting gRNAs. Representative genotyping results for PRDM16ko and unedited NUP98::NSD1/WT1ko cells. **(G)** Percentage of CD34+ stem cells and CD33+CD11b+ mature myeloid cells in *in vitro*-cultured NUP98::NSD1/WT1ko control and PRDM16ko after 2 and 4 weeks. Error bars represent standard deviation. *p < 0.05 and ****p < 0.0001, determined by unpaired t-test, n = 3 biological replicates per condition. **(H)** OCR in NUP98::NSD1/WT1ko and NUP98::NSD1/WT1ko with PRDM16ko cells treated with or without etomoxir, measured using Seahorse assay. The analysis includes basal respiration, ATP-linked respiration, maximal respiration, and spare respiration. Error bars represent SEM. ****p < 0.0001, determined by two-way ANOVA, n = 6-8 biological replicates per condition. **(I)** Methylcellulose colony formation assay of FL-derived CD34+ HSPCs and NUP98::NSD1/WT1ko leukemia cells with PRDM16ko. The number of colonies, along with the percentage of CD34+ cells and CD33+CD11b+ myeloid cells, were analyzed by flow cytometry. Error bars represent standard deviation. **p < 0.01, ***p < 0.01, and ****p < 0.0001, determined by unpaired t-test. n = 4 biological replicates, each containing more than 20 colonies per condition and replicate. **(J)** Apoptosis analysis in FL-derived CD34+ HSPCs and NUP98::NSD1/WT1ko leukemia cells, 10 days after PRDM16 knockout. The percentages of early and late apoptosis were measured in CD45+ cells. Error bars represent standard deviation. *p < 0.05, determined by unpaired t-test, n = 3 biological replicates per condition. **(K)** ChIP-seq tracks of *PRDM16* and H3K27ac at *MYC* gene in NUP98::NSD1/WT1ko leukemia cells. **(L)** Bar plot displaying the percentage of each cell state in NUP98::NSD1/WT1 PRDM16ko and control xenografts based on scRNA-seq.

**Supplementary Figure 7.**
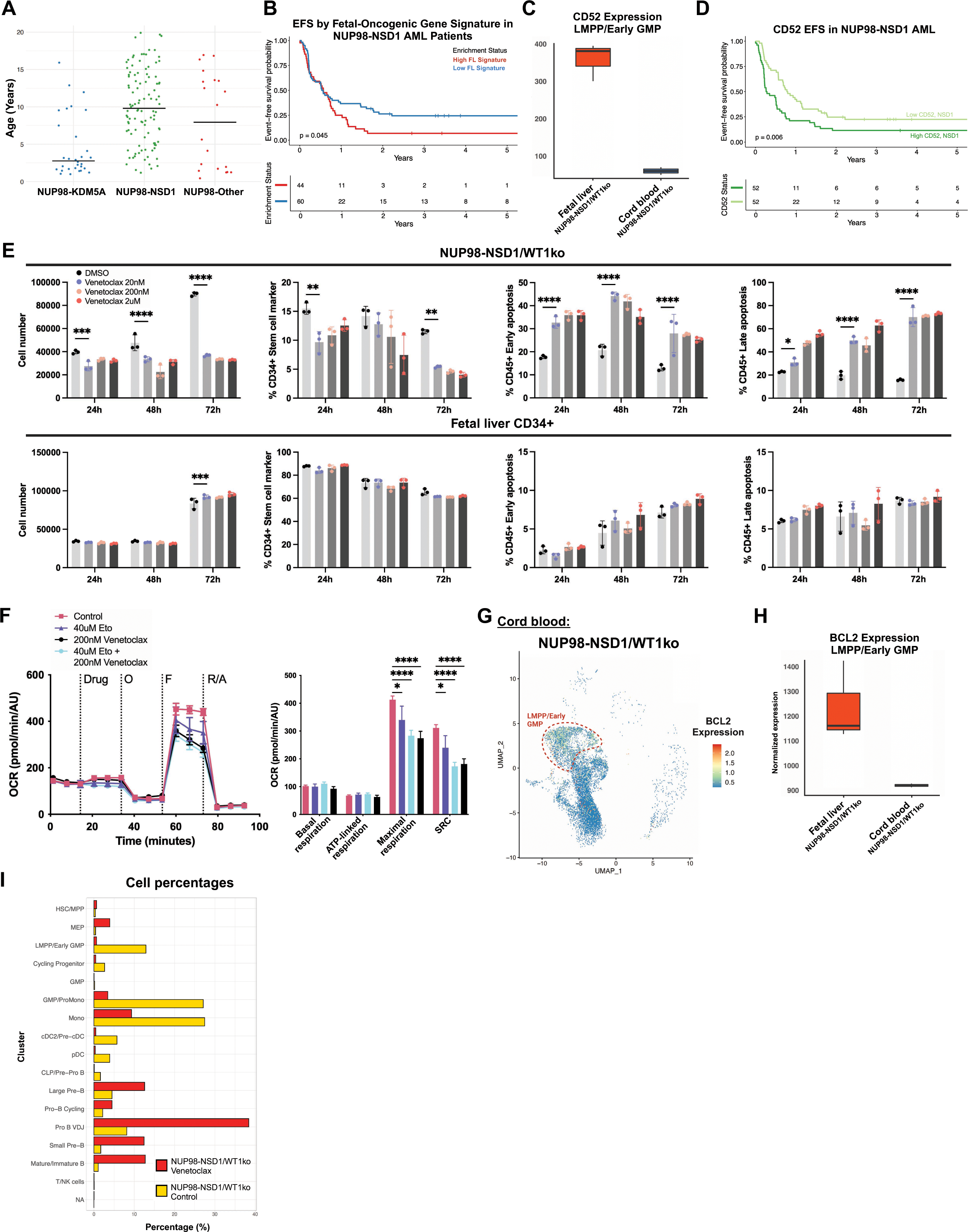
Venetoclax Targets OXPHOS and BCL2 Dependency in Fetal-Origin NUP98::NSD1/WT1ko AML. **(A)** Age distribution in the clinical cohort of NUP98-rearranged pediatric patients: NUP98::KDM5A (n = 30), NUP98::NSD1 (n = 104), and NUP98 with other gene partners (n = 20). **(B)** Event-free survival of NUP98::NSD1 AML patients stratified by the fetal-oncogenic gene signature. The light blue and red curves represent patients with a low or high fetal-oncogenic gene signature, respectively. Data were analyzed using the Kaplan-Meier and log-rank tests, with n = 104 patients. **(C)** Boxplot of *CD52* expression in LMPP/Early GMPs from FL NUP98::NSD1/WT1ko and CB NUP98::NSD1/WT1ko, based on pseudobulk DESeq2 analysis. The solid lines within each box represent the median expression value, and the height of the box indicates the interquartile range. FC = 2.57, p = 7.07e-50. **(D)** Event-free survival of NUP98::NSD1 AML patients stratified by *CD52* expression level. The light green and green curves show the survival of patients with low and high *CD52* expression, respectively. Analysis was performed using the Kaplan-Meier method and log-rank test, n = 104 patients. **(E)** Effects of venetoclax on FL-derived CD34+ HSPCs and NUP98::NSD1/WT1ko leukemia cells *in vitro*. Total cell count, percentage of CD34+ stem cell marker, and early and late apoptotic cells were assessed at multiple time points after venetoclax treatment using flow cytometry. Error bars represent standard deviation, **p < 0.01, ***p < 0.001, and ****p < 0.0001, using unpaired t-test, n = 3 biological replicates per condition. **(F)** OCR in NUP98::NSD1/WT1ko cells treated with control, etomoxir, venetoclax, and a combination of etomoxir and venetoclax, measured by Seahorse assay. Analysis of basal respiration, ATP-linked respiration, maximal respiration, and spare respiration. Error bars represent SEM. *p < 0.05 and ****p < 0.0001, using two-way ANOVA, n = 8-10 biological replicates per condition. **(G)** UMAP of *BCL2* expression in CB NUP98::NSD1/WT1ko xenografts overlaid on the merged scRNA-seq reference. **(H)** Boxplots of *BCL2* normalized expression in LMPP/Early GMPs from FL NUP98::NSD1/WT1ko and CB NUP98::NSD1/WT1ko xenografts based on pseudobulk DESeq2 analysis. The solid lines within each box represent the median expression value, and the height of the box indicates the interquartile range. FC = 0.34, p = 0.015. **(I)** Bar plot displaying the percentage of each cell state in venetoclax-treated and control NUP98::NSD1/WT1ko xenografts based on scRNA-seq data.

## Supplementary Tables

**Table S1.** Differential transcription factor motifs between NUP98::NSD1 fusion-negative and - positive cells in MEPs.

**Table S2.** Comparison of chromatin accessibility of motifs in LMPP/Early GMPs under different conditions

**Table S3.** Differentially expressed genes in LMPP Early GMPs.

**Table S4.** Metabolites detected by mass spectrometry in NUP98::NSD1/WT1ko non-LSC and LSC fractions.

**Table S5.** PRDM16 binding genes in NUP98::NSD1/WT1ko by ChIP-seq.

## Acknowledgments

We thank Reid Joseph and Imani Henry for assistance with sample processing; Travis Dawson, Darwin D’Souza, Rachel Chen, Kai Nie and Seunghee Kim-Schulze from the Human Immune Monitoring Center (Mount Sinai) for single-cell library generation; Linda Lee and Deniz Demircioglu from the Bioinformatics for Next Generation Sequencing (BiNGS) Shared Resource (Mount Sinai) for computational support; Edgardo Ariztia, Guillermo Villegas and Jordi Ochando from the Flow Cytometry CoRE (Mount Sinai) for assistance with flow cytometry; Sebastian Elghaity-Beckley and Joann Arandela from the Hematological Malignancies Tissue Bank (Mount Sinai) for providing a subset of biospecimens for this work; Franco Carlos, Lenny Martinez, Chineta Pullin, Kelly Yamada and Jonathan Cohen from the Center for Comparative Medicine and Surgery (Mount Sinai) for support with mouse work; Damien Laudier and Alan Soto from the Biorepository and Pathology CoRE (Mount Sinai) for assistance with pathology. We thank the Developmental Origin of Health and Disease (DOHaD) Biorepository, including Ya-Wen Chen, Mikal Kizilbash, and Meghana Sreenath, and the Birth Defects Research Laboratory (BDRL). The BDRL group author list includes: Ian A. Glass^1^, Kimberly A. Aldinger^1,2^, Dan Doherty^1^, Ian G. Phelps^1^, Jennifer C. Dempsey^1^, Kevin J. Lee^1^ and Lucinda A. Cort^1^ (Affiliations: #1 University of Washington and #2 Seattle Children’s Research Institute). We thank members of the Wagenblast lab for their comments on the manuscript. Figures were created in BioRender.

## Author Contributions

K.W., S.S., and E.W. designed, performed, analyzed experiments, and prepared figures. N.P., S.C., and D.H. provided computational expertise, analyzed all scRNA-seq, scATAC-seq, and ChIP-seq datasets, and prepared figures. S.A., M.S., I.G.M., C.C., M.Q.A., G.F., and M.Z. assisted with *in vivo* experiments. J.H.P., R.E.R., and S.M. provided clinical cohort data and performed computational analyses. A.H.C.M. performed western blots. N.R. and J.E.C. assisted with Seahorse assays. L.M., D.A.L., and F.I. assisted with GoT-ChA experiments. K.L. performed immunofluorescence assays. Z.D. and M.M.G. assisted with sample processing. J.Z., G.P., and D.J.P. provided metabolism expertise. D.F. and E.B. assisted with ChIP-seq assays. M.B. and D.J.P. carried out metabolomics analysis. T.B., M.W., S.K.T., and B.K.M. provided NUP98-rearranged PDX samples. J.A.R. and A.D. performed metabolomics. I.A.G. and BDRL coordinated patient consent and sample collection. C.M.S. and E.P.P. provided study consultation. Z.C. supervised single-cell library generation and sequencing. S.S. wrote the paper with assistance from K.W., and E.W. E.W. supervised the study.

## STAR METHODS

### KEY RESOURCES TABLE

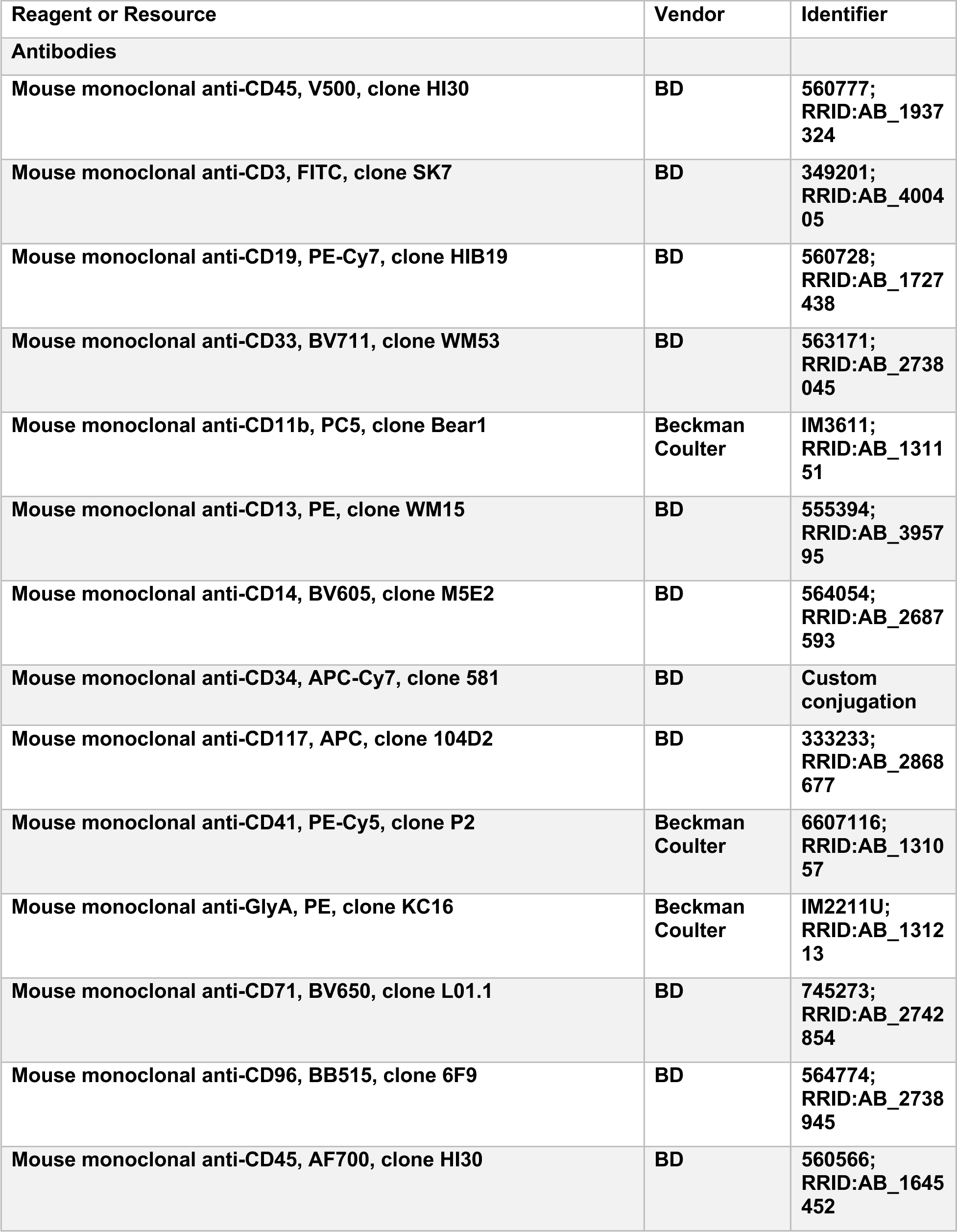
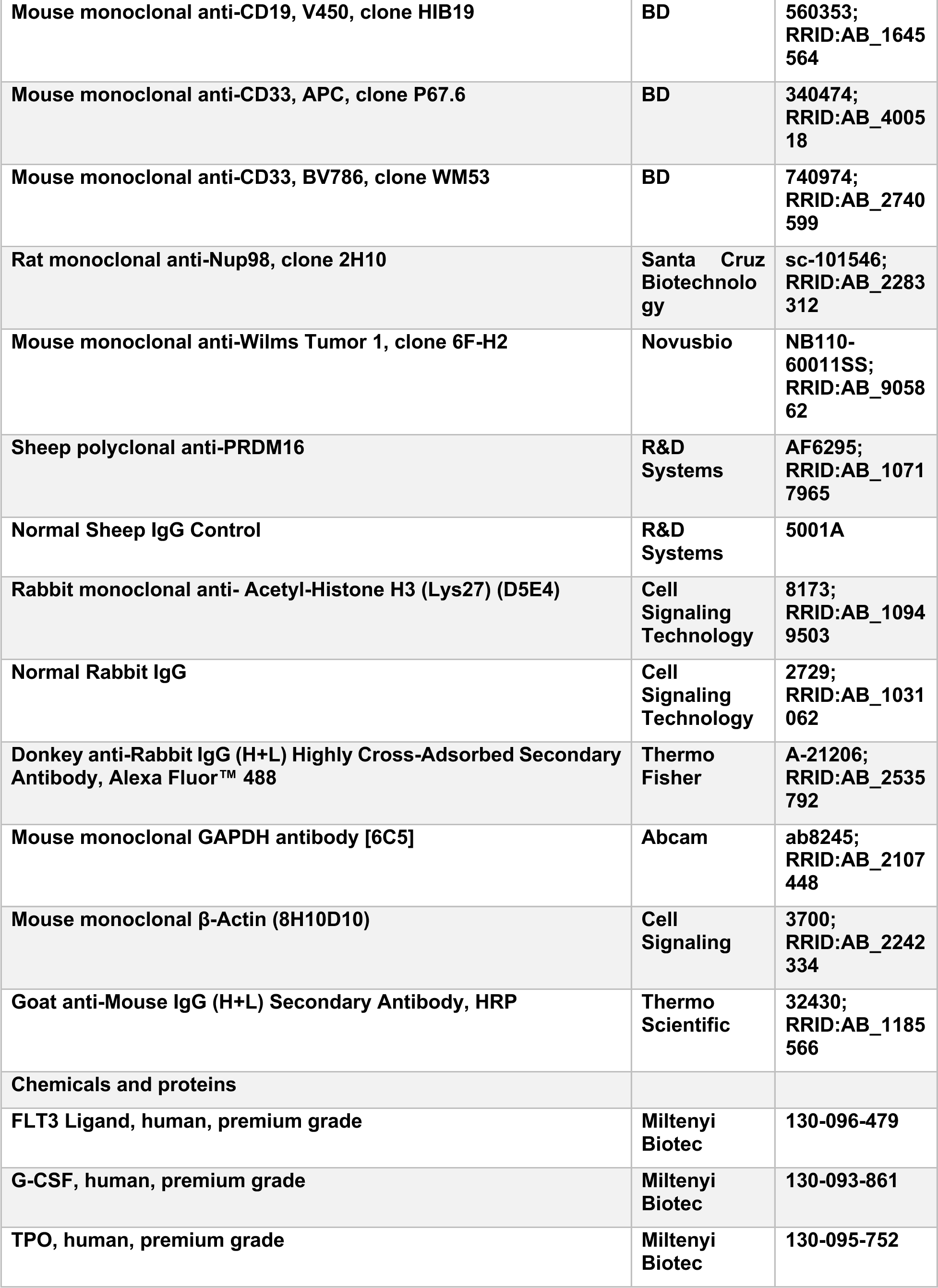
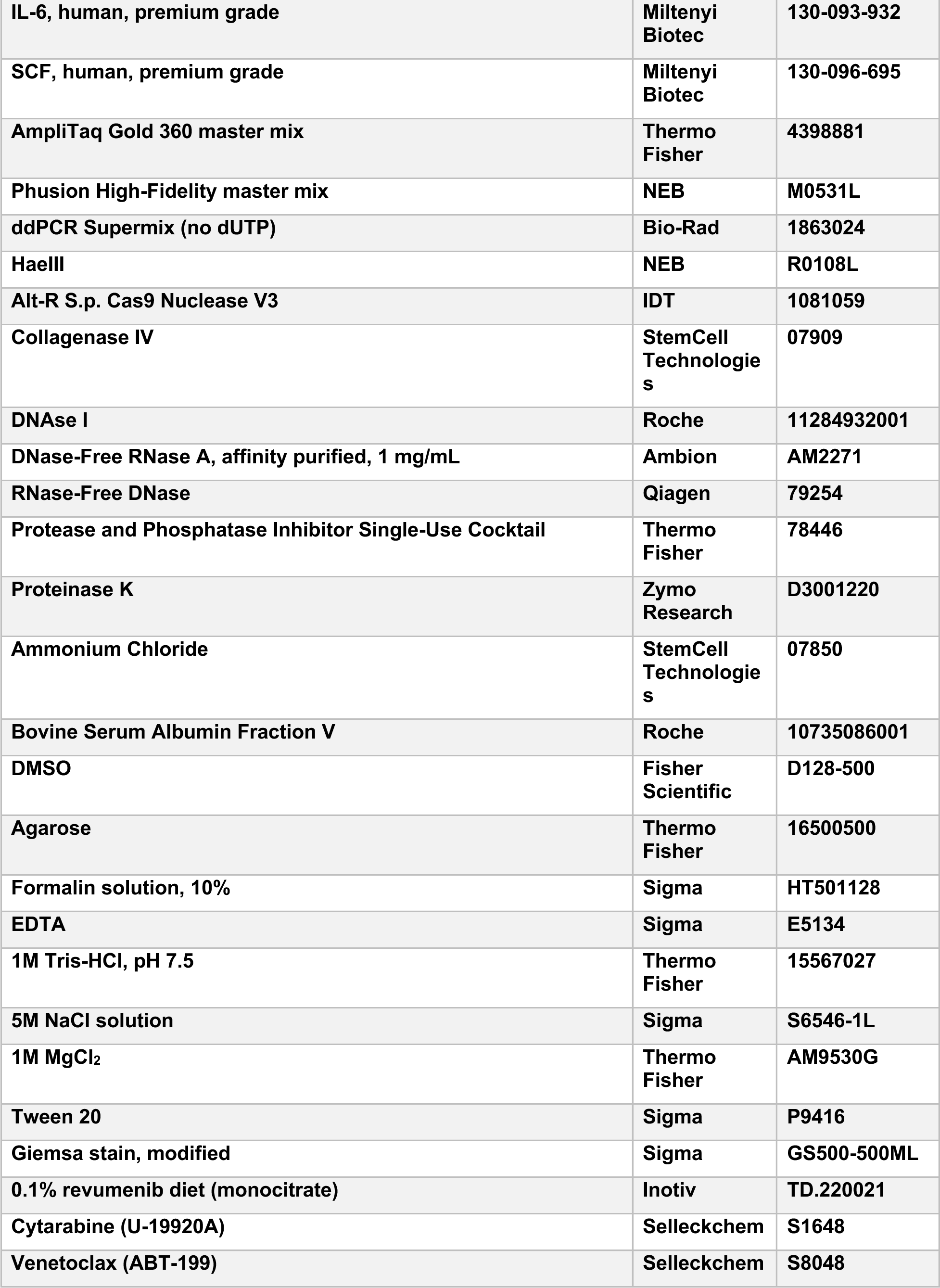
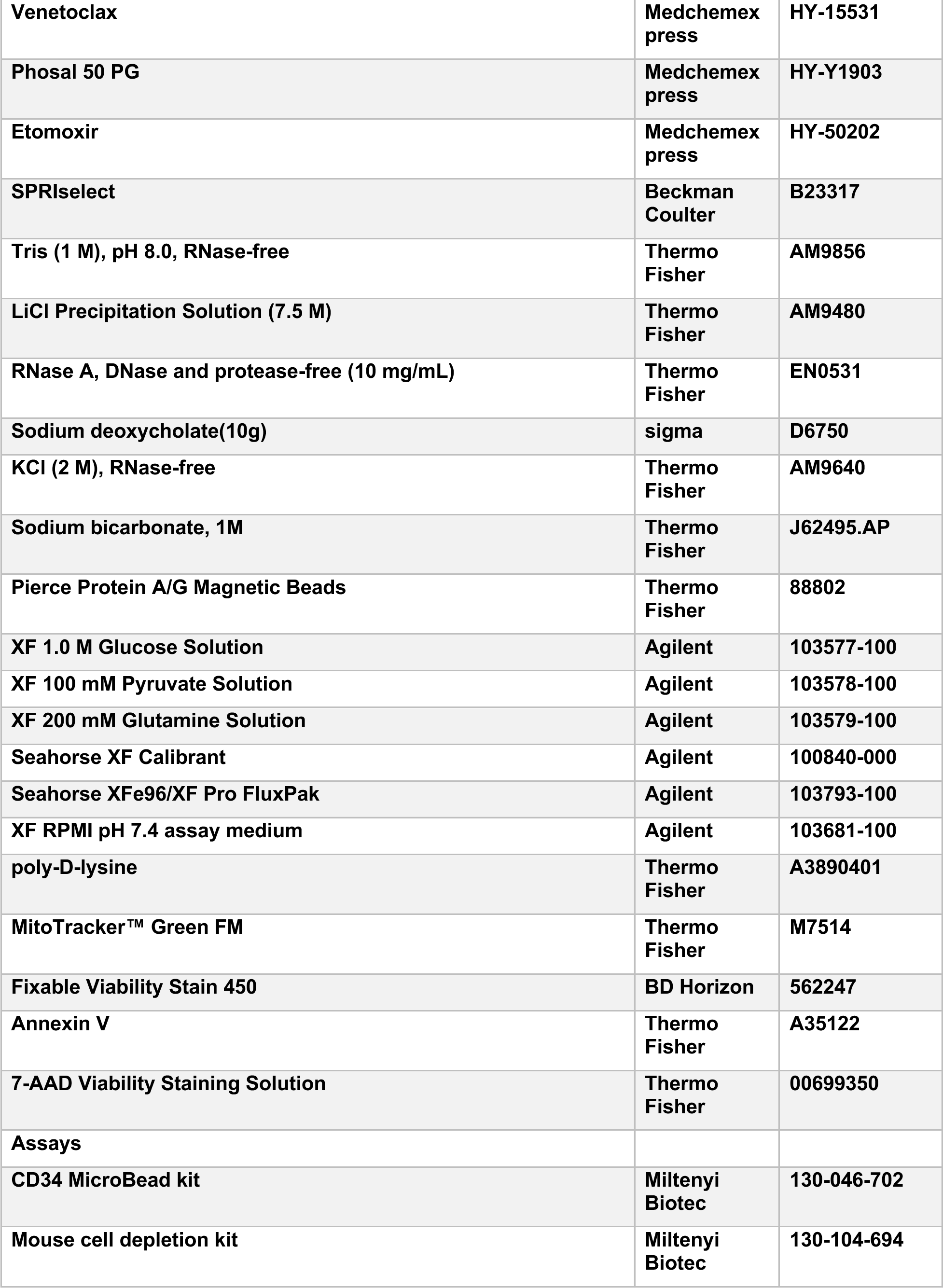
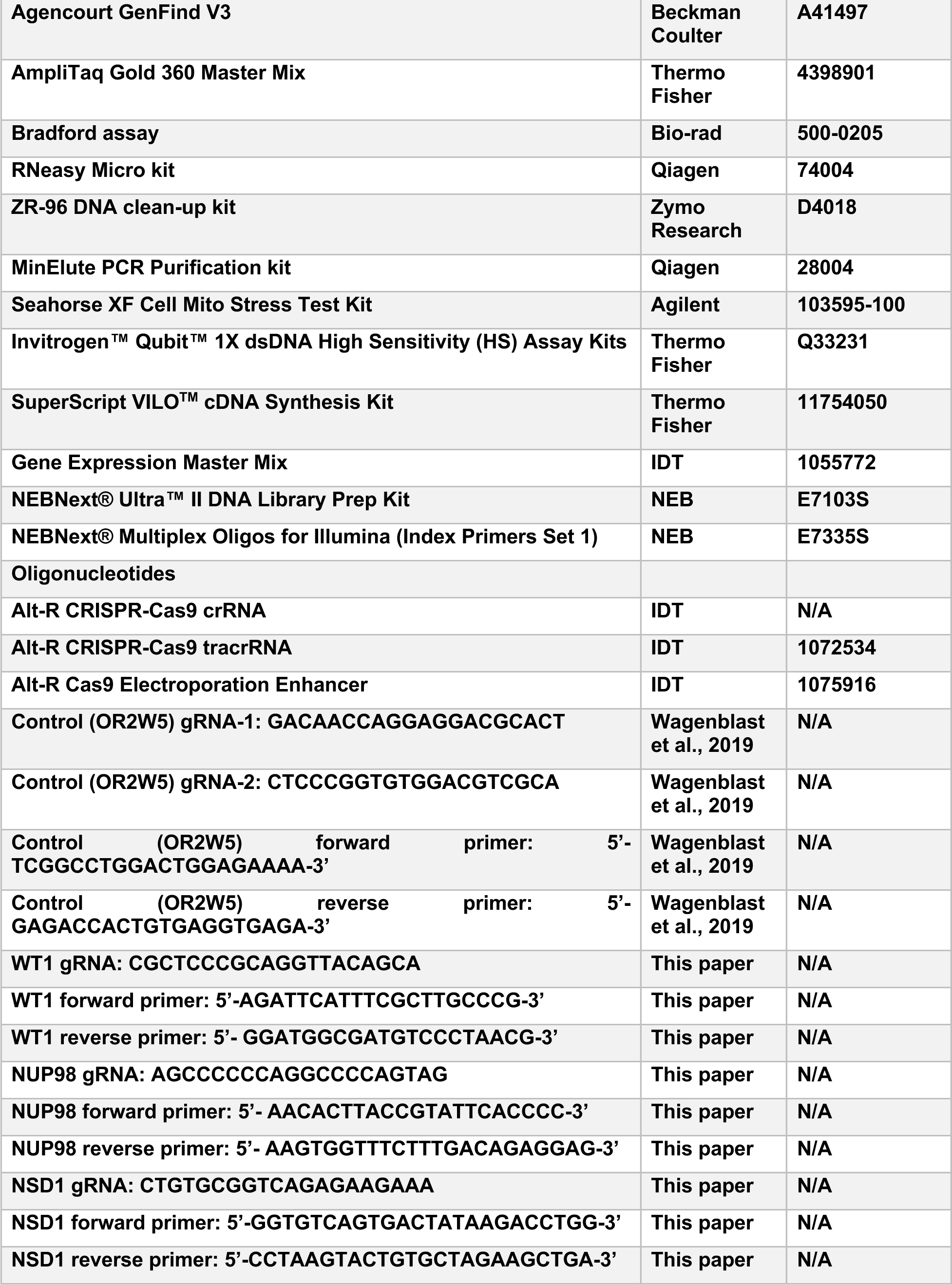
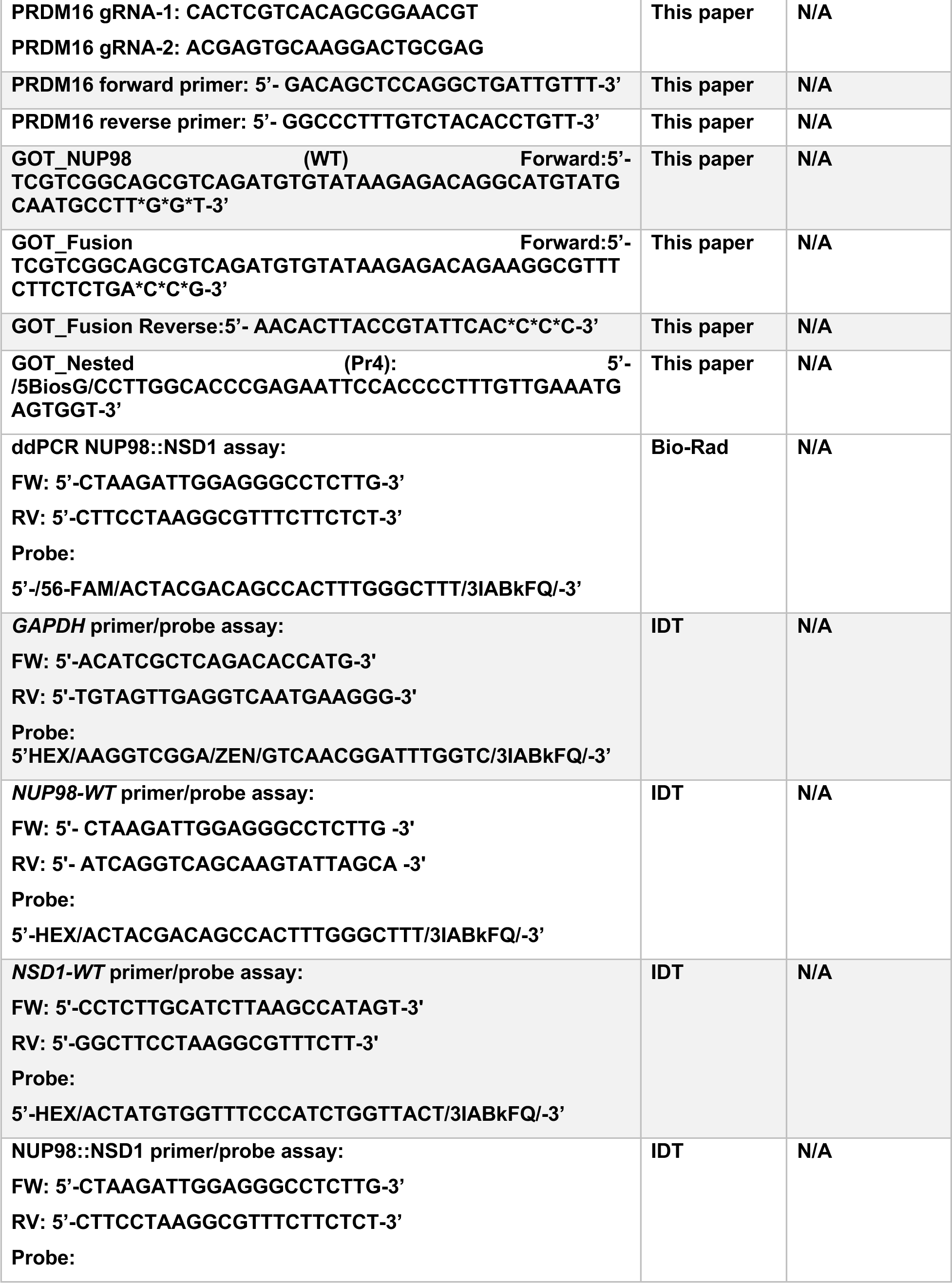
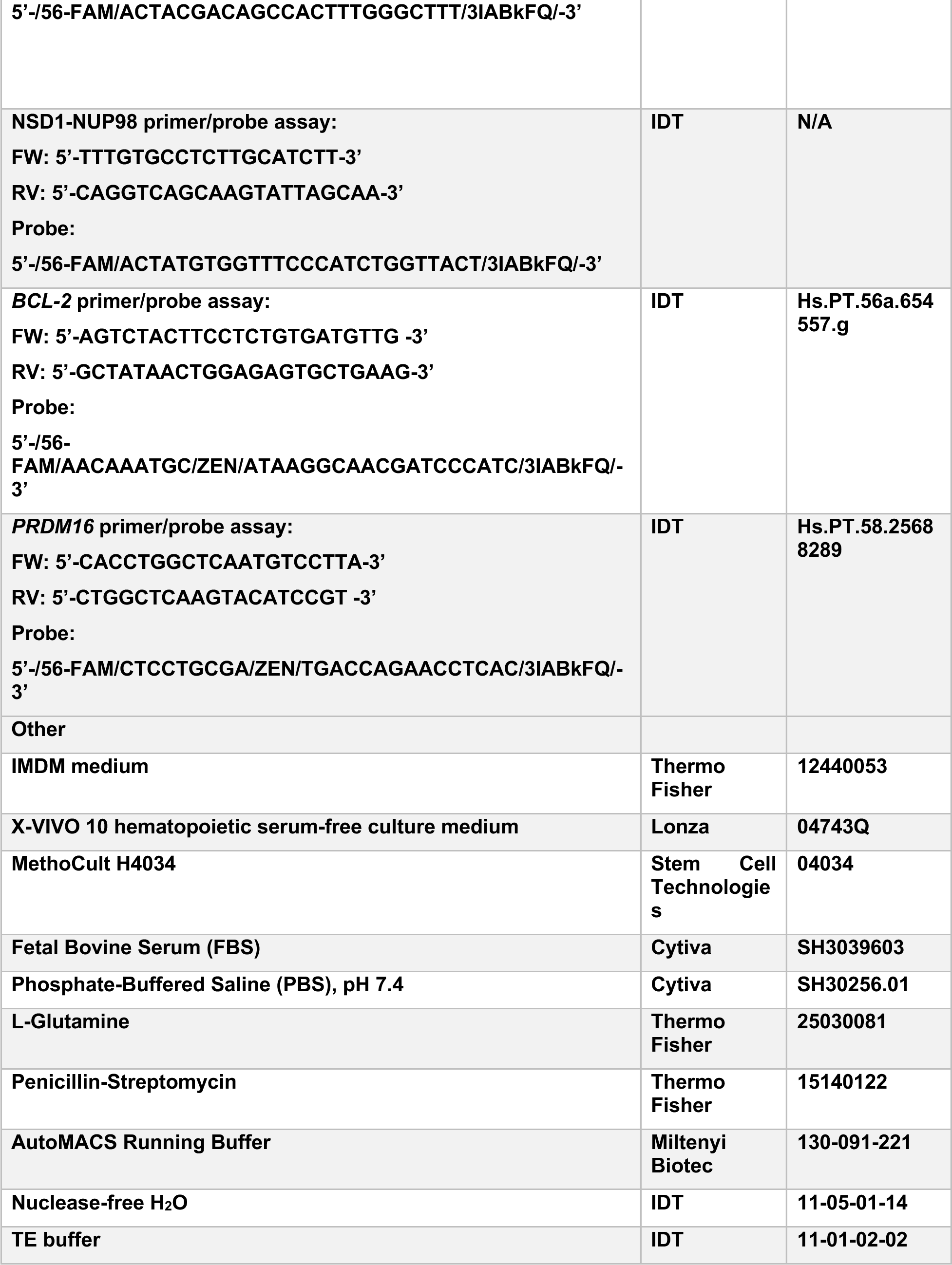
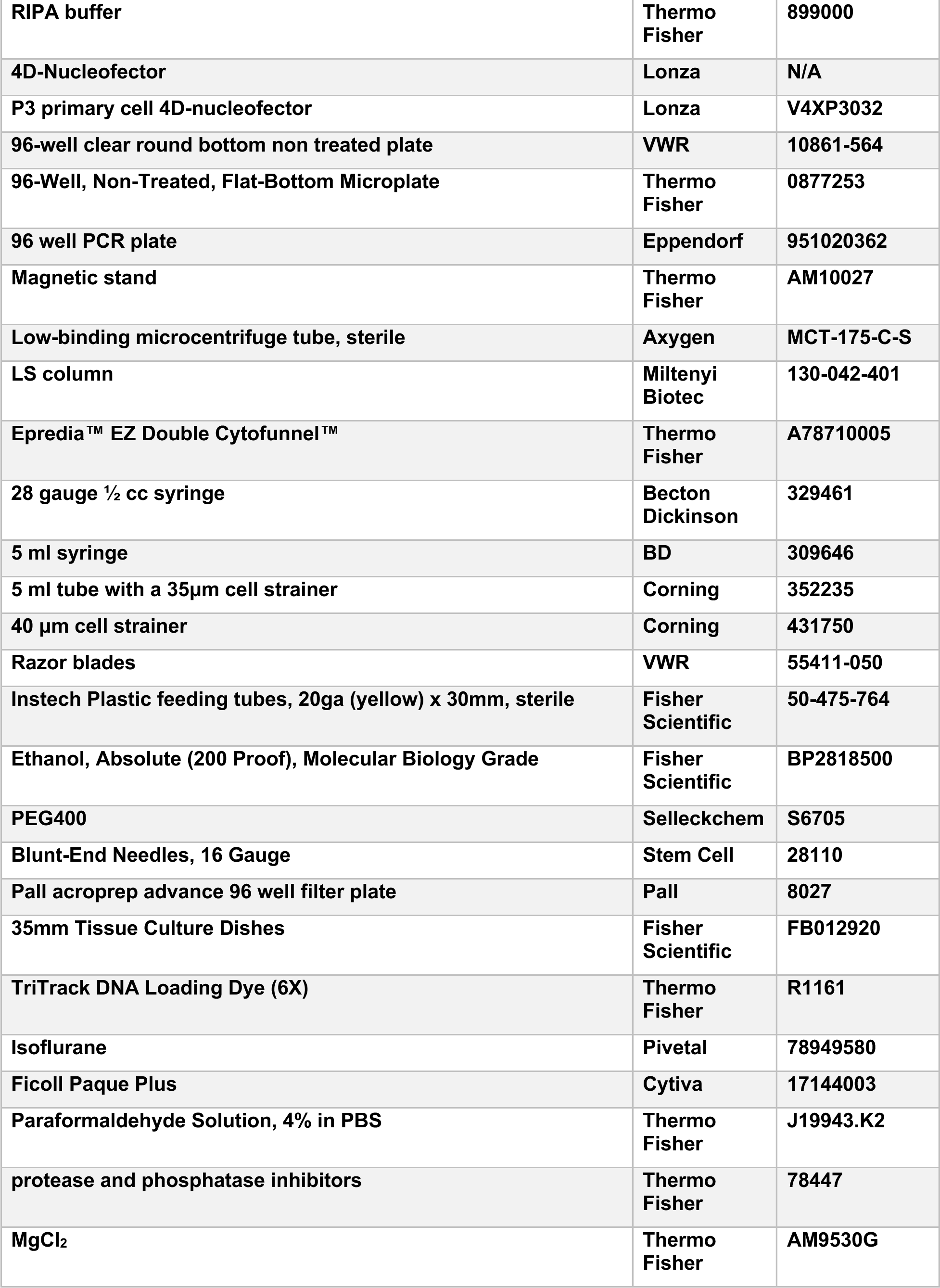
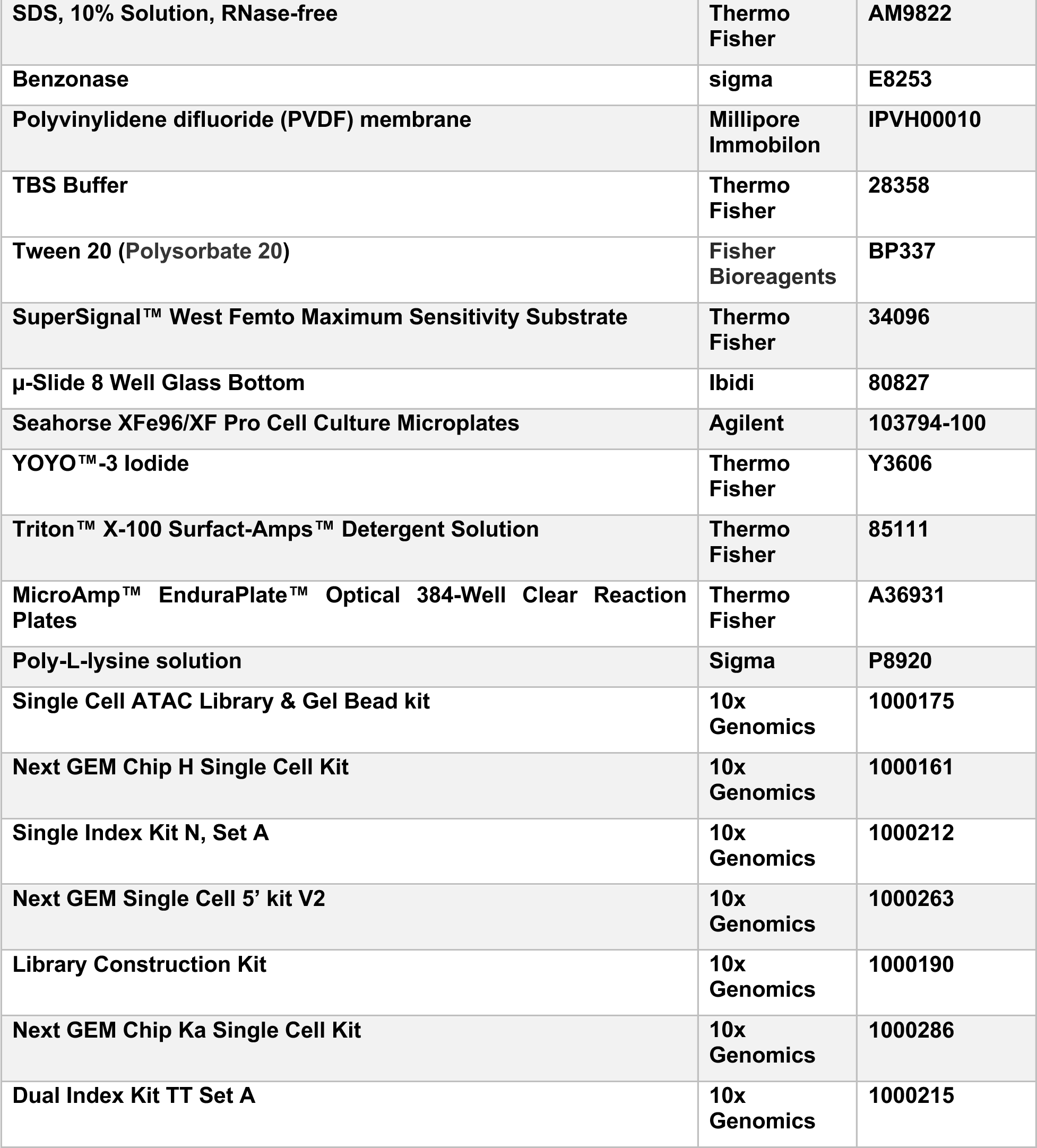

### EXPERIMENTAL MODEL AND STUDY PARTICIPANT DETAILS

#### Animal studies

All mouse experiments were approved by the Icahn School of Medicine at Mount Sinai Institutional Animal Care and Use Committee. We confirm that all experiments conform to the relevant regulatory and ethical standards. All xenotransplantations were performed in 8 to 12-week-old *NOD.Cg-Prkdc^scid^Il2rg^tm1Wjl^/SzJ* (NSG) mice (JAX) that were sublethally irradiated with 230 cGy, 24 hours before transplantation. Littermates were randomly assigned to experimental groups. Human cells were transplanted via intrafemoral injections.

#### Human patient samples

Human fetal liver samples were obtained from elective pregnancy terminations from the Birth Defects Research Laboratory (BDRL) at the University of Washington with informed consent following guidelines approved by the University of Washington and Icahn School of Medicine at Mount Sinai Institutional Review Board and from the Developmental Origins of Health and Disease (DOHaD) Biorepository at the Icahn School of Medicine at Mount Sinai with with informed consent following guidelines approved by the Icahn School of Medicine at Mount Sinai Institutional Review Board. Fetal liver samples were collected between 16 to 21 weeks’ gestation. Human umbilical cord blood samples were obtained from the Upstate New York Cord Blood Bank and Carolinas Cord Blood Bank with informed consent per guidelines approved by the Icahn School of Medicine at Mount Sinai Institutional Review Board. CD34+ enriched HSPCs were processed from pooled cord blood samples. Pediatric and adult bone marrow samples of CD34+ enriched HSPCs were purchased from Ossium Health. The AML PDX samples were obtained from Cincinnati Children’s Hospital, Children’s Hospital of Philadelphia and Mount Sinai Hospital. The genotype of PDX #1 is NUP98::NSD1 and FLT3-ITD, PDX #2 is NUP98::NSD1, FLT3-ITD and WT1 indel and PDX #3 is NUP98::NSD1, FLT3-ITD and WT1 indel.

#### Clinical cohort description

Patients with a NUP98 fusion were obtained from clinical trials AAML03P1, AAML0531, AAML1031, and CCG-2961 and used for analysis.^13^ NUP98::NSD1 (n = 104) made up most cases, followed by NUP98::KDM5A (n = 30) and NUP98 with other gene partners (n = 20). Consent, following the Declaration of Helsinki, was obtained from all study participants. The Fred Hutchinson Cancer Research Center Institutional Review Board and the COG Myeloid Biology Committee approved and oversaw the conduct of this study. The median (range) age of all NUP98 cases was 8.7 (0.4-19.9) years. For NUP98::NSD1, the median (range) age was 9.8 (1.2-19.9) years. For NUP98::KDM5A, the median (range) age was 2.8 (1.0-15.9) years. Patients with other NUP98 partners had a median (range) age of 7.9 (0.4-16.9) years.

#### Data availability

Raw sequence and processed data is available at Gene Expression Omnibus (GSE289849, GSE289851, GSE289855 and GSE289857). All other data is available in the manuscript or the supplementary material. For clinical cohort data, individual expression data files are available through CAVATICA in the Germline and Somatic Variants in Myeloid Malignancies in Children project. Access to protected files hosted on the Sequence Read Archive (SRA), such as raw sequencing data in bam or fastq format, is available through dbGaP TARGET: Acute Myeloid Leukemia study (accession: phs000465.v20.p8).

## METHOD DETAILS

### Fetal liver CD34+ HSPC isolation

The fetal liver sample was finely minced using razor blades and was then put into two 50 ml Falcon tubes, each with 40 ml of pre-warmed IMDM medium (Thermo Fisher), 5 ml of collagenase IV (Stem Cell Technologies), and 25 μl of 10 mg/ml DNAse I (Roche). The 50 ml tubes were incubated on a shaker at 37° C for 30 min. Following incubation, the dissociated sample was filtered through a 40 μm cell strainer (Thermo Fisher). The remaining tissue pieces were pushed through the strainer using the black rubber end of a 5 ml syringe (BD). The cells were then centrifuged at 350 x g for 10 min. The red blood cells were lysed with 5 ml ammonium chloride (StemCell Technologies) per 50 ml tube for 5 min at room temperature. Lysis was stopped with IMDM medium (Thermo Fisher), and cells were centrifuged at 350 x g for 10 min. The remaining cell pellets were combined and re-suspended in 500 μl MACS auto-running buffer (Miltenyi Biotec). Subsequently, CD34+ cells were subsequently enriched using the human CD34 MicroBead kit (Miltenyi Biotec) according to the manufacturer’s protocol. In total, 12 LS columns (Miltenyi Biotec) were used per fetal liver sample. CD34+ fetal liver cells were viably stored in a cryopreservation solution consisting of 90 % FBS (Cytiva) and 10 % DMSO (Sigma) in liquid nitrogen.

### Cord blood CD34+ HSPC isolation

Umbilical cord blood samples from multiple individuals were pooled and diluted with phosphate-buffered saline (PBS) in a 1:1 ratio. 12.5 ml of Ficoll-Paque plus lymphocyte separation media was aliquoted into 50 ml Falcon tubes. 37.5 ml of diluted blood was carefully overlayed into each 50 ml Falcon tube. Falcon tubes were centrifuged for 25 min at 350 x g without brake at 4° C. The “buffy coat” was collected in another 50 ml Falcon tube with 25 ml MEM medium and centrifuged for 10 minutes at 350 x g. The supernatant was removed and 5 ml ammonium chloride (StemCell Technologies) was added to each 50 ml tube, and incubated for 20 min at at 4° C. Cells were washed with 20 ml MEM medium, centrifuged and resuspended in 1 ml MACS auto-running buffer (Miltenyi Biotec). Magnetic labeling and CD34+ cell enrichment was performed as per manufacturer’s instructions (Miltenyi Biotec). CD34+ cord blood cells were viably stored in a cryopreservation solution consisting of 90 % FBS (Cytiva) and 10 % DMSO (Sigma) in liquid nitrogen.

### Primary CD34+ HSPCs

CD34+ enriched HSPCs were thawed via slow dropwise addition of thawing medium, made up of X-VIVO 10 medium (Lonza) with 50 % FBS (Cytiva) and DNase I (100 μg/ml, Roche). Cells were centrifuged at 350 x g for 10 min and then re-suspended in PBS (Cytiva) + 2.5 % FBS (Cytiva). CD34+ HSPCs were resuspended in 1000 μl PBS + 2.5 % FBS per 5×x10^6^ cells and stained for 60 min at 4° C. The following antibodies were used (volume per 1×x10^6^ cells): CD45 V500 (1:100, BD) and CD34 APC-Cy7 (1:200, BD). Afterwards, cells were washed and resuspended. The viability stain 450 (BD) was added (1:1000). CD34+ HSPCs were sorted using the FACSMelody (BD). Cell sorting purity checks were performed after each sorted sample (>95 %).

### CD34+ HSPCs *in vitro* culture

FACS-sorted CD34+ HSPCs were cultured in serum-free X-VIVO 10 medium (Lonza) supplemented with 1 % bovine serum albumin fraction V (Roche), 1×x L-glutamine (Thermo Fisher), 1×x penicillin-streptomycin (Thermo Fisher) and the following cytokines (Biotechne): FLT3 Ligand (100 ng/mL), G-CSF (10 ng/mL), SCF (100 ng/mL), TPO (15 ng/mL) and IL-6 (10 ng/mL). Cells were cultured in 96 well round-bottom plates (VWR).

### CRISPR/Cas9 RNP electroporation

Sorted CD34+ HSPCs were cultured for 48 hours in serum-free X-VIVO 10 medium (Lonza) as previously described.^61^ CRISPR/Cas9 RNP electroporations were performed using chemically synthesized gRNAs (IDT), recombinant Cas9 nuclease (IDT), and the 4D-Nucleofector (Lonza). For the preparation of the RNP complex, Alt-R CRISPR/Cas9 crRNA and tracrRNA (IDT) were reconstituted to a concentration of 200 μM in TE Buffer (IDT). The crRNA and tracrRNA were then mixed in a 1:1 ratio and annealed using a thermocycler at 95° C for 5 min, followed by cooling to room temperature. If two gRNAs were used simultaneously, such as for NUP98::NSD1, both cRNAs were annealed to the tracrRNA in a single tube at a 1:1 ratio. For each electroporation reaction, 1.2 μl of crRNA:tracrRNA complex, 1.7 μl of Cas9 protein, and 2.1 μl of PBS (Cytiva) were combined in a low-binding Eppendorf tube and incubated at room temperature for 15 min to form the complex. If multiple genes were targeted in the same electroporation reaction, such as for NUP98::NSD1/WT1ko, 1 μl of each crRNA:tracrRNA complex, 1.7 μl of Cas9 protein, and 1.3 μl of PBS were combined into the same Eppendorf tube. After incubation at room temperature for 10 min, 1 μl of 100 μM electroporation enhancer (IDT) was added to the mixture.

Pre-cultured cells were washed with pre-warmed PBS and centrifuged at 350 x g for 10 min. Approximately 2.5×x10^5^ cells were resuspended in 20 μl of Buffer P3 (Lonza) per reaction and added to the tube containing the CRISPR/Cas9 gRNA RNP complex. The mixture was thoroughly mixed by pipetting up and down and then transferred to the electroporation chamber (Lonza).

The cells were electroporated using the 4D-Nucleofector with the program DZ-100. Immediately following electroporation, 180 μl of pre-warmed X-VIVO 10 medium (as described above) was added to the electroporation chamber, and cells were transferred to a 96-well round-bottom plate. The electroporated cells were allowed to recover overnight at 37° C in a tissue culture incubator. For each CRISPR/Cas9 electroporation, a small subset of cells was cultured for 5-7 days to validate CRISPR/Cas9 efficiency.

### gRNA sequences

The gRNA sequences for NUP98 and NSD1 were designed based on naturally occurring break sites.^14^ The WT1 gRNAs were predicted using the CRoatan algorithm^62^, and the PRDM16 gRNAs were designed using Benchling. For control, two gRNAs targeting the olfactory receptor OR2W5 were used.

### Real-time quantitative RT-PCR

For NUP98::NSD1 expression quantification, NUP98::NSD1 or NUP98::NSD1/WT1ko leukemic cells were cultured in X-VIVO 10 medium. 5×x10^5^ cells were collected, total mRNA was isolated, and cDNA was generated. qPCR was performed on the QuantStudio 7 Pro (Thermo Fisher) using NUP98::NSD1 fusion gene primer/probe sets. For *BCL2* mRNA expression quantification, NUP98::NSD1/WT1ko leukemic cells were cultured, and *PRDM16* was knocked out. After 10 days, total mRNA was isolated, and cDNA was generated. Then, qPCR was performed on the QuantStudio 7 Pro (Thermo Fisher) using *BCL2* primer/probe sets. All signals were quantified using the ΔΔCt method and were normalized to the levels of *GAPDH*.

### Chromosomal rearrangement efficiency by droplet digital PCR

To determine the NUP98::NSD1 chromosome rearrangement efficiency upon CRISPR/Cas9 editing, droplet digital PCR (ddPCR, Bio-Rad) was carried out. A FAM primer/probe set detecting a 117 bp NUP98::NSD1 fusion and a HEX primer/probe set detecting a 160 bp wild-type NUP98 genomic region were designed. Genomic DNA was isolated from CRISPR/Cas9-edited HSPCs and subjected to ddPCR with 1 μl of genomic DNA (15 and 30 ng/μl), and no HaeIII was used. The ddPCR program was: 95° C for 10 min, followed by 94° C for 30 s, 60° C for 1 min (40 cycles), 98° C for 10 min and then 4° C hold.

### Karyotyping analysis

CRISPR/Cas9-edited NUP98::NSD1 and NUP98::NSD1/WT1ko leukemic cells were cultured and expanded in serum-free X-VIVO 10 medium (Lonza) supplemented with 1 % bovine serum albumin fraction V (Roche), 1×x L-glutamine (Thermo Fisher), 1×x penicillin-streptomycin (Thermo Fisher) and the following cytokines (Biotechne): FLT3 Ligand (100 ng/mL), G-CSF (10 ng/mL), SCF (100 ng/mL), TPO (15 ng/mL) and IL-6 (10 ng/mL). The karyotyping of chromosomes was prepared according to standard procedures. Briefly, metaphase slides were prepared, banded with trypsin, and stained with Leishman’s stain. G-banded slides were scanned, and metaphases were analyzed using the imaging system Ikaros (Meta Systems). For each condition, 20 metaphases were analyzed by G-banded karyotyping to identify numerical and structural chromosomal abnormalities. Karyotyping analyses were routinely completed for FL NUP98::NSD1, FL NUP98::NSD1/WT1ko and CB NUP98::NSD1/WT1ko leukemic cells.

### Western blot assay

NUP98::NSD1 and NUP98::NSD1/WT1ko leukemic cells expanded from secondary xenograft were used. 1×x10^6^ cells were lysed in cold RIPA buffer (Thermo Fisher) with protease and phosphatase inhibitors (Thermo Fisher), 10 U/ml Benzonase (Sigma), and 2 mM MgCl2. Then samples were centrifuged at 13,000 x g for 15 min at 4° C and the supernatants were subsequently used. Protein concentration was determined by Bradford assay (Bio-Rad) according to the manufacturer’s instructions. 20 μg of lysate was resolved on a 10% SDS-PAGE gel and transferred to a polyvinylidene difluoride (PVDF) membrane (Millipore Immobilon). Blocking and antibody dilutions were performed in 5% nonfat dry milk (NFDM) dissolved in TBS (Thermo Fisher) with 0.1% Tween-20 (Fisher Bioreagents). Blocking and secondary antibodies were incubated for 1 hour at room temperature with gentle rocking. Primary antibodies were incubated overnight at 4° C with gentle rocking. Blots were visualized with Supersignal West Femto Chemiluminescent Substrate (Thermo Fisher). Antibodies used for these experiments are: WT1 (Novus Bio, NB110-60011, 1:1000), GAPDH (Abcam, ab8245, 1:20,000) and goat anti-mouse IgG HRP conjugate (Thermo Scientific, 32430, 1:20,000).

### Methylcellulose colony formation assay

For methylcellulose colony formation assays, CD34+ HSPCs sorted from human fetal liver, postnatal cord blood, pediatric bone marrow, and adult bone marrow were cultured in X-VIVO-10 medium (as described above). After 48 hours of culture, the cells proceeded to CRISPR/Cas9 RNP electroporation as described. Two days post-electroporation, cells were collected and counted, and a specific number of cells were resuspended in MethoCult H4034 optimum methylcellulose medium (StemCell Technologies) and plated in 35 mm cell culture dishes. These cells were labeled as day 1 cells. After 12-14 days, individual colonies were picked and genotyped to analyze the CRISPR/Cas9-edited loci. Cells were maintained in liquid culture, with media changed or cells split every other day to ensure high viability. At subsequent time points, including day 7, day 14, day 21, and day 28, cells were plated again on methylcellulose medium as described for day 1. 96 individual colonies per condition were picked, washed with PBS, and stored in a 96-well PCR plate (Eppendorf) at -80° C. For genotyping, genomic DNA isolation and PCR amplification were performed. Genomic DNA was isolated from colonies using the Agencourt GenFind V3 kit (Beckman Coulter). The isolation process was performed in 96-well PCR plates (Eppendorf) using a magnetic stand (Thermo Fisher), following the manufacturer’s protocol with the following volume modifications: 100 μl lysis buffer, 2.3 μl of proteinase K, 75 μl magnetic particles, 200 μl wash buffer 1, 125 μl wash buffer 2 and 60 μl TE buffer (IDT) for elution. PCR was used to amplify the CRISPR/Cas9-modified genomic locus. Each PCR reaction contained 23 μl of genomic DNA, 1 μl of forward and reverse primer (10 μM, IDT), and 25 μl of AmpliTaq Gold 360 master mix (Thermo Fisher). The PCR program was: 95° C for 10 min, followed by 95° C for 30 s, 56° C for 30 s and 72° C for 1 min (40 cycles), and then 72° C for 7 min. For the NUP98::NSD1 genotype, the fusion gene was identified by a 540 bp product in fusion-positive samples. For control-(OR2W5), expected deletions were confirmed through gel electrophoresis of the PCR products. For WT1ko, PCR products were purified, and Sanger sequencing was performed using the reverse PCR primer. Chromatographs were analyzed using TIDE^63^ to verify editing efficiencies. To further analyze the immunophenotypes of the colonies derived from methylcellulose assays, remaining cells were collected and pooled by adding 3 mL of PBS into each 35 mm dish and washed three times with PBS with 2.5% FBS. The collected cells were then stained with antibodies, and their immunophenotypes were analyzed by flow cytometry.

### Immunofluorescence

To evaluate biomolecular condensate formation in NUP98::NSD1-rearranged cells, NUP98::NSD1 and NUP98::NSD1/WT1ko leukemic cells expanded from secondary xenografts were used. μ-Slide Glass wells (ibidi) were coated with 200 μL of Poly-L-Lysin (Sigma) for 30 min at room temperature. Appropriate cells were washed once with PBS and resuspended in PBS. Poly-L-Lysin was removed from the well and 2,000 - 3,000 cells were added per well. After spinning at 200 x g for 10 min, the supernatant was carefully removed. Cells were fixed with 4% PFA for 10 min at room temperature, washed three times with PBS, permeabilized with PBS + 0.5% Triton for 15 minutes at room temperature, and washed again with PBS. Blocking solution (PBS + 10% FBS + 5% BSA) was added for 30 minutes at room temperature, followed by incubation with the primary antibody (NUP98 (2H10), Santa Cruz, diluted in blocking solution 1:50) overnight at 4° C. After washing with PBS + 0.025% Tween, cells were incubated with a secondary antibody (diluted in PBS + 0.025% Tween 1:50) for 1.5 hours at room temperature in the dark, washed again, and stained with DAPI (1 μg/mL) for 5 minutes at room temperature. Following three final PBS washes, 200 μl PBS was added to the wells. Cells were visualized using a Zeiss LSM880 Airyscan microscope.

### Primary xenotransplantation

The electroporated cells were allowed to recover overnight at 37° C in a tissue culture incubator. Then, cells were transplanted into sublethally irradiated NSG mice via intrafemoral injections. Mice were anesthetized with isoflurane, and their left knee was secured in a bent position. A 27-gauge needle was used to drill a hole into the left femur, followed by the injection of CRISPR/Cas9-edited HSPCs in 30 μl of PBS using a 28-gauge ½ cc syringe (BD). Mice from the primary xenotransplantation of CRISPR/Cas9-edited HSPCs were euthanized at week 16. Engraftment was considered positive if the human CD45+ engraftment level in the bone marrow was greater than 1%, with the population presenting as dense clusters rather than dispersed signals. To ensure data robustness, only mice with confirmed CRISPR/Cas9 edits (>90% efficiency in control and WT1ko groups, and NUP98::NSD1 fusion-positive status) were included in downstream analyses.

### Secondary transplantation and limiting dilution assay

For secondary transplantation, cells harvested from primary xenografts (bone marrow) that had been validated for CRISPR/Cas9 edits were thawed as described in the fetal liver sorting protocol. Multiple xenograft samples were pooled, and murine cells were depleted using the mouse cell depletion kit (Miltenyi Biotec) and processed according to the manufacturer’s instructions. In the limiting dilution assay (LDA), cells were counted and serially diluted into specific cell doses for transplantation. Typically, four different cell doses were prepared, with five mice assigned to each dose group. The engraftment frequency was calculated based on the presence or absence of human CD45+ cells in the bone marrow of recipient mice after 10 weeks. Engraftment was considered positive if the human CD45+ cells exceeded 1% in the bone marrow and appeared as concentrated clusters rather than dispersed populations. ELDA online software (https://bioinf.wehi.edu.au/software/Elda/) was used to estimate the frequency of stem cells or leukemia stem cells.

To evaluate the self-renewal capacity of LSC and non-LSCs in our model. LSCs (CD34+CD117+) and non-LSCs (non-CD34+CD117+) were sorted from NUP98::NSD1 or NUP98::NSD1/WT1ko xenografts after being depleted of murine cells using the mouse cell depletion kit (Miltenyi Biotec). Cells from each condition were counted and diluted into specific cell doses for transplantation. The prepared cells were then transplanted intrafemorally into NSG mice. Mice were euthanized at week 10, and engraftment was evaluated. Engraftment was considered positive if the human CD45+ cells exceeded 1% in the bone marrow and appeared as concentrated clusters rather than dispersed populations. Similarly, to evaluate the self-renewal capacity of CD19+ and non-CD19+cells in our model, cells were sorted from NUP98::NSD1/WT1ko xenografts following mouse cell depletion. These sorted cells were transplanted intrafemorally into NSG mice. After 10 weeks, mice were euthanized, and engraftment and leukemia stem cell frequencies were assessed as described above.

### Patient-derived xenograft models of AML

NUP98::NSD1 with or without WT1 mutation PDX cells were thawed carefully following the protocol described before. Murine cells were depleted using the mouse cell depletion kit (Miltenyi Biotec). The cells were then spun down, resuspended in PBS, and counted. The prepared cells were transplanted into NSG mice using the same method as above. Depending on the timing outlined in the *in vivo* treatment design, the mice were euthanized between 8 - 14 weeks. The PDX samples from Cincinnati Children’s Hospital and the Mount Sinai Hospital were used for scRNAseq studies, the PDX sample from Children’s Hospital of Philadelphia was used for *in vivo* drug studies.

### Genotyping of xenotransplantations

To assess the efficiency of each xenograft, genomic DNA was isolated from bulk bone marrow cells using the Agencourt GenFind V3 kit (Beckman Coulter), following the same procedure as previously described. PCR and resulting analyses were performed using the same protocol as previously described for assessing editing efficiencies. For PRDM16ko-edited xenografts, expected deletions were confirmed through gel electrophoresis of the PCR products. Mice included in the final analysis met the following criteria: CRISPR/Cas9 editing efficiency >90%, as determined by PCR or Sanger sequencing. CD45+ engraftment level > 1% in the bone marrow, with the population presenting as dense clusters rather than dispersed signals. These exclusion criteria were determined and pre-specified before analysis.

### Flow cytometry

For xenografted cells, the bone marrow, including left and right femurs, of each NSG mouse was flushed with 1 ml staining buffer (PBS + 2.5 % FBS). Collected cells were centrifuged at 350 x g for 10 min and resuspended in 500 μl staining buffer. For antibody staining, 50 ul of cells per sample were mixed with 50 μl of antibody mix and incubated for 60 minutes at 4° C, protected from light. After staining, cells were washed once with staining buffer and resuspended in 250 μl staining buffer. In addition, 25 μl of each sample were transferred to a 96-well PCR plate (Eppendorf) and stored at -80°C for genotyping. The remaining cells were used for flow cytometry analysis. The following antibodies were used (all from BD, unless stated otherwise): CD45 V500 (1:100), CD33 BV711 (1:100), CD19 PE-Cy7 (1:100), CD41 PE-Cy5 (1:200, Beckman Coulter), GlyA PE (1:100, Beckman Coulter), CD71 BV650 (1:100), CD3 FITC (1:100), CD34 APC-Cy7 (1:100), CD11b PC5 (1:100, Beckman Coulter), CD13 PE (1:100), CD14 BV605 (1:100), CD33 BV786 (1:100), fixable Viability Stain 450 (1:1000). A pan-lineage panel was designed and used for primary xenograft analysis including: CD45, CD19, CD3, CD33, CD41, GlyA, CD71, CD34 and CD117. A myeloid panel was designed and used for secondary and serial xenograft analysis including CD45, CD19, CD33, CD11b, CD13, CD14, CD34 and CD117. Flow cytometry was performed using the FACSFortessa (BD Biosciences). The remaining unstained cells from each xenografted mouse were viably frozen and stored in liquid nitrogen. For *in vitro* cultured cells, at least 100,000 cells were collected and stained for each experiment. The cells were stained for 60 minutes at 4° C, protected from light. Cells were then washed once with staining buffer and resuspended in 250 μl of staining buffer. Flow cytometry analysis was performed using the FACSFortessa (BD) or Attune (Thermo Fisher). All data were analyzed with FlowJo.

### Cellular morphology

The secondary xenograft was used for cell morphology study. Murine cells were depleted using the mouse cell depletion kit (Miltenyi Biotec). Cells were stained for 15 min at 4° C and then processed through an LS column (Miltenyi Biotec) to isolate human cells. Human cells were cytospun onto glass slides using the EZ Double Cytofunnel (Thermo Fisher) and the CytoSpin 4 instrument (Thermo Fisher) at 112 x g for 10 min with medium acceleration. Slides were air-dried overnight, fixed with methanol for 5 min, and then stained with standard Giemsa staining. For morphological analysis, blasts were defined using the basic morphological characteristics established for human diseases, including high nuclear/cytoplasmic ratio, easily visible nucleoli, and usually, but not invariably, fine nuclear chromatin. All quantifications were conducted in a blinded manner to avoid bias. Cells were visualized using a Zeiss Axio Imager Z2(M) microscope equipped with a 63×x oil objective.

### ScRNA-seq and scATAC-seq library generation

Xenografted cells were thawed following the previously described protocol. Human cells were isolated using mouse depletion kit (Miltenyi Biotec), then cells were stained with CD45 V500 (1:100) and CD34 APC-Cy7 (1:100) in staining buffer and incubated for 60 minutes at 4° C, protected from light. Fixable Viability Stain 450 (1:1000) was added 10 minutes before sorting and viable human CD45+ cells were sorted using FACSMelody (BD). On average, 3-5×x10^5^ cells per sample was submitted for scRNA-seq and scATAC-seq library generation.

Viability of single cells was assessed using Acridine Orange/Propidium Iodide viability staining reagent (Nexcelom), and debris-free suspensions of >80% viability were deemed suitable for experiments. The Chromium 10×x Genomics 5’ GEX v2 protocol was used for scRNA-seq. Briefly, Gel-Bead in Emulsions (GEMs) were generated on the sample chip using the Chromium X system. Barcoded cDNA was extracted from the GEMs following post-GEM RT cleanup and amplified for 12 cycles. For GEX library preparation, amplified cDNA was fragmented and subjected to end-repair, poly A-tailing, adapter ligation, and 10×x-specific sample indexing, following the manufacturer’s protocol. The GEX libraries were quantified using TapeStation (Agilent) and QuBit (ThermoFisher) and sequenced in paired-end mode on a NovaSeq 6000 instrument (Illumina), targeting a depth of 25,000 reads per cell. Raw Fastq files were aligned to the GRCh38 reference genome (2020-A) using Cell Ranger v5.0.1 (10×x Genomics).

scATAC-seq was performed using the 10×x Genomics Chromium Next GEM Single Cell ATAC Reagent Kit v1.1, following the manufacturer’s protocol. Briefly, cells were washed and resuspended in nuclei lysis buffer (10 mM Tris-HCl [pH 7.4], 10 mM NaCl, 3 mM MgCl₂, 0.1% Tween-20, 0.1% Nonidet P40 substitute, 0.01% digitonin, 1% BSA; 10×x Genomics) and incubated on ice for 4-5 min, followed by the addition of chilled wash buffer (10 mM Tris-HCl [pH 7.4], 10 mM NaCl, 3 mM MgCl₂, 1% BSA, 0.1% Tween-20; 10×x Genomics). Nuclei were pelleted by centrifugation, resuspended in 1×x diluted nucleus buffer (10×x Genomics), and counted using the Acridine Orange/Propidium Iodide viability staining reagent on a Cellometer Auto2000 automated cell counter (Nexcelom). The nuclei suspensions were incubated in a Transposition Mix (10×x Genomics), which simultaneously cuts accessible chromatin regions and inserts a short DNA adapter sequence to the end of the DNA fragments. Gel beads-in-emulation (GEMs) were subsequently generated by combining barcoded gel beads, transposed nuclei, a master mix, and partitioning oil on a Chromium Next GEM Chip H, using Chromium X (10×x Genomics). Upon GEM generation, a barcoding process was achieved within the droplets where (i) an Illumina P5 sequence, (ii) a 16 nt 10×x Barcode and (iii) a Read 1 (Read 1N) sequence were attached to the DNA fragments via PCR reactions. The transposed and amplified DNA fragments were cleaned up using SPRIselect and processed to incorporate indexing primers for library preparation. The libraries were sequenced in a paired-end fashion on a NovaSeq 6000 sequencer (Illumina) at 25,000 reads per nucleus with the following parameter: 50/8/16/50.

### ScRNA-seq data quality control and pre-processing

Sequenced FASTQ files were aligned, filtered, barcoded, and unique molecular identifier (UMI) counted using Cell Ranger Chromium Single Cell RNA-seq (v 7.1.0 and v5.0.1) (10X Genomics) with the hg38 human genome reference (version 2020-A; RRID: SCR_017344). Cells were filtered to retain those with ≥1000 UMIs, ≥400 expressed genes, and <15% of reads mapping to the mitochondrial genome. UMI counts were then normalized to a total of 10,000 UMIs per cell and log-transformed with a pseudocount of 1 using the “LogNormalize” function in the Seurat package (version 4.0.3; RRID: SCR_016341).^64,65^ The top 2,000 highly variable genes were identified with the “vst” method in the “FindVariableFeatures” function, and the data were scaled using the “ScaleData” function. Cell cycle scoring was performed using the “CellCycleScoring” function, which assigns cell cycle phase scores (G1, S, and G2/M) based on canonical marker gene expression.

### ScRNA-seq data dimensionality reduction and integration

Principal component analysis was performed using the top 2,000 highly variable features (“RunPCA” function) and the top 30 principal components were used in the downstream analysis. K-Nearest Neighbor graphs were obtained using the “FindNeighbors” function whereas the UMAPs were obtained using the “RunUMAP” function. The Louvain algorithm was used to cluster cells based on expression similarity. The top cluster biomarkers were identified by “FindAllMarkers” function. Cell density estimations were performed using the stat_density_2d function of the ggplot2 (version 3.3.5) package. AUCell (v1.16.0) was used to evaluate gene set activity by calculating the Area Under the Curve (AUC) for each gene set in individual cells, providing a measure of enrichment based on ranked gene expression.^44^ A Wilcoxon rank-sum test was performed to assess differences in AUCell scores between two conditions and violin plots were generated with ggplot2. The cells from cord blood and leukemia patient samples were projected into the xenograft UMAP using MapQuery function in Seurat. Gene regulatory network analysis and cell-state-specific regulon activity inference were performed using pySCENIC.^44^

### ScRNA-seq data cell type annotation and differential biomarker analysis

To annotate clusters with relevant cell types, the sc-transcriptomes were projected against a hematopoietic reference map using BoneMarrowMap R package.^22^ To visualize the contribution of each annotated cell type across genotypes, the percentage of cells in each cell type within each genotype was represented with barplots. Differential comparisons in cell-type abundance between conditions were done using the miloR (v1.2.0) package.^66^ For the FL and CB sample groups, pseudo-bulk expression profiles were generated from scRNA-seq data by summing raw counts across cells within each cluster. The aggregated counts were normalized using the DESeq2 pipeline to perform downstream differential expression analysis and box plots were generated using the normalized counts with ggplot2. For cytarabine, revumenib and PRDM16ko samples, differential markers for each cluster were identified using the Wilcox test (“FindMarkers” function) with adjusted p-value < 0.01, absolute log2 fold change > 0.25, and minimum 10 % of cells expressing the gene in both comparison groups using 1,000 random cells to represent each cluster. The differentially expressed genes were then used to perform overrepresentation analysis (ORA) with enricher function from the clusterProfiler package using the collections from the Molecular Signatures Database (MSigDB). The enriched pathways were visualized using barplots. UMAPs of HOX cluster genes expression was calculated based on a defined expression module including HOXA9, HOXA10, HOXA11, HOXA11-AS, HOXB2, HOXB3, HOXB-AS3, HOXB4, HOXB5, HOXB6, HOXB7, HOXB8 and HOXB9.

### Velocity and pseudotime analysis

RNA velocity analysis was performed using the scvelo Python package (version 0.2.5, RRID: SCR_018168).^67^ The unspliced and spliced count matrices are generated from Cell Ranger outputs using the run10×x function from the velocyto package (version 0.17.17, RRID: SCR_018167).^26^ For each sample and integrated dataset, the Seurat object is converted into anndata (version 0.8.0, RRID: SCR_018209)^68^ and merged with the unspliced and spliced count matrices using scanpy (v1.9.3, RRID: SCR_018139)^69^. Moments for velocity estimation were calculated using 30 principal components and 30 neighbors. RNA velocity was estimated using stochastic model with default parameters using scvelo. Cell type labels from the integrated dataset were projected to visualize the velocity streams. Diffusion pseudotime (DPT) was calculated using the destiny R package^27^, which computes pseudotime based on transition probabilities. Cells were ordered along pseudotime, and results were visualized with violin plots and UMAP projections.

### ScATAC-Seq data processing and motif analysis

Sequenced FASTQ files were aligned, filtered, barcoded and UMI counted using Cell Ranger ARC version 2.0.0, by 10X Genomics, with the hg38 human genome reference. Peaks were called using ‘CallPeaks’, the MACS2 wrapper function, in Signac (v1.6.0, RRID:SCR_021158).^70^ Cells were filtered to retain only those with total number of fragments in peaks > 1,000 and < 50,000, percentage of reads aligned in peak regions > 20%, blacklist ratio < 0.002, nucleosome signal < 1.5 and the TSS enrichment score > 2. Blacklist ratio is calculated as the number of reads aligned to blacklist region divided by the total number of reads, nucleosome signal was calculated by ‘NucleosomeSignal’ function in Signac and TSS enrichment score was calculated by ‘TSSEnrichment’. Latent semantic indexing (LSI) was then applied to the top 95% of peaks in terms of their variability, which combines the term frequency-inverse document frequency (TFIDF) used for normalization and singular-value decomposition (SVD) used for dimensional reduction. The first LSI component was excluded from all downstream analysis as it highly correlated with sequencing depth. De-novo clustering was performed by using ‘FindNeighbors’ and FindClusters functions in Signac. Motif position frequency matrices were retrieved from JASPAR 2020 database (RRID: SCR_003030) and was added to the scATAC data in Signac.^71^ ChromVAR motif analysis was performed using the RunChromVAR function in Signac, and results of selected transcription factor were visualized on UMAP using FeaturePlot function.^23^ Gene activity scores were computed using the GeneActivity function of Signac by aggregating the accessibility of genomic regions within a gene body and promoter (2 kb upstream of the transcription start site).

### Genotyping of targeted loci with chromatin accessibility (GoT-ChA) library preparation

Primary FL- and CB-derived CD34+ HSPCs were sorted and cultured in X-VIVO 10 medium and supplied with growth factors as described previously. After 48 hours, CRISPR/Cas9 RNP-based electroporation was carried out to generate NUP98::NSD1/WT1ko. Then, cells were cultured and expanded for 28 days. On day 28, cells were harvested and washed with PBS once. Cells were subjected to GoT-ChA.^19^ Briefly, the nuclei isolation procedure is the same as scATAC-seq, and the isolated nuclei were processed according to the Chromium Next GEM Single Cell ATAC Solution User Guide (version CG000209 Rev F, 10×x Genomics) with the following modifications:

a. GEM Generation and Barcoding: During the GEM generation and barcoding reaction (step 2.1 in 10×x protocol), 1 µl of a 22.5 µM GoT-ChA primer mix was added to the barcoding mixture. The primers used in this step were: NUP98::NSD1 fusion forward: 5’ - TCGTCGGCA GCGTCAGATGTGTATAAGAGACAGAAGGCGTTTCTTCTCTGA* C*C*G. NUP98 WT forward: 5’ - TCGTCGGCAGCGTCAGATGTGTATAAGAGACAGGCATGTATGCAATGCCTT*G*G*T. Reverse: AACACTTACCGTATTCAC*C*C*C.
b. Post-GEM Incubation and Cleanup: During the post-GEM incubation clean-up (step 3.2 in 10×x protocol), 45.5 µl of Elution Solution I was used to elute material from SPRIselect beads (Beckman Coulter). A total of 5 µl was used for GoT-ChA library construction, and the remaining 40 µl was used for ATAC library construction as per the standard protocol.
c. GoT-ChA Library Construction: Two additional PCR reactions were performed using the 5 µl set aside during step 3.2 to generate the GoT-ChA library. The first PCR amplifies genotyping fragments before sample indexing. The primers used were: P5 (binds to the P5 Illumina sequencing handle): 5’-GATACGGCGACCACCGAGATCTACAC. GoT–ChA nested (biotinylated nested primer with a partial TruSeq small RNA read 2 handles): 5’-/5Bioag/ CCTTGGCACCCGAGAATTCCACCCCTTTGTTGAAATGAGTGGT. Thermocycling conditions are 95° C for 3 minutes, followed by 15 cycles of 95° C for 20 s, 65° C for 30 s, and 72° C for 20 s. The final extension is 72° C for 5 minutes, followed by holding at 4° C. After a 1.2 x SPRIselect cleanup, biotinylated PCR products were bound using Dynabeads M-280 Streptavidin magnetic beads (Thermo Fisher) at room temperature for 15 minutes. The beads were washed twice with 1×x SSPE buffer and once with 10 mM Tris-HCl (pH 8.0) before resuspending in water. The bead-bound fragments were amplified and indexed using the following primers: P5 (for Illumina sequencing handle), RPI-X (adds a sample index and P7 Illumina sequencing handle: CAAGCAGAAGACGGCATACGAGATXXXXXXXXGTGACTGGAGTTCCTTGGCACCCGAGAA TTCCA, where X represents a user-defined sample index). Thermocycling conditions are 95° C for 3 minutes, followed by 6-10 cycles of 95° C for 20 s, 65° C for 30 s, and 72° C for 20 s, final extension: 72° C for 5 minutes, followed by holding at 4° C. The final libraries were quantified using a High-Sensitivity DNA Chip (Agilent Technologies) run on a Bioanalyzer 2100 system (Agilent Technologies). Libraries were sequenced on the NovaSeq 6000 system (Illumina) with the following parameters: 50/8/16/50 for ATAC and 50/16/50 for the GoT-ChA library. The sequencing depth is 25,000 read pairs per nucleus for ATAC libraries and 5,000 read pairs per nucleus for GoT-ChA libraries.

### GoT-ChA data analysis

For GoT-ChA, the FASTQ files from chromatin accessibility data were processed to generate the cell-by-gene count matrix with CellRanger v.5.0.1 using default parameters, with GRCh38 as the human genome reference. The data were filtered to retain cells with at least 3,000 peaks detected, a minimum of 50% of reads in peaks, nucleosome signal below 4 and TSS enrichment score above 3. Dimensionality reduction was performed in Seurat (v.5.1.0) applying term frequency-inverse document frequency (TF-IDF) followed by single value decomposition for latent semantic indexing (LSI). For graphical representation, dimensions were further reduced via UMAP, removing the first LSI dimension which showed a high correlation (> 0.5) with sequencing depth. Cell types were assigned based on gene activity scores as calculated by the GeneActivity function from Seurat (v.5.1.0) and concordance with the labels assigned via bridge dictionary mapping.^72^ Transcription factor motif accessibility was calculated via chromVar.^23^ Genotyping FASTQ files were processed with the Gotcha R package (v.1.0), using wild-type and NUP98::NSD1 fusion sequences for perfect match. Genotype calls were included in Seurat object metadata for downstream analysis. Differential analysis was performed via differential linear mixture models using the DiffLMM function from Gotcha (v.1.0) as previously done.^19^

### *In vitro* drug studies

Primary FL-derived CD34+ cells or NUP98::NSD1/WT1ko leukemic cells were cultured and expanded in X-VIVO 10 medium supplied with growth factors as described before. 40,000 cells/wells were seeded in a 96-well flat bottom plate with 3 replicates for each condition. DMSO or venetoclax was diluted in the medium and added into the cells per the experimental design. At the designated time points, cells were harvested and stained with antibodies, Annexin V (Thermo Fisher), and 7-AAD Viability Staining Solution (Thermo Fisher) to assess percentage of immunophenotypic stem cells, myeloid differentiation, and apoptosis. Flow cytometry was performed using the Attune Flow Cytometer (Thermo Fisher), and the cell number in each well was recorded. All flow cytometry data were analyzed using FlowJo (FlowJo, LLC).

### *In vivo* xenograft drug studies

NUP98::NSD1 and NUP98::NSD1/WT1ko leukemic cells isolated from xenografts were transplanted into NSG mice via intra-femur injections (2-4×x10^5^ cells per mouse). After 4-6 weeks, mice were treated with cytarabine (Selleck Chemicals), revumenib (Syndax), venetoclax (Medchemexpress), or the combination. Cytarabine was given five consecutive days (50 mg/kg/day) by intraperitoneal injection, and mice were allowed to get 5 days rest. Revumenib chow food was given for 14 days (0.1% SNDX-5613 monocitrate diet). Venetoclax was given for 14 days by oral gavage (100 mg/kg/day). Mice were euthanized at weeks 6-8, and engraftment was considered positive if the human CD45+ cells exceeded 1% in the bone marrow and appeared as concentrated clusters rather than dispersed populations. In combination treatment conditions, revumenib or venetoclax were given 14 days, cytarabine was given five consecutive days (50 mg/kg/day) by intraperitoneal injection from day 4.

### MitoTracker assay

FL-derived CD34+ HSPCs, NUP98::NSD1 and NUP98::NSD1/WT1ko leukemic cells, were cultured and expanded in X-VIVO 10 medium. Cells were collected and spun down at 350 x g for 10 min at room temperature. Cells were resuspended gently in 37° C staining buffer (PBS with 2.5% FBS) containing the MitoTracker Green (Thermo Fisher, 25nM) and incubated at 37° C for 20 min. After staining is complete, cells were centrifuged and resuspended in a fresh prewarmed staining buffer. Cells were analyzed by flow cytometry. MitoTracker Green signals were normalized by cell size (forward side scatter).

### Seahorse assay

NUP98::NSD1, NUP98::NSD1/WT1ko, and NUP98::NSD1/WT1ko with PRDM16ko or control edited leukemic cells were cultured and expanded in X-VIVO 10 medium. On the experimental day, cells were resuspended in XF RPMI pH 7.4 assay medium (Agilent) supplemented with 1 mM pyruvate (Agilent), 10 mM glucose (Agilent), 2 mM L-glutamine (Agilent), and their respective treatment conditions. Cells were seeded at a density of 200,000 cells/well onto poly-D-lysine (ThermoFisher)-coated plates (Agilent) and centrifuged at 200 x g for 3 minutes. The plates were incubated in a non-CO2 incubator at 37° C for 15 minutes. For the pre-treatment experiment, cells were treated with venetoclax (Selleckchem) at different dosages (0, 20 nM, 200 nM, 2 nM) for 24 hours. Mito stress assays were processed using the manufacturer’s instructions to measure oxygen consumption rates (OCR) and extracellular acidification rates (ECAR). OCR was measured three times before and after the administration of each of the following: oligomycin (1.5 μM), carbonyl cyanide-4-(trifluoromethoxy) phenylhydrazone (FCCP, 1 μM), and a combination of rotenone and antimycin A (0.5 μM). The XFe96 analyzer was used. After the completion of the assay, normalization was performed by treating the cells with 0.1 % Triton and 0.125 μM YOYO-3 Iodide (Thermo Fisher) and incubating in the dark overnight at room temperature. Fluorescence was measured at 599/640 nm (excitation/emission) using the Synergy H1 Hybrid multimode microplate reader (BioTek). Fluorescence unit values were normalized by dividing by the 20% trimmed mean calculated from all wells.

### Mass spectrometry-based metabolomics

CD34+CD117+ LSC-enriched cells and non-CD34+CD117+ leukemic cells were sorted from xenografts of NUP98::NSD1 and NUP98::NSD1/WT1ko. Cells were washed with cold PBS three times. Cell pellets with 200,000 cells per sample (devoid of all supernatants) were kept at -80 °C in 1.5 mL tubes until submitted for mass-spectrometry-based metabolomics. For each condition, 3 biological replicates were used. Metabolites were extracted from frozen cell pellets at 4 x 10^6^ cells per mL by vigorous vortexing in the presence of ice-cold 5:3:2 MeOH:MeCN:ddH2O (v/v/v) for 30 minutes at 4° C. Supernatants were then clarified by centrifugation (10 min, 12,000 x g, 4° C). The resulting extracts were analyzed (10 µL per injection) by ultra-high-pressure liquid chromatography coupled to mass spectrometry (UHPLC-MS - Vanquish and Q Exactive, Thermo Fisher). Hydrophilic metabolites were resolved on a Kinetex C18 column using a 5 minute gradient exactly as previously described^73^ with the following parameters: spray voltage 4.0 kV, sheath gas 45, auxiliary gas 25, auxiliary gas heater 30° C, capillary temperature 320° C, S-lens RF level 50, AGC target 3e6, maximum injection time 200 ms, scan range 65-975 m/z, Orbitrap resolution (MS^1^) 70,000. Following data acquisition, .raw files were converted to .mzXML using RawConverter then metabolites assigned and peaks integrated at the MS^1^ level using El-Maven (Elucidata) in conjunction with the KEGG database and an in-house standard library. Quality control was assessed as using technical replicates run at beginning, end, and middle of each sequence as previously described.^74^

### Metabolomics analysis

For metabolomics analysis, data were log-transformed and autoscaled (mean-centered and divided by the standard deviation of each variable) prior to performing statistical analysis. Significant differences were determined by an unpaired t-test followed by the Benjamini-Hochberg post hoc test and defined FDR of 0.05. Pathway enrichment was performed using Metaboanalyst v6.0, with HMDB ID as a compound identifier and the KEGG pathway database as a source.^75^

### ChIP-seq library preparation

Approximately 1×x10^7^ NUP98::NSD1/WT1ko leukemic cells were crosslinked and processed for ChIP according to this study^76^. Briefly, cells were harvested in a 15 mL Falcon tube, washed three times with PBS, and cross-linked with 10 mL of freshly prepared 1% Formaldehyde/PBS for 10 min. The reaction was quenched by adding 2.5 M Glycine to a final concentration of 0.125 M, followed by a 5 min incubation at room temperature. Then cells were centrifuged at 1000g for 10 min at 4° C, and the supernatant was discarded. Nuclei were then isolated, and chromatin was fragmented by sonication using a Bioruptor (Diagenode) (high intensity, 24 cycles, 30 seconds ON/30 seconds OFF). A 50 µL chromatin aliquot was saved as an input control, while immunoprecipitation was performed using PRDM16 (R&D, 5 μg per reaction) and H3K27ac (Cell Signaling, 5 μg per reaction) antibodies. The resulting DNA was purified with SPRIselect beads (Beckman Coulter), and quality control was assessed using a TapeStation (Agilent). Finally, DNA libraries were generated following the NEBNext Ultra II DNA Library Prep Kit protocol. TapeStation was used to evaluate the library’s quality and sequencing was conducted on a NextSeq 500 in 75 bp single-end mode.

### ChIP-seq analysis

Reads were first evaluated for their quality using FastQC (v0.11.8, RRID: SCR_014583). Reads were trimmed for adaptor sequences using Trim Galore! (v0.6.6, RRID:SCR_011847) and aligned to the GRCh38/hg38 using Bowtie2 (version 2.1.8)^77^. Picard (v2.2.4, RRID: SCR_006525) was used to remove duplicated reads. Binary alignment map (BAM) files were generated with samtools v1.11^78^, and were used in downstream analysis. Significant peaks were identified with MACS2 (version 2.1.0) using input as control (q-value<0.01). Peaks in ENCODE blacklisted regions were removed. The called peaks were annotated to the closest gene using ChIPSeeker (v1.30.3) with annotatePeak function and the feature distribution was visualized with the plotAnnoBar function. Overrepresentation analysis (ORA) was performed with the R package clusterProfiler v4.2.2^79^ using the enricher function. Significant pathways were called with an adjusted p value < 0.05 using the Benjamini and Hochberg procedure. Transcription factor motif enrichment analysis was performed on the called peak set. De novo motifs were identified within a ±200bp window around ChIP peak summit using HOMERv4.11 findMotifsGenome.pl function with parameters hg38 -size 200, with the default background regions generated by HOMER^80^. Coverage tracks were generated from Bam files using deepTools (v3.2.1, RRID: SCR_016366) bamCoverage with parameters –normalizeUsingRPKM --binsize 10 --extendReads=200.^81^ Bigwig files were uploaded to the UCSC genome browser for visualization. Heatmaps of genomic regions were generated with deepTools. The command computeMatrix with reference-point mode was used to calculate scores at genomic regions and generate a matrix file. To visualize the signals at binding sites, plotHeatmap was used to generate heatmaps with profile plots. To identify the genes related to Fatty Acid Metabolism, the PRDM16 peak associated genes were overlapped with the hallmark geneset and 25 genes were identified in PRDM16 binding sites: ACADM, EPHX1, LDHA, ACSM3, ACAA2, IL4I1, MDH1, GPD2, ACAA1, CPOX, MGLL, RAP1GDS1, METAP1, ACSL1, BPHL, ELOVL5, BCKDHB, ME1, SERINC1, MDH2, CD36, ALDH1A1, SMS, HSD17B10, XIST.

### Bulk RNA-seq of clinical cohort

Pediatric patients with de novo AML enrolled in COG trials CCG-2961, AAML03P1, AAML0531, and AAML1031, were included for bulk RNA-seq when biological samples were available. Total RNA from diagnostic peripheral blood or bone marrow was extracted and purified using the QIAcube automated system with AllPrep DNA/RNA/miRNA Universal Kits (Qiagen). Libraries were prepared for 75-bp strand-specific paired-end sequencing using the ribodepletion v2.0 protocol by the British Columbia Genome Sciences Center (BCGSC). Libraries were sequenced on the Illumina HiSeq 2000/2500 and aligned to the hg19 (GRCh37-lite) reference genome using BWA v0.5.7 with default parameters, except the addition of “-s” option, and duplicate reads were marked with Picard Tools.

### Fusion detection of clinical cohort

The NUP98 fusions were detected by either karyotype or combined fusion detection algorithms STAR-fusion v1.8.1, TransAbyss v1.4.10, and CICERO v0.1.8 completed on RNA-seq. STAR-fusion was run using default parameters with a premade GRCh37 resource library with Gencode v19 annotations. The TransAbyss software was executed with the GRCh37-lite reference genome, and the following parameters included: fusion breakpoint reads ≥1, flanking pairs, and spanning reads ≥2 counts. CICERO fusion detection was performed with default parameters with GRCh37-lite. Fusions detected computationally were verified using Fusion Inspector v.1.8.1 (Broad Institute, Cambridge) and visualized on IGV and BAMBINO.

### Gene expression analysis of clinical cohort

Resultant gene expression was calculated in transcripts per million (TPM). NUP98::NSD1 samples had a higher expression of PRDM16 (median 12.9 TPM, range 0.1-55.8 TPM) than NUP98::KDM5A (median 4.6 TPM, range 0.0-15.2 TPM). Other NUP98 fusion partners had the lowest expression of PRDM16 (median 0.0 TPM, range 0.0-1.3 TPM).

### Survival analyses of clinical cohort

Fetal-oncogenic gene signature included the top 100 higher expressed genes (based on LogFC and adjusted p-value) between FL-versus CB-derived LMPP/Early GMPs. Log2 fold-change was multiplied by -log10 times the adjusted p-value to generate a rank score for each gene. These ranks were then multiplied by the transcript expression (transcripts per million) for the corresponding gene for each patient. These values were then summed, creating an enrichment score for each patient. A positive enrichment score indicated a transcriptome that was more fetal-like, whereas a negative enrichment score was more CB-like. 5-year event-free (EFS) and overall survival (OS) outcomes were analyzed using the Kaplan-Meier method. Significance was measured using log-rank p-values.

### Statistical analysis

Statistical analysis was performed using GraphPad Prism software Version 10.2.3. Statistical significance was calculated using a two-tailed unpaired t-test, two way-ANOVA test and Pearson’s chi-squared test. Event-free (EFS) and overall survival (OS) outcomes were analyzed using the Kaplan-Meier method, significance was measured using log-rank test. Other relevant statistical methods are detailed in the methods section and figure legends. For all analyses, p < 0.05 was considered statistically significant.

## References

1. Kentsis, A. (2024). Toward a Unified Theory of Why Young People Develop Cancer. Cold Spring Harb. Perspect. Med. 14, a041658. 10.1101/cshperspect.a041658.

2. Chapman, M.S., Mitchell, E., Yoshida, K., Williams, N., Fabre, M.A., Ranzoni, A.M., Robinson, P.S., Kregar, L.D., Wilk, M., Boettcher, S., et al. (2025). Prolonged persistence of mutagenic DNA lesions in somatic cells. Nature, 1–10. 10.1038/s41586-024-08423-8.

3. Greaves, M. (2005). In utero origins of childhood leukaemia. Early Hum. Dev. 81, 123–129. 10.1016/j.earlhumdev.2004.10.004.

4. Körber, V., Stainczyk, S.A., Kurilov, R., Henrich, K.-O., Hero, B., Brors, B., Westermann, F., and Höfer, T. (2023). Neuroblastoma arises in early fetal development and its evolutionary duration predicts outcome. Nat. Genet. 55, 619–630. 10.1038/s41588-023-01332-y.

5. Wagenblast, E., Araújo, J., Gan, O.I., Cutting, S.K., Murison, A., Krivdova, G., Azkanaz, M., McLeod, J.L., Smith, S.A., Gratton, B.A., et al. (2021). Mapping the cellular origin and early evolution of leukemia in Down syndrome. Science 373. 10.1126/science.abf6202.

6. Society, A.C. Cancer Facts & Figures 2025.

7. Bolouri, H., Farrar, J.E., Triche, T., Ries, R.E., Lim, E.L., Alonzo, T.A., Ma, Y., Moore, R., Mungall, A.J., Marra, M.A., et al. (2018). The molecular landscape of pediatric acute myeloid leukemia reveals recurrent structural alterations and age-specific mutational interactions. Nat. Med. 24, 103–112. 10.1038/nm.4439.

8. Umeda, M., Ma, J., Westover, T., Ni, Y., Song, G., Maciaszek, J.L., Rusch, M., Rahbarinia, D., Foy, S., Huang, B.J., et al. (2024). A new genomic framework to categorize pediatric acute myeloid leukemia. Nat. Genet. 56, 281–293. 10.1038/s41588-023-01640-3.

9. Cooper, T.M., Alonzo, T.A., Tasian, S.K., Kutny, M.A., Hitzler, J., Pollard, J.A., Aplenc, R., Meshinchi, S., and Kolb, E.A. (2023). Children’s Oncology Group’s 2023 blueprint for research: Myeloid neoplasms. Pediatr. Blood Cancer 70, e30584. 10.1002/pbc.30584.

10. Mercher, T., and Schwaller, J. (2019). Pediatric Acute Myeloid Leukemia (AML): From Genes to Models Toward Targeted Therapeutic Intervention. Front Pediatr 7, 401. 10.3389/fped.2019.00401.

11. Wang, G.G., Cai, L., Pasillas, M.P., and Kamps, M.P. (2007). NUP98-NSD1 links H3K*36* methylation to Hox-A gene activation and leukaemogenesis. Nat Cell Biol 9, 804–812. 10.1038/ncb1608.

12. Thanasopoulou, A., Tzankov, A., and Schwaller, J. (2014). Potent co-operation between the NUP98-NSD1 fusion and the FLT3-ITD mutation in acute myeloid leukemia induction. Haematologica 99, 1465–1471. 10.3324/haematol.2013.100917.

13. Bertrums, E.J.M., Smith, J.L., Harmon, L., Ries, R.E., Wang, Y.-C.J., Alonzo, T.A., Menssen, A.J., Chisholm, K.M., Leonti, A.R., Tarlock, K., et al. (2023). Comprehensive molecular and clinical characterization of NUP98 fusions in pediatric acute myeloid leukemia. Haematologica 108, 2044–2058. 10.3324/haematol.2022.281653.

14. Hollink, I.H.I.M., Heuvel-Eibrink, M.M. van den, Arentsen-Peters, S.T.C.J.M., Pratcorona, M., Abbas, S., Kuipers, J.E., Galen, J.F. van, Beverloo, H.B., Sonneveld, E., Kaspers, G.-J.J.L., et al. (2011). NUP98/NSD1 characterizes a novel poor prognostic group in acute myeloid leukemia with a distinct HOX gene expression pattern. Blood 118, 3645–3656. 10.1182/blood-2011-04-346643.

15. Wang, Y., Xiao, M., Chen, X., Chen, L., Xu, Y., Lv, L., Wang, P., Yang, H., Ma, S., Lin, H., et al. (2015). WT1 recruits TET2 to regulate its target gene expression and suppress leukemia cell proliferation. Mol. Cell 57, 662–673. 10.1016/j.molcel.2014.12.023.

16. Umeda, M., Ma, J., Huang, B.J., Hagiwara, K., Westover, T., Abdelhamed, S., Barajas, J.M., Thomas, M.E., Walsh, M.P., Song, G., et al. (2022). Integrated genomic analysis identifies UBTF tandem duplications as a recurrent lesion in pediatric acute myeloid leukemiaUBTF tandem duplications in pediatric acute myeloid leukemia. Blood Cancer Discov. 3, 194–207. 10.1158/2643-3230.bcd-21-0160.

17. McNeer, N.A., Philip, J., Geiger, H., Ries, R.E., Lavallée, V.-P., Walsh, M., Shah, M., Arora, K., Emde, A.-K., Robine, N., et al. (2019). Genetic mechanisms of primary chemotherapy resistance in pediatric acute myeloid leukemia. Leukemia 33, 1934–1943. 10.1038/s41375-019-0402-3.

18. Niktoreh, N., Walter, C., Zimmermann, M., Neuhoff, C. von, Neuhoff, N. von, Rasche, M., Waack, K., Creutzig, U., Hanenberg, H., and Reinhardt, D. (2019). Mutated WT1, FLT3-ITD, and NUP98-NSD1 Fusion in Various Combinations Define a Poor Prognostic Group in Pediatric Acute Myeloid Leukemia. J Oncol 2019, 1609128. 10.1155/2019/1609128.

19. Izzo, F., Myers, R.M., Ganesan, S., Mekerishvili, L., Kottapalli, S., Prieto, T., Eton, E.O., Botella, T., Dunbar, A.J., Bowman, R.L., et al. (2024). Mapping genotypes to chromatin accessibility profiles in single cells. Nature, 1–9. 10.1038/s41586-024-07388-y.

20. Chandra, B., Michmerhuizen, N.L., Shirnekhi, H.K., Tripathi, S., Pioso, B.J., Baggett, D.W., Mitrea, D.M., Iacobucci, I., White, M.R., Chen, J., et al. (2021). Phase Separation Mediates NUP98 Fusion Oncoprotein Leukemic TransformationPhase Separation Drives Oncogenesis by NUP98 Fusion Proteins. Cancer Discov 12, 1152–1169. 10.1158/2159-8290.cd-21-0674.

21. Terlecki-Zaniewicz, S., Humer, T., Eder, T., Schmoellerl, J., Heyes, E., Manhart, G., Kuchynka, N., Parapatics, K., Liberante, F.G., Müller, A.C., et al. (2021). Biomolecular condensation of NUP98 fusion proteins drives leukemogenic gene expression. Nat. Struct. Mol. Biol. 28, 190–201. 10.1038/s41594-020-00550-w.

22. Zeng, A.G.X., Iacobucci, I., Shah, S., Mitchell, A., Wong, G., Bansal, S., Chen, D., Gao, Q., Kim, H., Kennedy, J.A., et al. (2024). Single-cell transcriptional mapping reveals genetic and non-genetic determinants of aberrant differentiation in AML. bioRxiv, 2023.12.26.573390. 10.1101/2023.12.26.573390.

23. Schep, A.N., Wu, B., Buenrostro, J.D., and Greenleaf, W.J. (2017). chromVAR: inferring transcription-factor-associated accessibility from single-cell epigenomic data. Nat. Methods 14, 975–978. 10.1038/nmeth.4401.

24. Fernández-Maestre, I., Cai, S.F., and Levine, R.L. (2024). A View of Myeloid Transformation through the Hallmarks of Cancer. Blood Cancer Discov. 5, 377–387. 10.1158/2643-3230.bcd-24-0009.

25. Zeng, A.G.X., Bansal, S., Jin, L., Mitchell, A., Chen, W.C., Abbas, H.A., Chan-Seng-Yue, M., Voisin, V., Galen, P. van, Tierens, A., et al. (2022). A cellular hierarchy framework for understanding heterogeneity and predicting drug response in acute myeloid leukemia. Nat Med 28, 1212–1223. 10.1038/s41591-022-01819-x.

26. Manno, G.L., Soldatov, R., Zeisel, A., Braun, E., Hochgerner, H., Petukhov, V., Lidschreiber, K., Kastriti, M.E., Lönnerberg, P., Furlan, A., et al. (2018). RNA velocity of single cells. Nature 560, 494–498. 10.1038/s41586-018-0414-6.

27. Haghverdi, L., Büttner, M., Wolf, F.A., Buettner, F., and Theis, F.J. (2016). Diffusion pseudotime robustly reconstructs lineage branching. Nat. Methods 13, 845–848. 10.1038/nmeth.3971.

28. Potluri, S., Assi, S.A., Chin, P.S., Coleman, D.J.L., Pickin, A., Moriya, S., Seki, N., Heidenreich, O., Cockerill, P.N., and Bonifer, C. (2021). Isoform-specific and signaling-dependent propagation of acute myeloid leukemia by Wilms tumor 1. Cell Rep. 35, 109010. 10.1016/j.celrep.2021.109010.

29. Udtha, M., Lee, S.-J., Alam, R., Coombes, K., and Huff, V. (2003). Upregulation of c-MYC in WT1-mutant tumors: assessment of WT1 putative transcriptional targets using cDNA microarray expression profiling of genetically defined Wilms’ tumors. Oncogene 22, 3821–3826. 10.1038/sj.onc.1206597.

30. Zhang, X., Xing, G., and Saunders, G.F. (1999). Proto-oncogene N-myc promoter is down regulated by the Wilms’ tumor suppressor gene WT1. Anticancer Res. 19, 1641–1648.

31. Ullmark, T., Järvstråt, L., Sandén, C., Montano, G., Jernmark-Nilsson, H., Lilljebjörn, H., Lennartsson, A., Fioretos, T., Drott, K., Vidovic, K., et al. (2017). Distinct global binding patterns of the Wilms tumor gene 1 (WT1) −KTS and +KTS isoforms in leukemic cells. Haematologica 102, 336–345. 10.3324/haematol.2016.149815.

32. Pan, X.-N., Chen, J.-J., Wang, L.-X., Xiao, R.-Z., Liu, L.-L., Fang, Z.-G., Liu, Q., Long, Z.-J., and Lin, D.-J. (2014). Inhibition of c-Myc Overcomes Cytotoxic Drug Resistance in Acute Myeloid Leukemia Cells by Promoting Differentiation. PLoS ONE 9, e105381. 10.1371/journal.pone.0105381.

33. Farge, T., Saland, E., Toni, F. de, Aroua, N., Hosseini, M., Perry, R., Bosc, C., Sugita, M., Stuani, L., Fraisse, M., et al. (2017). Chemotherapy-Resistant Human Acute Myeloid Leukemia Cells Are Not Enriched for Leukemic Stem Cells but Require Oxidative Metabolism. Cancer Discov. 7, 716–735. 10.1158/2159-8290.cd-16-0441.

34. Reddy, J.C., Hosono, S., and Licht, J.D. (1995). The Transcriptional Effect of WT1 Is Modulated by Choice of Expression Vector (∗). J. Biol. Chem. 270, 29976–29982. 10.1074/jbc.270.50.29976.

35. Ritchie, M.F., Zhou, Y., and Soboloff, J. (2011). WT1/EGR1-mediated control of STIM1 expression and function in cancer cells. Front. Biosci. 16, 2402. 10.2741/3862.

36. Min, I.M., Pietramaggiori, G., Kim, F.S., Passegué, E., Stevenson, K.E., and Wagers, A.J. (2008). The Transcription Factor EGR1 Controls Both the Proliferation and Localization of Hematopoietic Stem Cells. Cell Stem Cell 2, 380–391. 10.1016/j.stem.2008.01.015.

37. Desterke, C., Bennaceur-Griscelli, A., and Turhan, A.G. (2021). EGR1 dysregulation defines an inflammatory and leukemic program in cell trajectory of human-aged hematopoietic stem cells (HSC). Stem Cell Res. Ther. 12, 419. 10.1186/s13287-021-02498-0.

38. Thomas, X. (2024). Small Molecule Menin Inhibitors: Novel Therapeutic Agents Targeting Acute Myeloid Leukemia with KMT2A Rearrangement or NPM1 Mutation. Oncol. Ther. 12, 57–72. 10.1007/s40487-024-00262-x.

39. Heikamp, E.B., Henrich, J.A., Perner, F., Wong, E.M., Hatton, C., Wen, Y., Barwe, S.P., Gopalakrishnapillai, A., Xu, H., Uckelmann, H.J., et al. The Menin-MLL1 interaction is a molecular dependency in NUP98-rearranged AML. Blood 139, 894–906. 10.1182/blood.2021012806.

40. Perner, F., Stein, E.M., Wenge, D.V., Singh, S., Kim, J., Apazidis, A., Rahnamoun, H., Anand, D., Marinaccio, C., Hatton, C., et al. (2023). MEN1 mutations mediate clinical resistance to menin inhibition. Nature 615, 913–919. 10.1038/s41586-023-05755-9.

41. Duffy, M.J., O’Grady, S., Tang, M., and Crown, J. (2021). MYC as a target for cancer treatment. Cancer Treat. Rev. 94, 102154. 10.1016/j.ctrv.2021.102154.

42. Mishra, S.K., Millman, S.E., and Zhang, L. (2023). Metabolism in acute myeloid leukemia: mechanistic insights and therapeutic targets. Blood 141, 1119–1135. 10.1182/blood.2022018092.

43. Rondeau, V., Berman, J.M., Ling, T., O’Brien, C., Culp-Hill, R., Reisz, J.A., Wunderlich, M., Chueh, Y., Jiménez-Camacho, K.E., Sexton, C., et al. (2024). Spermidine metabolism regulates leukemia stem and progenitor cell function through KAT7 expression in patient-derived mouse models. Sci. Transl. Med. 16, eadn1285. 10.1126/scitranslmed.adn1285.

44. Aibar, S., González-Blas, C.B., Moerman, T., Huynh-Thu, V.A., Imrichova, H., Hulselmans, G., Rambow, F., Marine, J.-C., Geurts, P., Aerts, J., et al. (2017). SCENIC: single-cell regulatory network inference and clustering. Nat. Methods 14, 1083–1086. 10.1038/nmeth.4463.

45. Yamato, G., Yamaguchi, H., Handa, H., Shiba, N., Kawamura, M., Wakita, S., Inokuchi, K., Hara, Y., Ohki, K., Okubo, J., et al. (2017). Clinical features and prognostic impact of PRDM16 expression in adult acute myeloid leukemia. Genes, Chromosom. Cancer 56, 800–809. 10.1002/gcc.22483.

46. Dao, F.-T., Chen, W.-M., Long, L.-Y., Li, L.-D., Yang, L., Wang, J., Liu, Y.-R., Jiang, H., Zhang, X.-H., Jiang, Q., et al. (2021). High PRDM16 expression predicts poor outcomes in adult acute myeloid leukemia patients with intermediate cytogenetic risk: a comprehensive cohort study from a single Chinese center. Leuk. Lymphoma 62, 185–193. 10.1080/10428194.2020.1817436.

47. Yamato, G., Kawai, T., Shiba, N., Ikeda, J., Hara, Y., Ohki, K., Tsujimoto, S.-I., Kaburagi, T., Yoshida, K., Shiraishi, Y., et al. (2022). Genome-wide DNA Methylation Analysis in Pediatric Acute Myeloid Leukemia. Blood Adv. 6, 3207–3219. 10.1182/bloodadvances.2021005381.

48. Gudmundsson, K.O., Nguyen, N., Oakley, K., Han, Y., Gudmundsdottir, B., Liu, P., Tessarollo, L., Jenkins, N.A., Copeland, N.G., and Du, Y. (2020). Prdm16 is a critical regulator of adult long-term hematopoietic stem cell quiescence. Proc. Natl. Acad. Sci. 117, 31945– 31953. 10.1073/pnas.2017626117.

49. Wang, Q., Li, H., Tajima, K., Verkerke, A.R.P., Taxin, Z.H., Hou, Z., Cole, J.B., Li, F., Wong, J., Abe, I., et al. (2022). Post-translational control of beige fat biogenesis by PRDM16 stabilization. Nature 609, 151–158. 10.1038/s41586-022-05067-4.

50. Wang, W., Ishibashi, J., Trefely, S., Shao, M., Cowan, A.J., Sakers, A., Lim, H.-W., O’Connor, S., Doan, M.T., Cohen, P., et al. (2019). A PRDM16-Driven Metabolic Signal from Adipocytes Regulates Precursor Cell Fate. Cell Metab. 30, 174–189.e5. 10.1016/j.cmet.2019.05.005.

51. Zhang, Y., Xie, X., Huang, Y., Liu, M., Li, Q., Luo, J., He, Y., Yin, X., Ma, S., Cao, W., et al. (2022). Temporal molecular program of human hematopoietic stem and progenitor cells after birth. Dev. Cell 57, 2745–2760.e6. 10.1016/j.devcel.2022.11.013.

52. Suresh, V., Bhattacharya, B., Tshuva, R.Y., Gotthold, M.D., Olender, T., Bose, M., Pradhan, S.J., Zeev, B.B., Smith, R.S., Tole, S., et al. (2024). PRDM16 co-operates with LHX2 to shape the human brain. Oxf. Open Neurosci. 3, kvae001. 10.1093/oons/kvae001.

53. Popescu, B., Stahlhut, C., Tarver, T.C., Wishner, S., Lee, B.J., Peretz, C.A.C., Luck, C., Phojanakong, P., Serrano, J.A.C., Hongo, H., et al. (2023). Allosteric SHP2 inhibition increases apoptotic dependency on BCL2 and synergizes with venetoclax in FLT3- and KIT-mutant AML. Cell Rep. Med. 4, 101290. 10.1016/j.xcrm.2023.101290.

54. Kasbekar, M., Mitchell, C.A., Proven, M.A., and Passegué, E. (2023). Hematopoietic stem cells through the ages: A lifetime of adaptation to organismal demands. Cell Stem Cell 30, 1403–1420. 10.1016/j.stem.2023.09.013.

55. Li, H., Côté, P., Kuoch, M., Ezike, J., Frenis, K., Afanassiev, A., Greenstreet, L., Tanaka-Yano, M., Tarantino, G., Zhang, S., et al. (2024). The dynamics of hematopoiesis over the human lifespan. Nat. Methods, 1–13. 10.1038/s41592-024-02495-0.

56. Xiang, X., Lu, Q., Xu, X., Cai, P., Chen, S., Pan, J., and Zeng, Z. (2022). Prognostic impact of PRDM16 expression in acute myeloid leukemia with normal cytogenetics. Hematology 27, 499–505. 10.1080/16078454.2022.2066306.

57. Bateman, C.M., Colman, S.M., Chaplin, T., Young, B.D., Eden, T.O., Bhakta, M., Gratias, E.J., Wering, E.R. van, Cazzaniga, G., Harrison, C.J., et al. (2010). Acquisition of genome-wide copy number alterations in monozygotic twins with acute lymphoblastic leukemia. Blood 115, 3553–3558. 10.1182/blood-2009-10-251413.

58. Greaves, M.F., Maia, A.T., Wiemels, J.L., and Ford, A.M. (2003). Leukemia in twins: lessons in natural history. Blood 102, 2321–2333. 10.1182/blood-2002-12-3817.

59. Sousos, N., Leathlobhair, M.N., Karali, C.S., Louka, E., Bienz, N., Royston, D., Clark, S.-A., Hamblin, A., Howard, K., Mathews, V., et al. (2022). In utero origin of myelofibrosis presenting in adult monozygotic twins. Nat Med 28, 1207–1211. 10.1038/s41591-022-01793-4.

60. Visvanathan, A., Saulnier, O., Chen, C., Haldipur, P., Orisme, W., Delaidelli, A., Shin, S., Millman, J., Bryant, A., Abeysundara, N., et al. (2024). Early rhombic lip Protogenin+ve stem cells in a human-specific neurovascular niche initiate and maintain group 3 medulloblastoma. Cell 187, 4733–4750.e26. 10.1016/j.cell.2024.06.011.

61. Wagenblast, E., Azkanaz, M., Smith, S.A., Shakib, L., McLeod, J.L., Krivdova, G., Araújo, J., Shultz, L.D., Gan, O.I., Dick, J.E., et al. (2019). Functional profiling of single CRISPR/Cas9-edited human long-term hematopoietic stem cells. Nature Communications 10, 4730–11. 10.1038/s41467-019-12726-0.

62. Erard, N., Knott, S.R.V., and Hannon, G.J. (2017). A CRISPR Resource for Individual, Combinatorial, or Multiplexed Gene Knockout. Mol. Cell 67, 348–354.e4. 10.1016/j.molcel.2017.06.030.

63. Brinkman, E.K., Chen, T., Amendola, M., and Steensel, B. van (2014). Easy quantitative assessment of genome editing by sequence trace decomposition. Nucleic Acids Res. 42, e168– e168. 10.1093/nar/gku936.

64. Butler, A., Hoffman, P., Smibert, P., Papalexi, E., and Satija, R. (2018). Integrating single-cell transcriptomic data across different conditions, technologies, and species. Nat. Biotechnol. 36, 411–420. 10.1038/nbt.4096.

65. Hao, Y., Hao, S., Andersen-Nissen, E., Mauck, W.M., Zheng, S., Butler, A., Lee, M.J., Wilk, A.J., Darby, C., Zager, M., et al. (2021). Integrated analysis of multimodal single-cell data. Cell 184, 3573–3587.e29. 10.1016/j.cell.2021.04.048.

66. Dann, E., Henderson, N.C., Teichmann, S.A., Morgan, M.D., and Marioni, J.C. (2022). Differential abundance testing on single-cell data using k-nearest neighbor graphs. Nat. Biotechnol. 40, 245–253. 10.1038/s41587-021-01033-z.

67. Bergen, V., Lange, M., Peidli, S., Wolf, F.A., and Theis, F.J. (2020). Generalizing RNA velocity to transient cell states through dynamical modeling. Nat. Biotechnol. 38, 1408–1414. 10.1038/s41587-020-0591-3.

68. Virshup, I., Rybakov, S., Theis, F.J., Angerer, P., and Wolf, F.A. (2021). anndata: Annotated data. bioRxiv, 2021.12.16.473007. 10.1101/2021.12.16.473007.

69. Wolf, F.A., Angerer, P., and Theis, F.J. (2018). SCANPY: large-scale single-cell gene expression data analysis. Genome Biol. 19, 15. 10.1186/s13059-017-1382-0.

70. Stuart, T., Srivastava, A., Madad, S., Lareau, C.A., and Satija, R. (2021). Single-cell chromatin state analysis with Signac. Nat. Methods 18, 1333–1341. 10.1038/s41592-021-01282-5.

71. Baranasic, D., Hörtenhuber, M., Balwierz, P.J., Zehnder, T., Mukarram, A.K., Nepal, C., Várnai, C., Hadzhiev, Y., Jimenez-Gonzalez, A., Li, N., et al. (2022). Multiomic atlas with functional stratification and developmental dynamics of zebrafish cis-regulatory elements. Nat. Genet. 54, 1037–1050. 10.1038/s41588-022-01089-w.

72. Hao, Y., Stuart, T., Kowalski, M.H., Choudhary, S., Hoffman, P., Hartman, A., Srivastava, A., Molla, G., Madad, S., Fernandez-Granda, C., et al. (2024). Dictionary learning for integrative, multimodal and scalable single-cell analysis. Nat. Biotechnol. 42, 293–304. 10.1038/s41587-023-01767-y.

73. Nemkov, T., Reisz, J.A., Gehrke, S., Hansen, K.C., and D’Alessandro, A. (2019). High-Throughput Metabolomics, Methods and Protocols. Methods Mol. Biol. 1978, 13–26. 10.1007/978-1-4939-9236-2_2.

74. Nemkov, T., Hansen, K.C., and D’Alessandro, A. (2017). A three-minute method for high-throughput quantitative metabolomics and quantitative tracing experiments of central carbon and nitrogen pathways. Rapid Commun. Mass Spectrom. 31, 663–673. 10.1002/rcm.7834.

75. Pang, Z., Xu, L., Viau, C., Lu, Y., Salavati, R., Basu, N., and Xia, J. (2024). MetaboAnalystR 4.0: a unified LC-MS workflow for global metabolomics. Nat. Commun. 15, 3675. 10.1038/s41467-024-48009-6.

76. Tian, B., Yang, J., and Brasier, A.R. (2011). Transcriptional Regulation, Methods and Protocols. Methods Mol. Biol. 809, 105–120. 10.1007/978-1-61779-376-9_7.

77. Langmead, B., Trapnell, C., Pop, M., and Salzberg, S.L. (2009). Ultrafast and memory-efficient alignment of short DNA sequences to the human genome. Genome Biol. 10, R25. 10.1186/gb-2009-10-3-r25.

78. Li, H., Handsaker, B., Wysoker, A., Fennell, T., Ruan, J., Homer, N., Marth, G., Abecasis, G., Durbin, R., and Subgroup, 1000 Genome Project Data Processing (2009). The Sequence Alignment/Map format and SAMtools. Bioinformatics 25, 2078–2079. 10.1093/bioinformatics/btp352.

79. Yu, G., Wang, L.-G., Han, Y., and He, Q.-Y. (2012). clusterProfiler: an R Package for Comparing Biological Themes Among Gene Clusters. OMICS: A J. Integr. Biol. 16, 284–287. 10.1089/omi.2011.0118.

80. Heinz, S., Benner, C., Spann, N., Bertolino, E., Lin, Y.C., Laslo, P., Cheng, J.X., Murre, C., Singh, H., and Glass, C.K. (2010). Simple combinations of lineage-determining transcription factors prime cis-regulatory elements required for macrophage and B cell identities. Mol. Cell 38, 576–589. 10.1016/j.molcel.2010.05.004.

81. Ramírez, F., Ryan, D.P., Grüning, B., Bhardwaj, V., Kilpert, F., Richter, A.S., Heyne, S., Dündar, F., and Manke, T. (2016). deepTools2: a next generation web server for deep-sequencing data analysis. Nucleic Acids Res. 44, W160–W165. 10.1093/nar/gkw257.

